# An original structural fold underlies the multitask P1, a silencing suppressor encoded by the Rice yellow mottle virus

**DOI:** 10.1101/2020.02.24.963488

**Authors:** Vianney Poignavent, François Hoh, Guillaume Terral, Yang Yinshan, François-Xavier Gillet, Jeong-Hyeon Kim, Frédéric Allemand, Eric Lacombe, Christophe Brugidou, Sarah Cianferani, Hélène Déméné, Florence Vignols

**Author notes:** These authors contributed equally to the work (François Hoh, Guillaume Terral).

## Abstract

The Rice Yellow Mottle sobemovirus (RYMV) belongs to the most damaging pathogens devastating rice fields in Africa. P1, a key protein for RYMV, was reported as a potent RNAi suppressor counteracting RNA silencing in plant reporter systems. Here we describe the complete 3D structure and dynamics of P1. Its N-terminal region contains ZnF1, a structural CCCC-type zinc finger strongly affine to zinc and a prominent short helix, rendering this region poorly amenable to structural changes. P1 C-terminal region contains ZnF2, an atypical HCHC-type ZnF that does not belong to any existing class of Zn finger proteins. ZnF2 appeared much less affine to zinc and more sensitive to oxidizing environments than ZnF1, and may serve as a sensor of plant redox status. The structure helped us to identify key residues essential for RYMV infectivity and spread in rice tissues through their participation in P1 oligomerization and folding. Altogether, our results provide the first complete structure of an antiviral silencing suppressor encoded by a virus infecting rice and highlight P1 structural and dynamical properties that may serve RYMV functions to infect and invade its host plant.

## INTRODUCTION

Despite their rudimentary genome that encodes little proteins to assume their host life cycle, viruses have to rapidly establish complex interactions with host cellular machinery to guarantee successful infection. RNA viruses usually present smaller genomes compared to DNA viruses, their genomes being under pressure for genome compression due to high mutation rate during viral replication (Holmes, 2003; Belshaw *et al*, 2007). At the same time, keeping a small size genome ensures the formation of smaller virions, which requires a minimal number of coat proteins (CP) for virion formation, thus maximizing virus production in host cells (Ben-Shaul & Gelbart, 2015). RNA viruses have evolved different mechanisms to multiply viral functions while maintaining the size of their genome, which is considered as creation of genomic novelty from a single nucleic acids sequence. At the RNA level, splicing, leaky scanning, overlapping open reading frames (ORF), sub-genomic RNA and ribosomal frame shift are major pathways allowing production of different proteins from only a single sequence (Löwer *et al*, 1995; Brierley, 1995; Xu *et al*, 2001; Baril & Brakier-Gingras, 2005; Dreher & Miller, 2006; Belshaw *et al*, 2007; Sztuba-Solinska *et al*, 2011; Firth & Brierley, 2012). On the other hand, viruses tend to use one protein for different functions, thus multiplying viral functions at their genome scale, while keeping the number of proteins constant. Because of cumulative functions, those so-called multifunctional proteins often present conformational flexibility and intrinsic disorders to ensure their distinct functions (Rantalainen *et al*, 2011; Plisson *et al*, 2003; Xue *et al*, 2010) and/or present multi-domains structure, each domain carrying a dedicated function (Deng *et al*, 2015; Sorel *et al*, 2014; Valli *et al*, 2018).

Plant viruses replicate in prime infected area, and rapidly spread in surrounding cells until they reach vascular tissues to establish systemic infection. Replication of RNA viruses is ensured by their viral RNA dependent RNA polymerase (RdRp), which often occurs in virus-modified membrane structures (Heinlein, 2015). Viral progeny moves intra-cellularly and cell-to-cell through plasmodesmata by high-jacking cellular machinery with viral encoded movement protein (MP) (Carrington *et al*, 1996; Waigmann *et al*, 2004; Heinlein, 2015). No extensive similarity of MP sequences or molecular mechanism involved in virus transport has been found, and each virus shows MP’s singularity and mode of action (Taliansky *et al*, 2008; Verchot-Lubicz *et al*, 2010; Harries & Ding, 2011). Nevertheless, to initiate systemic infection, viruses have to overcome sophisticated defense mechanisms set up by host plants such as RNA interference (RNAi) based on small interfering RNA (siRNA). RNAi is the main cellular mechanism involved in plant protection against invading RNA viruses (Pumplin *et al*., 2013). Viral dsRNA replicative intermediates or structured RNA genomes elicit cleavage by Dicer-like (DCL4) or its surrogate DCL2 to release 21 to 24 nts siRNA. siRNAs are loaded on Argonaute 1 (AGO1) or AGO2 to target viral RNA genome cleavage. To counteract this major defense mechanism, RNA phytoviruses have evolved viral suppressors of RNA silencing (VSR) capable of preventing RNAi-triggered immunity (Pumplin & Voinnet, 2013; Csorba *et al*, 2015). VSRs were intensively studied during last decades due to their crucial role in viral cycles and their potential for antiviral targeting molecules (Ghosh *et al*, 2017). Like MPs, VSR proteins do not show sequence or molecular organization similarities and often present different mechanisms to interfere with the host RNAi pathway (Lakatos *et al*, 2006; Zhang *et al*, 2006; Bortolamiol *et al*, 2007; Pumplin & Voinnet, 2013; Csorba *et al*, 2015; Baumberger *et al*, 2007). Among all VSRs described, only five complete structures were solved at atomic resolution among which the p19 protein of Tombusivirus (Vargason *et al*, 2003; Baulcombe & Molnár, 2004), B2 of Flock House virus (Chao *et al*, 2005), the Cucumovirus 2b protein (Chen *et al*, 2008), p21 encoded by the Beet Western Yellow virus (Ye *et al*, 2003) and NS3 of the Rice Hoja Blanca Tenuivirus (Yang *et al*, 2011). For some VSRs, solving 3D protein structure allowed unraveling interference mechanisms with the RNAi machinery (Xia *et al*, 2009; Ye *et al*, 2003; Vargason *et al*, 2003; Yang *et al*, 2011). This paucity of structural data contrasts with the numerous 3D structures of CP assembled in virion obtained up to know (DiMaio *et al*, 2015; Clare *et al*, 2015; Hesketh *et al*, 2015; Wang *et al*, 2015). Despite many improvements during last decades, getting 3D protein structures still represents a challenge, especially for individual viral proteins because of their intrinsic disordered or flexibility properties (Pushker *et al*, 2013; Xue *et al*, 2014). In plant viruses, just a few 3D structures of non-structural proteins are currently available, contrasting with the huge number of publications on viral proteins.

The sobemovirus *Rice yellow mottle virus* (RYMV) belongs to this plant virus category for which only the CP assembled virion 3D structure has been solved (Opalka *et al*, 2000). Revealed in 1966 in East Africa by Bakker (Bakker, 1970), RYMV rapidly spread in rice cultures from East to West Africa (Abubakar *et al*, 2003; Fargette *et al*, 2004; Traore *et al*, 2005) with a worrying epidemiological profile, generating crop losses ranging from 20 to 100% depending on the year and the environmental conditions. RYMV’s genome is a positive single-stranded, non-polyadenylated, linear RNA of 4.450 kb long. The genomic organization is related to the Cocksfoot mottle virus (CfMV), and includes 5 ORFs (Ling *et al*, 2013; Sõmera *et al*, 2015). The most emblematic multifunctional protein in RYMV is P1, a protein encoded by ORF1 of the genome. P1 is a small cysteine-rich protein of 18 kDa, first identified as required for virus movement (Bonneau *et al*, 1998; Siré *et al*, 2008), suspected to promote viral replication (Bonneau *et al*., 1998), and described as a viral suppressor of RNA silencing directed against exogenous genes in plant reporter systems (Voinnet *et al*, 1999; Siré *et al*, 2008; Lacombe *et al*, 2010; Fusaro *et al*, 2012). The P1 protein presents the highest diversity among the other viral proteins with 17,8% of amino acid sequence divergence (Siré *et al*, 2008; Sérémé *et al*, 2014). All functions achieved by P1 play a crucial role during early steps of viral infection, however little is known concerning its mode of action. P1 presents two couples of cysteines (Cys) organized in Cys-X_2_-Cys motifs similar to a zinc finger (ZnF) (Gillet *et al*, 2013) and highly conserved among RYMV P1 diversity (Cys 64-X_2_-Cys 67 and Cys 92-X_2_-Cys 95), and remarkably, in the P1 of sobemovirus genus (Sõmera *et al*, 2015). Point mutations introduced on Cys residue of each Cys-X_2_-Cys motif (Cys 64 and Cys 95) impaired silencing suppression and cell-to-cell movement in transgenic rice respectively, indicating a central role of Cys residues for P1 functions (Siré *et al*, 2008). Using a biochemical approach coupled with mass spectrometry, we showed that P1 binds two Zn atoms in its reduced state, whereas its oxidation induced complete Zn release accompanied by disulfide bonds formation, structural modification and oligomerization (Gillet *et al*, 2013). This process could be reverted by adding a reducing agent, indicating that P1 is a Zn binding protein with redox flexibility and oligomerization properties *in vitro*, highlighting the importance of Cys residues for P1 behavior. Nevertheless, the molecular organization of P1 and of its two Zn binding domains remains unclear and the precise role of P1 in viral infection widely unknown.

In this paper, we describe the elucidation of the structure and dynamics of the full multifunctional P1 protein from the *Rice Yellow Mottle Virus*, described both as a movement protein and as a VSR. By combining biochemistry, mass spectrometry and structural biophysics, we have determined the molecular organization of this multifunctional protein and localized two Zinc fingers (ZnF), one of them being a new HCHC-type of ZnF undescribed to date and providing both redox sensitivity and flexibility to the C-terminal portion with respect to P1 N-terminal region. By mutating residues identified as potential critical sites by the structure analysis, we have identified key areas involved in virus accumulation and movement into the host plant.

## RESULTS

### Delineation of P1 zinc binding domains

In a previous work, we found that P1 binds two zinc atoms in a reduced environment (Gillet *et al*, 2013), but the high number of rigorously conserved cysteine (7) and histidine (6) residues in P1 diversity (Appendix Fig S1) prevented the identification of the amino acids involved. To address this question, we designed a series of recombinant P1 regions (Fig 1A, Appendix Supplementary Text, Appendix Fig S2-S3, Appendix Table 1-Table 2) and evaluated their capacity to bind zinc using a 4-(2-Pyridylazo) resorcinol (PAR) labeling assays. Figure 1B shows that zinc detected by the PAR probe co-localized with all P1 regions assayed in a SDS-PAGE. Re-sizing [1-100] and [50-157] regions into shorter [50-100] and [102-157] amino acid regions, respectively, allowed us to definitely assign the zinc binding capacity to these two latter central and C-terminal regions of P1 (Fig 1B). Native ESI-MS confirmed the zinc binding at [50-100] and [102-157] amino acid regions (Table 1, Appendix Fig S4-S5), further termed ZnF1 and ZnF2, respectively.

**Figure 1.**
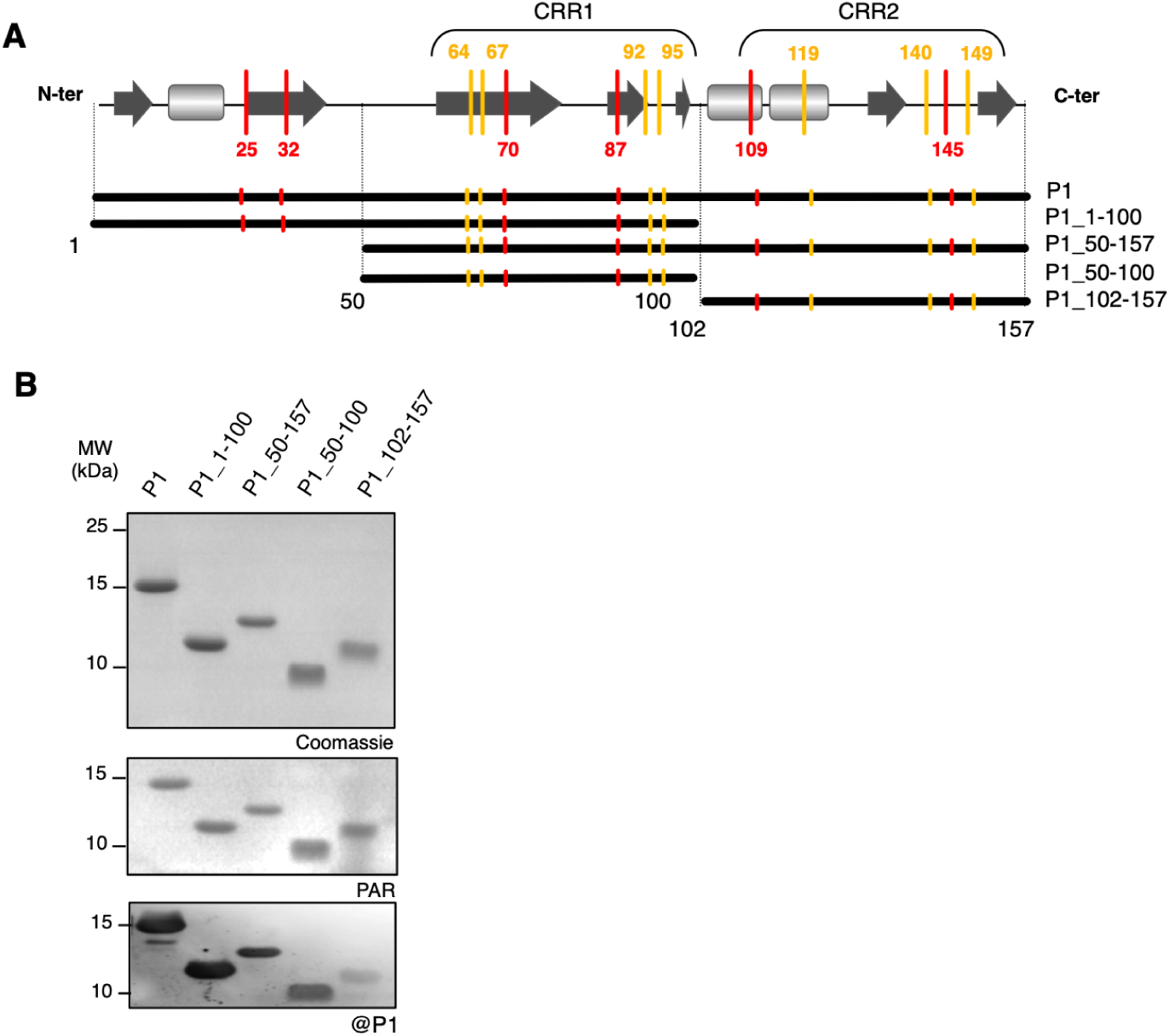
The viral RYMV P1 protein binds two zinc atoms at its central and C-terminal regions. **A** Schematic representation of P1 regions designed for production of recombinant proteins in *E. coli* BL21. Cys and His residues outside or inside cysteine-rich regions (CRR) are positioned along the P1 scheme with yellow and red bars respectively. Predicted secondary structures are represented by dark grey arrows for β-sheets and by light grey squares for α-helices. Resizing the full-length P1 protein into different sub regions is represented by thick black lines under the predicted secondary structure, with related P1 region names referring to the size of each truncated protein (amino acids numbering). **B** *In-gel* co-localization of zinc with P1 recombinant sub regions. Purified recombinant proteins were separated on non-reducing SDS-PAGE 18%. Zinc detection was performed by gel incubation in a Tris 50 mM pH 8 buffer containing the PAR probe (mid panel) before staining by Coomassie blue (upper panel). All proteins assayed were recognized by purified anti-P1 antibody used at a 1:1000 dilution in an immunoblot assay (lower panel).

**Table 1.** Quantification of zinc atoms in P1 recombinant sub regions and incidence of EDTA treatments on Zinc release. The incidence of EDTA treatments on P1 sub domains was analyzed by native ESI-MS. The molecular mass determined in Daltons for each isomer is compared to that obtained using denaturing ESI-MS and to theoretical values determined based on the primary sequence of each P1 sub region (Appendix Fig S4). Only values from major forms without zinc, or with one or two zinc atoms of the different isomers comprising the N-terminal methionine are given. Deconvoluted ESI mass spectra for the different proteins are given in the Appendix Fig S5 and Appendix Fig S6. Both P1_50-100 and P1_50-157 proteins underwent immediate degradation after EDTA treatments, independent of the concentrations used and did not produce any signal during ESI-MS analyses.

| P1 proteins | Theoretical molecular mass | Molecular mass by denaturing ESI-MS | Molecular mass by native ESI-MS before EDTA | Difference between native ESI-MS and theoretical mass | Number of zinc atoms (63 to 65 Da) before EDTA | Molecular mass by native ESI-MS after EDTA | Number of zinc atoms (63 to 65 Da) after EDTA | Molecular mass by native ESI-MS after EDTA + ZnOAc treatment | Number of zinc atoms (63 to 65 Da) |
| --- | --- | --- | --- | --- | --- | --- | --- | --- | --- |
| P1 | 17929.01 | 17928,2 ± 0,1 | 18056.1 ± 0.3 | 127,09 | 2 | 17992.2 ± 0.3 | 1 | 18056.6 ± 0.4 | 2 |
| P1_1-100 | 11408.80 | 11408,81 ± 0,31 | 11471.66 ± 0.04 | 62,86 | 1 | 11472.61 ± 0.84 | 1 | - | - |
| P1_50-100 | 5816.57 | 5816,13 ± 0,01 | 5880.1 ± 0.3 | 63,53 | 1 | No signal | - | No signal | - |
| P1_50-157 | 12336.78 | 12338,12 ± 0,39 | 12462.3 ± 0.3 | 125,52 | 2 | No signal | - | No signal | - |
| P1_102-157 | 6556.26 | 6555,8 ± 0,4 | 6620.31 ± 0.22 | 64,05 | 1 | 6556.46 ± 0.4 | 0 | 6620.34 ± 0.27 | 1 |

**Table 2.** Data collection and refinement statistics.

|  | P1_1-100 | P1_102-157-phasing | P1_102-157 |
| --- | --- | --- | --- |
| **Data collection** |  |  |  |
| Wavelength (Å) | 0.976 | 1.28 | 1.24 |
| Space group | F 41 3 2 | P 1 21 1 | P 1 21 1 |
| **Cell dimensions** |  |  |  |
| a, b, c (Å) | 195.61 | 38.14 31.23 42.03 |  |
| $\alpha, \beta, \gamma$ (°) | | 90.00 101.08 90.00 | 90.00 101.10 90.00 |
| Resolution (Å)<sup>a</sup> | 112.94-2.10 (2.21-2.10) | 41.25-2.10 (2.21-2.10) | 37.37-1.98 (2.09-1.98) |
| Rmerge | 0.051 (0.559) | 0.154 (0.448) | 0.098 (0.573) |
| N° of reflections | 724744 (104965) | 55242 (7957) | 22349 (3282) |
| N° of unique reflections | 19296 (2756) | 5834 (839) | 6911 (992) |
| *I*/*sI* | 8.3 (1.1) | 12.4 (5.0) | 8.6 (2.0) |
| Completeness (%) | 99.97 (100.0) | 100.0 (99.9) | 99.2 (99.4) |
| Multiplicity | 37.5 (38.1) | 9.5 (9.5) |  |
| Anomalous completeness | 100.0 (100.0) | 99.9 (99.9) |  |
| Anomalous multiplicity | 20.1 (19.9) | 4.6 (4.5) |  |
| **Refinement** |  |  |  |
| R<sub>work</sub> / R<sub>free</sub> (%) | 0.185 / 0.214 |  | 0.193 / 0.227 |
| N° atoms | 1688 |  | 899 |
| Protein | 1547 |  | 847 |
| Zn | 3 |  | 2 |
| Water | 140 |  | 50 |
| B-factors (Å<sup>2</sup>) |  |  |  |
| Protein | 45 |  | 27 |
| Zn | 52 |  | 22 |
| Water | 53 |  | 29 |
| rmsd |  |  |  |
| Bond lengths (Å) | 0.009 |  | 0.007 |
| Bond angles (°) | 1.166 |  | 1.069 |
| Ramachandran favored (%) | 98.95 |  | 97.88 |
| Ramachandran outliers (%) | 0.0 |  | 0.0 |

### ZnF1 and ZnF2 in P1 exhibit different zinc binding affinity

Our previous analyses suggested that P1 binds its two zinc atoms with differential affinity (Gillet *et al*, 2013). To further address this question, we first performed EDTA treatments followed by native ESI-MS analyses on the full-length P1 protein. We found that treatments up to 400 molar equivalents (eq) of EDTA were sufficient to provoke the release of one zinc atom (Table 1 and Appendix Fig S6A, compare upper and mid panels). This zinc atom could partially reload into P1 after treatment of the EDTA-chelated protein with 100 µM ZnOAc (Table 1 and Appendix Fig S6A, lower panel). All attempts to provoke the release of the second zinc atom from P1 using higher EDTA concentrations were unsuccessful.

Similar assays were then performed on P1 shorter regions [1-100] and [102-157], the two other proteins P1_50-100 and P1_50-157 being improper for ESI-MS analyses after EDTA treatments. Treatments up to 400 molar equivalents of EDTA applied to the ZnF1-containing P1_1-100 protein failed to release the unique zinc atom present in this shorter P1 protein (Table 1 and Appendix Fig S6B). In order to determine ZnF1 zinc affinity constant, we first subjected P1_1-100 (40 µM) to N,N,N,N-Tetrakis(2-pyridylmethyl)-ethylenediamine) (TPEN), a heavy metal chelator with a Ka constant of 10^16^ M^-1^ (Anderegg *et al*, 1977) stronger than that of EDTA (Ka _EDTA_ = 10^14^ M^-1^) (Smith & Martell, 1989). After chelation by TPEN, we could estimate a zinc affinity constant Ka_ZnF1_ of 2,15×10^16^ M^-1^, using a PAR/PMPS zinc release assay according to (Jakob *et al*, 2000) (Appendix Fig S7).

Regarding ZnF2, a treatment with 50 eq of EDTA was sufficient to release the zinc atom (Table 1 and Appendix Fig S6C), indicating that the ZnF2 domain is much less affine for zinc than ZnF1. Despite several assays, we were unable to determine the zinc affinity constant of ZnF2, suggesting that ZnF2 zinc affinity constant could be lower than that of EDTA itself. We additionally observed that the chelated P1_102-157 protein could reload one zinc atom similarly to the full-length P1 protein once treated with 60 µM ZnOAc (Table 1 and Appendix Fig S6). This indicates that the zinc atom rapidly released from the full length P1 protein after EDTA treatment and easily reloaded is the one bound to the ZnF2 domain.

Because oxidizing environments provoke zinc release from P1 and its oligomerization, (Gillet *et al*, 2013) (Appendix Fig S8A), we also investigated whether the differential zinc-binding affinities of ZnF1 and ZnF2 were related to the respective redox sensitivity of [1-100] and [102-157] regions. Using SDS-PAGE under non-reducing conditions, we found that high H_2_0_2_/protein ratios (80 eq corresponding to 2 mM) were required to induce the release of zinc atom from the recombinant P1 and P1_[1-100] proteins, respectively, without affecting their mobility and folding (Appendix Fig 8B, left panel). Reversely, lower H_2_0_2_/protein ratios (10 eq corresponding to 1 mM) were sufficient to provoke zinc release from the recombinant P1_[102-157] protein, accompanied by a mobility shift due to oligomerization and slight conformational changes observed by CD (Appendix Fig 8C).

Altogether, these results demonstrate that the N-terminal [1-100] region of P1 comprising ZnF1 is the most affine domain to zinc and also the less sensitive to oxidizing environments, both properties conferring strong structure maintenance of this region. They also identify P1 C-terminal [102-157] domain as the most sensitive region to chelation or oxidant molecules and the most amenable to structural flexibility, sustained by the low zinc affinity of ZnF2.

### P1 N- and C-terminal regions exhibit different structural and zinc binding geometries

To go further in the structural characterization of P1, we subjected the protein to crystallography. As all attempts to crystallize the whole protein remained unsuccessful, we took advantage of the different soluble recombinant regions of P1 (Fig 1A) for further analyses, of which P1_1-100 for N-terminal region and P1_102-157 for C-terminal region (P1_102-157) were successfully crystallized at a resolution of 1.9 and 2.1 Å, respectively (Fig 2). All crystallographic and refinement statistics are given in Table 2.

**Figure 2.**
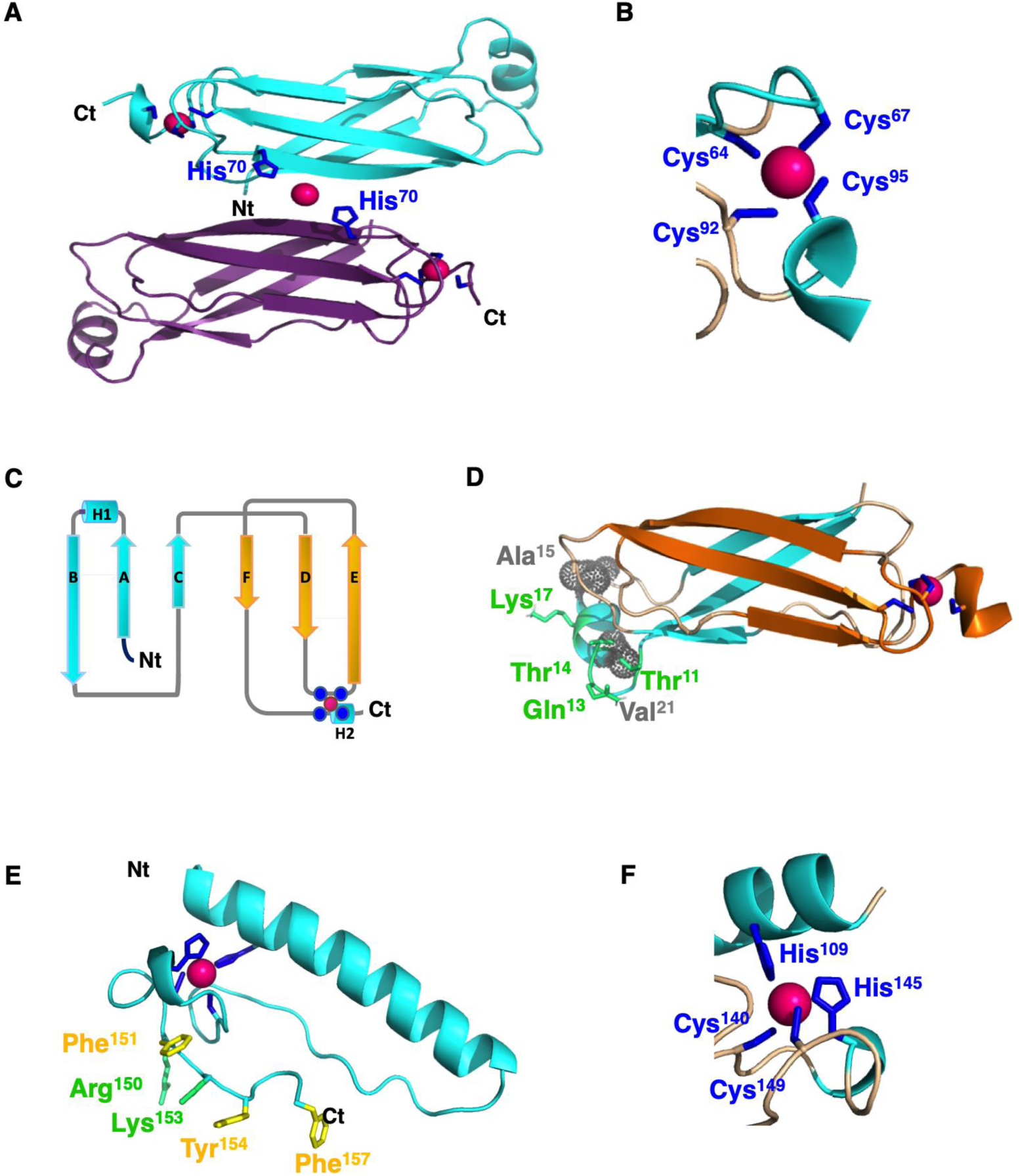
Crystallographic structure of P1 N- and C-terminal regions. **A** The crystal structure of P1_1-100 (N-terminal region) reveals a dimer in an anti-parallel configuration. Each monomer contains a zinc coordination site constituted by the four cysteines Cys64, Cys67, Cys92 and Cys95 (Cys treble-clef). In addition, a zinc atom (represented as a pink sphere in all panels) is also present at the dimer interface, where it is chelated by the His70 side chain (dark blue sticks) of each monomer. Nt and Ct labels point to the N-terminal and C-terminal parts of P1_1-100, respectively. **B** Close-up of the Cys-treble clef of P1_1-100, with the side chains of zinc-chelating cysteine residues in stick representation. **C** Schematic diagram representation of P1_1-100 structure. Helices and β-sheets are depicted as cylinders and arrows, respectively. Cys residues chelating the zinc atom are represented as blue plain circles. **D** Each monomer of P1_1-100 presents a small helix with a Lysine side chain residue (Lys17) completely solvent exposed and tolerating only Lys to Arg substitutions. The helix position is stabilized by contacts of hydrophobic residues (side chains represented as grey dotty spheres) which interact with the protein core, and by a network of hydrogen bonds involving the conserved Thr14, Gln13 and Thr11 residues. **F** Cartoon representation of P1_102-157 ribbon. Side chains of conserved positively charged and aromatic residues are depicted in stick representation, in green and yellow respectively. The side chains of the HCHC ZnF2 are colored in blue. **E** Close-up of the HCHC ZnF2 in P1_102-157, with the side chains of zinc-chelating cysteine and histidine residues in stick representation.

A remarkable feature of P1_1-100 crystal structure was the formation of a dimer from paired antiparallel monomers (Fig 2A), a property that we confirmed by using an *in vivo* yeast-two hybrid assay (Appendix Fig S9). Each monomer contains a zinc atom (Fig 2A-B) and consists in a sandwich composed of 6 strands (A-F, Fig 2C-D) and 2 short helices (H1 and H2, Fig 2C-D) at each extremity. The overall sandwich structure itself arises from the duplication of a subdomain containing three strands, the two sub domains (A-C and D-F strands) superimposing with a r.m.s.d. of 1 Å on Cα atoms. Searching for similar folds among known structures with DALI (Holm & Rosenström, 2010) returned only one significant hit with SCOP fold families (sandwich, 6 strands, 2 sheets), suggesting that P1_1-100 could be documented as a new member of this family. The presence of the zinc finger at the C-terminal extreme side of the sandwich, involving Cys64, Cys67, Cys92 and Cys95 as zinc ligands (Fig 2A-B), validates the identification of a CCCC-type ZnF1 domain at the [50-100] and [1-100] regions of P1 by our SDS-PAGE/PAR coupled assays and ESI-MS analyses (Fig 1B, Table 1). The importance of this CCCC-type ZnF1 in the maintenance of the sandwich fold and P1 solubility was demonstrated by additional SDS-PAGE and ESI-MS experiments on single and double cysteinic and histidinic recombinant mutants (Appendix Supplementary Results and Appendix Table 1 & 2) of both the entire P1 (Appendix S10 and S11) and its P1_50-100 region (Appendix S12 and S13), this latter region corresponding to the D-F subdomain (Fig 2). According to the crystal structure, both Cys64 and Cys67 belong to a β turn between strands D and E, whereas Cys95 is located at the beginning of helix 2 (Fig 2B-C). Hence, ZnF1 belongs to the treble clef group, albeit not a canonical form as Cys92 does not belong to helix 2 (Krishna *et al*, 2003; Kaur & Subramanian, 2016). Another remarkable feature of P1_1-100 is the presence of a helix harboring a lysine (Lys17) completely solvent exposed (Fig 2D). This lysine is highly conserved within P1 diversity and position 17 indeed tolerates only replacement by arginine residues (one sequence among 51 inspected, Appendix Fig S1), suggesting that this residue could be a potential site for interaction with protein or nucleic acid partners. Interestingly, this helix is stabilized by a network of hydrogen bonds between highly conserved polar residues (Thr11, Gln13 and Thr14, Fig 2D and Appendix Fig S1) and by contacts between the hydrophobic Ala15 and Val21 residues, whose lateral chains face a hydrophobic pocket within the sandwich.

Contrary to P1_1-100, the final model of P1_102-157 at the resolution of 2.1 Å was monomeric. This observation is in accordance with the absence of homo-dimerization ability of this P1 region in an Y2H assay (Appendix Fig S9). P1_102-157 crystals revealed an original fold consisting in one α helix (residues 103-126) at the N-terminal region of this domain and a rigid part devoid of any secondary structure (residues 127-157, Fig 2E). This rigid part contains three zinc ligands Cys140, His145 and Cys149 whereas the fourth ligand His109 belongs to the N-terminal helix (Fig 2F). The compact structure of this P1 C-terminal region is ensured by the zinc finger folding, as demonstrated by additional SDS-PAGE experiments on cysteinic and histidinic mutants of this 102-157 region (Appendix Supplementary Results, Appendix Fig S12B). EDTA-mediated zinc chelation promoted its unfolding, as shown by its ^1^H NMR spectrum (Appendix Fig S14). These observations validate the identification of a HCHC-type ZnF2 domain at the [102-157] region of P1, confirming our previous predictions (Gillet *et al*, 2013). The striking architecture of P1 ZnF2 (3 ligands in a loop and one on a helix, Fig 2F) does not belong to any existing class of Zn finger (Krishna *et al*, 2003). Searching for similar structures with DALI neither returned significant hits. Thus P1 ZnF2 seems to be a member of a new ZnF class, not only because of its typology, but also because of its “HCHC” box nature. Another striking feature of this domain is its high number of positively charged residues, most of them with their side chain exposed to solvent (Fig 2E). In particular, the extreme C-terminal part presents a patch of highly conserved solvent-exposed lysine and arginine residues (Arg150 and Lys153) which only tolerates Arg to Lys or Lys to Arg replacement, overlapping with a stretch of aromatic residues (Phe151, Tyr154 and Phe157) which tolerates no or only replacement with aromatic residues (Tyr154 excepted, for which a replacement with a cysteine is also observed in one sequence among the 51 inspected) (Appendix Fig S1).

### Deciphering the overall P1 structure by NMR

To unravel the structure of the full-length P1 protein, we led a parallel NMR investigation on the full length P1 and its two sub-regions, P1_1-100 and P1_102-57. Backbone ^15^N and ^1^H chemical shifts are exquisite reporters of protein conformation. The perfect superimposition of cross peaks between the ^1^H-^15^N HSQC spectrum of full length P1 and that of its [1-100] and [102-157] regions (Fig 3A and 3B, respectively) established that the structure of the individual regions solved by X-Ray (Fig 2C and 2E) is conserved in the whole protein. It also indicates that the interaction between both N-terminal and C-terminal domains of P1, if any, was minimal which is supported by the absence of interaction between P1_1-100 and P1_102-157 in a yeast two-hybrid assay (Appendix Fig S9). The titration of ^15^N-labelled C-terminal domain by the non-labeled P_1-100 (Fig 3C) neither showed evidence of interaction. Furthermore, analysis of the chemical shifts (Hα, ^13^Cα, ^13^Cβ, CO) by TALOS (Shen & Bax, 2013) (Appendix Table 3) and of the ^1^H-^1^H NOEs (Fig 3D) in the linker region established that the C- and N-terminal helices seen in X-Ray structures of P1_1-100 and P1_102-157 (Ala94 to Glu96 and Glu104 to Glu126, respectively) were extending to the entire linker region between both domains in the full-length protein. This folding of the linker into a helix was retrieved in structure calculations, where the linker was only restrained by ^1^H-^1^H NOEs and TALOS-derived dihedral angles. In these calculations of full P1 structure, the N- and C-terminal domains were fixed to the X-Ray structures, with their mutual orientation restrained by residual dipolar coupling constants (RDC) (Fig 4A). The helix linking both domains is however not entirely rigid. Both RDC measurements (Fig 3E) and ^15^N relaxation data (Fig 3F and Appendix Table 4) established that P1_1-100 and P1_102-157 domains have some degree of flexibility with respect to each other. RDC measurements enabled the derivation of a global order parameter of 0.78 of ZnF2 relatively to ZnF1 (Tolman *et al*, 2001). This value corresponds to a motion amplitude of ± 30 degrees, following the formalism of Tolman under the assumption of an axially symmetric “wobble-in-a-cone” movement (Tolman *et al*, 2001).

**Figure 3.**
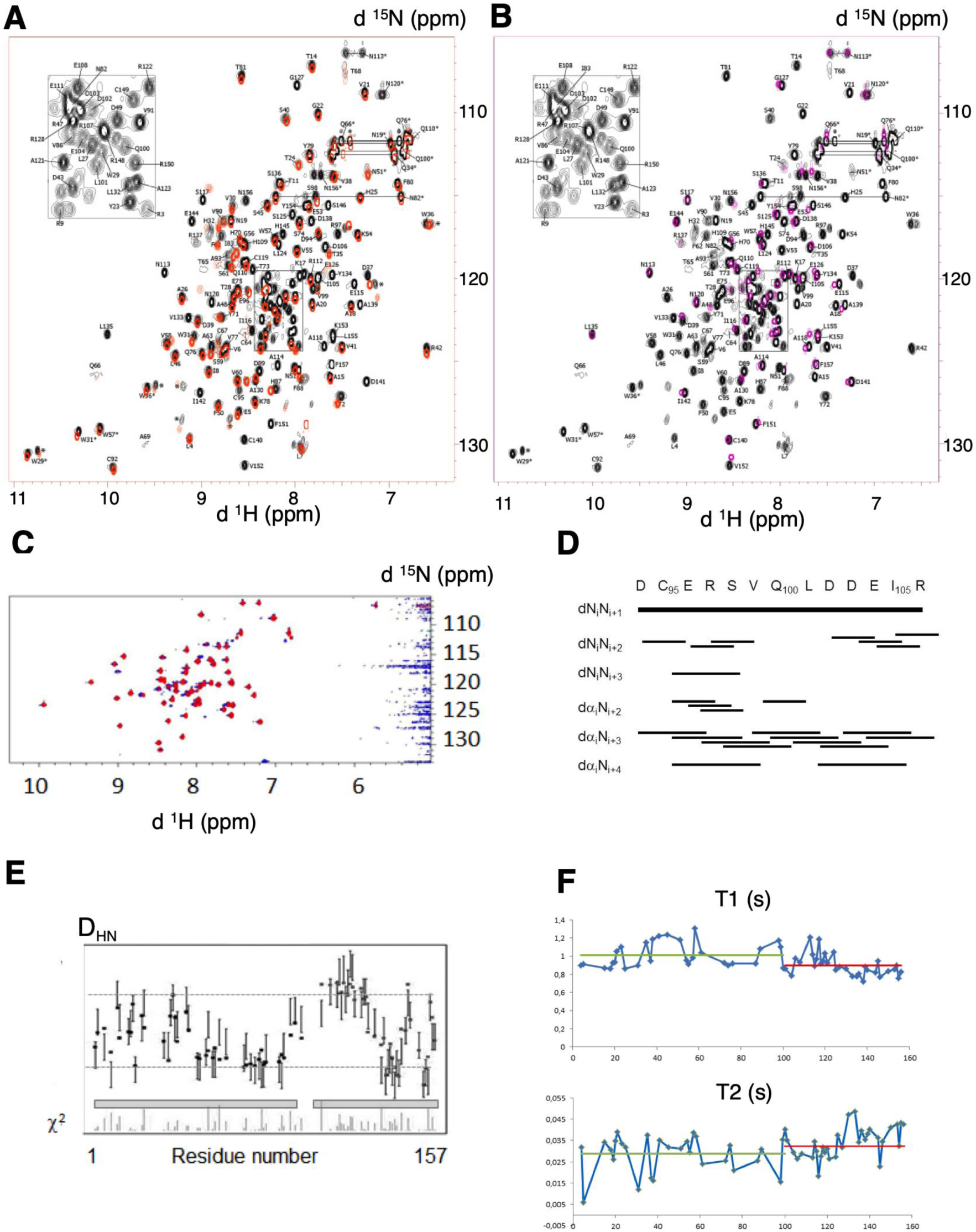
NMR characterization of P1 fragments and full structure. **A** Superposition of fingerprint regions of ^1^H-^15^N HSQC of full length P1_1-157 (black cross peaks) and P1_1-100 (red cross-peaks). Sequence specific resonance assignments are indicated by number for the backbone amide atoms and the one-letter amino acid code. Pairs of side-chain NH_2_ resonances are connected by *horizontal lines*. Tryptophan side chain indole peaks are also indicated with asterisks (*). **B** Superposition of fingerprint regions of ^1^H-^15^N spectra HSQC of full length P1_1-157 (black cross peaks) and P1_102-157 (magenta cross-peaks). Sequence specific resonance assignments are indicated by the one-letter amino acid code and the residue number. Pairs of side-chain NH_2_ resonances are connected by *horizontal lines*. Tryptophan side chain indole peaks are also indicated with asterisks (*). **C** Superposition of fingerprint regions of ^1^H-^15^N spectra HSQC of ^15^N-labelled P1_102-157 (c= 0.5 mM) alone (blue) and in the presence of 2 equivalents of non labeled P1_1-100 (c= 1 mM). The resonances of the P1_102-157 are identical in terms of line width and position, supporting the lack of interaction between domains. **D** Sequential and medium range NOEs reported between proton atoms for the linker sequence (amino acids 94-106) characterizing their folding into a helix. **E** ^1^D_HN_ residual dipolar couplings measured for full-length P1_1-157 in phage medium depicted as a function of the protein sequence. The positions of P1_1-100 and P1_102-157 regions are indicated as grey segments, with maximal ^1^D_HN_ values of the P1_1-100 domain enlightened by dotted lines. The P1_102-157 domain exhibits several ^1^D_HN_ below and above these threshold values, showing that both regions exhibit different alignment properties. The individual χ^2^ for each amino acid i: (Dcalc(i)-Dexp(i))^2^/∑_tot_ Dexp(i)^2^ is also represented on the bottom line. **F** ^15^N T_1_ and T_2_ relaxation times recorded for full-length P1_1-157, as a function of residue number. The mean values for P1_1_100 and P1_102-157 domains are depicted as green and red horizontal lines, respectively. The differences observed support a significant degree of flexibility between them.

**Figure 4.**
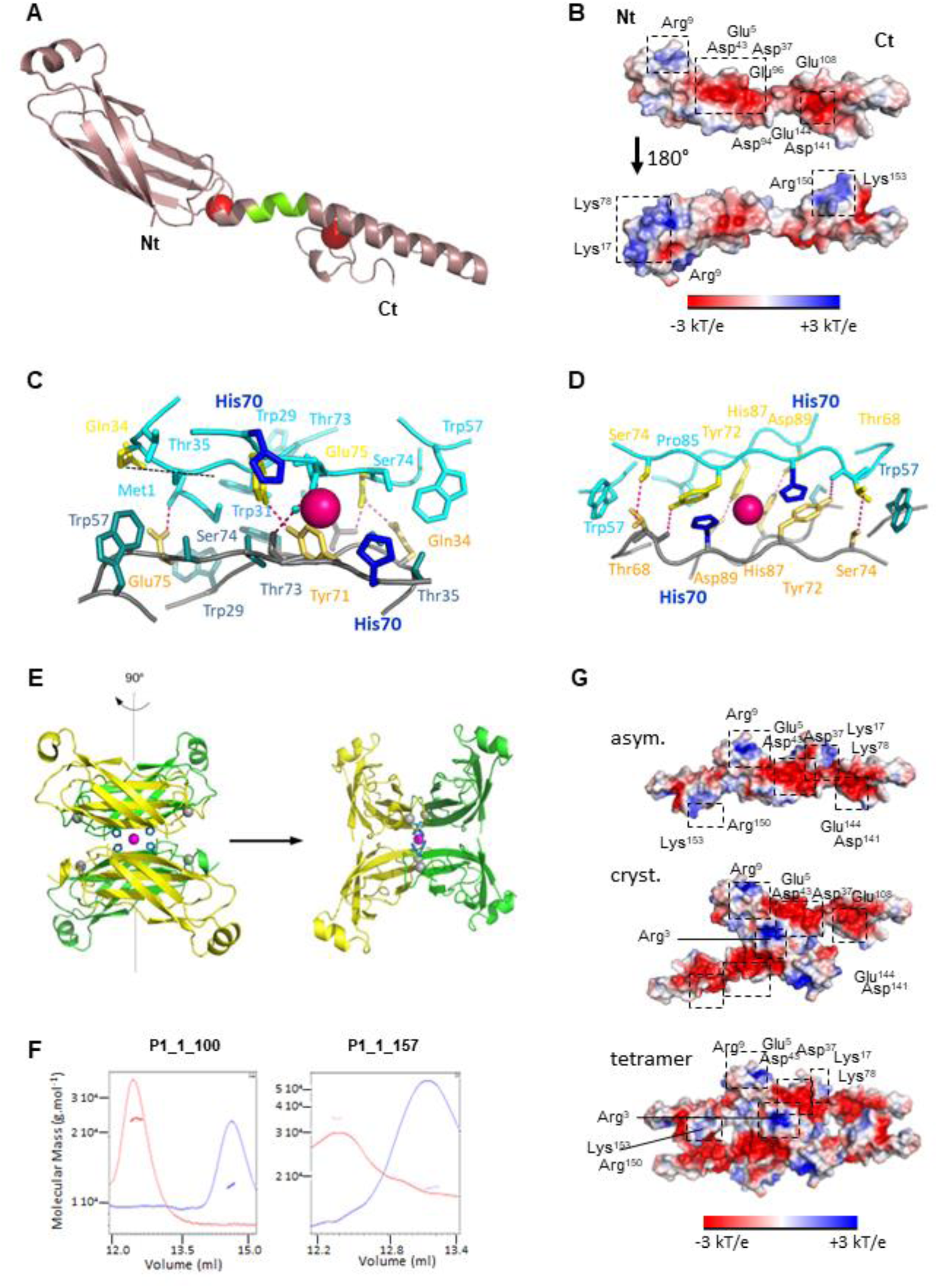
Structure of full-length P1 and oligomerization states by NMR. **A** Model of full-length P1_1-157 built from rigid-body calculations based on the crystallographic structures of individual P1 domains and NMR restraints. The missing segment in crystallographic structures in the linker region has been reconstructed on the basis of NMR restraints; it adopts a helicoidal fold (green portion) extending the long helix of the second zinc finger. **B** Electrostatic surface potential representation of the full length P1, showing on one side a large central negatively charged surface and on the other side, two extremal positive poles. Conserved positive and negative patches among the reported 51 P1 sequences are framed. **C** Dimer interface of the asymmetric unit. The central Zn atom is represented as a pink sphere, chelated by His70 (dark blue) in stick representation. Side chains of residue involved in polar and hydrophobic contacts in the dimeric interface are represented in stick and are colored in yellow and cyan blue, respectively. **D** Dimer interface of the crystallographic dimer. The central Zn atom is represented as a pink sphere, chelated by His70 (dark blue) in stick representation. Side chains of residues involved in polar and hydrophobic contacts in the dimeric interface are represented as stick colored in yellow and cyan blue, respectively. **E** Cartoon representation of the P1-Nt tetramer (UA in green). Central Zn atom is represented in magenta and chelating His70 residues, as blue, stick form balls. **F** Zinc-dependent oligomerization of the P1_1-100 and full length P1_1-157 proteins. SEC MALLS profile of the P1_1-100 and full length P1_1-157 is depicted. UV absorbance is shown in red in buffer containing zinc and **G** Representation of the electrostatic surface of the asymmetric P1 dimeric model (above), of P1 crystallographic dimer (middle) and of tetrameric (below) P1. Conserved positive and negative patches are framed.

In the NMR-derived structure, the whole P1 protein presents a surface with an original polarized profile (Fig 4B). On one face, the central region harbors a large negatively charged patch, formed by conserved Glu5, Asp37, Asp43, Asp94, Glu96, Glu108, Asp141, and Glu144. The other side harbors two extremal positively charged poles at the N- and C-terminal parts, which overlap with the potential interaction motifs described above. Indeed, the first positive pole is constituted in ZnF1 by Arg9, Lys17 and Lys78 side chains forming a hydrophilic and positively charged cleft. The second positive pole is constituted in ZnF2 by the C-terminal extremity (Arg150, Lys153), which alternates with aromatic residues (Fig 2E).

### Oligomeric structure of P1

The superimposition of two [1-100] monomers in the crystal of P1_1-100 (Fig 2A) and yeast-two hybrid assays (Appendix Fig S9) indicated that P1 forms a dimer through its N-terminal region. In the asymmetric unit (Fig 4C and Appendix Fig S15A, upper scheme), the dimeric interface involves five hydrogen bonds between residues Met1 and Glu75, Gln34 and Glu75, Tyr71 and Tyr72, together with 10 hydrophobic contacts involving residues Gln34, Gln76, Trp29, Trp31, Glu75, Ser33, Ser74, Trp57, Thr35, Thr73. The dimer interface is further strengthened by four salt bonds between Met1 and Glu75. The overall interface surface was estimated at 506 Å^2^ (Appendix Fig S15B and Appendix Table 5), hence 8.4 % of the total surface of one monomer (6019 Å^2^), classifying this dimeric form as labile. Accordingly, ESI-MS analyses detected only a faint proportion of P1 dimers for full length P1 (Gillet *et al*, 2013).

Analysis of the crystal packing indicated that P1_1-100 could also undergo into a tetrameric structure stabilized by the coordination of a fifth zinc atom by the 4 His70 residues belonging to the 4 P1 N-terminal regions (Fig 4E). This crystallographic interface, estimated at 574 Å^2^, involves six hydrogen bonds linking His87 with Asp89, Thr68 with Tyr72, and Ser74 with Thr68 of each monomer, but also twenty hydrophobic contacts implicating Pro85, His70, Tyr72, Thr68, Gln66, Ser79, and Trp57 in each monomer. This interface is also strengthened by Ser59-Ala63 and Ser61-Ser61 saline bonds via the bridging of water molecules. To challenge such possibility, we subjected the full-length P1 and its truncated form P1_1-100 to size-exclusion chromatography coupled with multi-angle laser light scattering (SEC-MALS). At low salt concentrations, both the full length and its N-terminal region [1-100] eluted as monomers in a zinc-free elution buffer and as a dimer in a zinc-containing elution buffer (Fig 4F). No trace of higher molecular weight proteins could be detected. On the other hand, NMR relaxation studies performed at high salt concentration were in favor of a monomer (Appendix Table 4). No relaxation data could be recorded at low salt concentration, as the protein was unstable in the protein concentration range needed (200 μM).

We next built the corresponding models of homodimeric and tetrameric P1 models based on the NMR-derived full-length P1 structure. No steric clash was observed. Interestingly, the dimeric interface (asymmetric unit) mixes the characteristics of the electrostatic surface of both sides of P1 monomeric form, in particular the presence of the central large negative patch and of the extremal positive poles in ZnF1 and Znf2 (Fig 4G). By contrast, a doubled negative central patch is observed in P1 built from the crystallographic dimer, interrupted by a positive spot constituted by the two side chains of the conserved Arg3 residue of each protomer. Consequently, the tetrameric form of P1 maintains the presence of the large central negative patch and of the extremal positive poles in ZnF1 and ZnF2, but the access to the positive and aromatic pole of ZnF2 (Arg150, Phe151, Lys153, Tyr154, Phe157) is hampered (Fig 4G).

### Oligomerization of P1 N-terminal region is essential for viral spread in rice

To evaluate the biological importance of P1 structural features and oligomerization properties in RYMV infectivity, we took advantage of FL5, a RYMV DNA clone allowing the production of infectious viral RNAs that can be directly inoculated in rice plantlets (Brugidou *et al*, 1995). P1 residues selected for their potential key position within P1 structure were therefore mutated in the P1 ORF within the FL5 sequence and rice plants infected by viral RNAs carrying mutations were recorded for RYMV fitness and infectivity. We first focused on the short solvent-exposed [Lys17-Val21] helix at the N-extremal part of the P1 sandwich that is potentially accessible for sandwich structure maintenance and for binding of intracellular partners (Fig 2D). By comparison with plants inoculated with infectious FL5 RNAs (Fig 5) and with a non-infectious FL5ΔP1 clone impaired in P1 translation (Fig 5, Appendix Supplementary Results and Appendix Fig S16), we found that inoculating plants with FL5 RNAs depleted of the P1 DNA region encoding residues 12 to 21 containing this protruding helix (FL5-P1^Δ12-21^, Fig 5A) did not provoke viral infection. Only very few amounts of viral particles were detected as revealed by DAS-ELISA, and inoculated plants showed similar height (Fig 5B) and green healthy leaves (Fig 5C) to plants inoculated with the non-infectious FL5ΔP1. Interestingly, we failed to produce a P1 recombinant protein depleted of this helix in *E. coli* cells, suggesting that the short N-terminal helix in P1 has a great importance for its folding. To go further, we used plant protoplasts as a reporter system to evaluate P1 behavior *in vivo*. By comparison with wild-type P1-EGFP and EGFP-P1 fusion proteins that both locate in the cytosol and in the nucleus in this reporter system (Appendix Fig S17), both P1^Δ12-21^-EGFP and EGFP-P1^Δ12-21^ fluorescent fusion proteins had low fluorescence capacities and a tendency to aggregate in the cytosol. Altogether, these experiments indicate that the short helix at the N-terminal region of P1 is essential at least for P1 folding.

**Figure 5.**
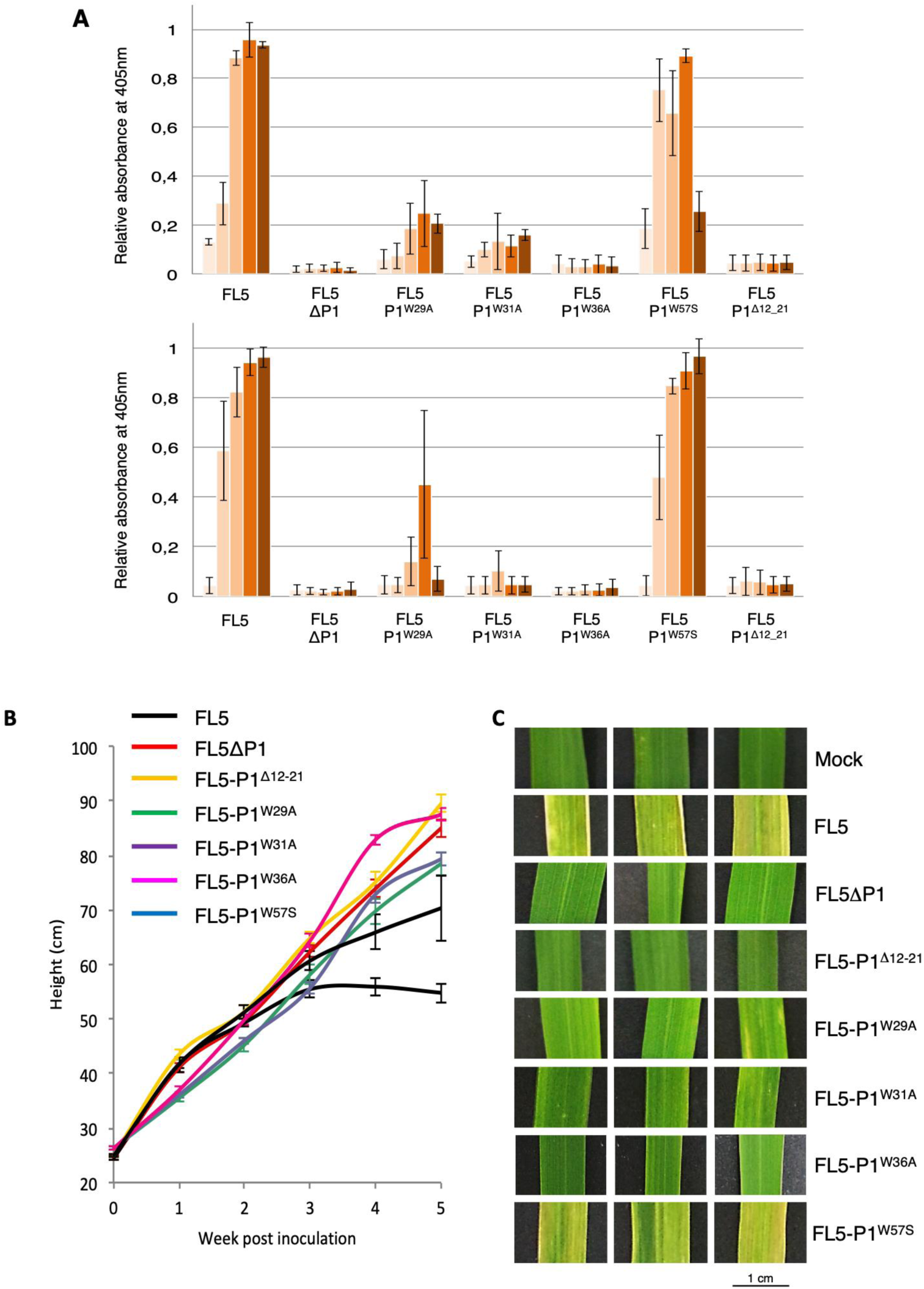
Incidence of mutations in P1 N-terminal region residues on RYMV infectivity in rice. **A** Consequences of alpha helix [12-21] deletion and of tryptophan residue mutations in P1 on RYMV particle accumulation in rice. Viral charge of rice plantlets infected by FL5-P1^Δ12-21^, FL5-P1^W29A^, FL5-P1^W31A^, FL5-P1^W36A^, and FL5-P1^W57S^ RNAs was recorded by DAS-ELISA on total proteins extracted from inoculated (upper panel) and systemic (lower panel) leaves collected every week during 5 weeks (light to dark bars). An antibody directed against the RYMV coat protein CP was used in DAS-ELISA for the detection of viral particles. The original infectious clone FL5 (Brugidou *et al*, 1995) and its non-infectious version FL5ΔP1 (Appendix Fig S16) were used as positive and negative controls for infection and viral systemic spread, respectively. **B** Plant height (in cm) of inoculated plants recorded all along the 5-week infection process. **C** Yellowing symptom recording at the leaf surface of inoculated plants. Three samples of rice leaves are shown as representative of symptoms visible at the overall plant scale. Leaves were photographed 5 weeks post inoculation. **Data information:** The first 3 leaves of 14-day old rice plantlets were selected for inoculation by synthetic viral RNA or by RYMV particles, while the following ones from the 4^th^ leaf corresponded to systemic tissues. Synthetic FL5-derived RNAs (5 µg) were used in a phosphate/carborandum buffer for inoculation, while mock plants were treated similarly but without RYMV-derived molecules (Appendix Supplementary Materials). All data were obtained from 30 plants per treatment, in two independent inoculation assays.

We also evaluated the role of tryptophan residues present in P1 N-terminal region by creating Trp-to-Ala or Trp-to-Ser replacements in the P1 sequence (Appendix Table 1). Trp residues (4 in P1) were selected firstly because they all locate in this region that undergoes oligomerization according to our data. Trp29, Trp31 and Trp57 have their side chain exposed to solvent and are accessible to intracellular partners in the monomeric form, and participate to hydrophobic contacts in the asymmetric interface (Fig 3D, Appendix Fig S15 & S18A). Trp57 additionally participates in the crystallographic interface, while Trp36, which is buried in a hydrophobic core of ZnF1 (Appendix Fig S18B), locates on the same β-strand as Trp29 and Trp31. All these Trp are strictly conserved among P1 diversity within the RYMV sobemovirus gender (Sérémé *et al*, 2014), which suggests their importance for P1 functions. Rice plants were therefore infected by viral genomic RNAs expressing Trp-mutated P1s and recorded for RYMV fitness and infectivity. By comparison with infection levels induced by the original FL5 RNA, we found that viral RNAs expressing either P1^W29A^ or P1^W31A^ proteins only induced a faint production of RYMV particles in infected leaves (Fig 5A), with a slight growth decrease only (Fig 5B), and with RYMV symptoms limited to very few yellowing areas, mostly in plants inoculated with FL5-P1^W29A^ RNAs (Fig 5C). This FL5 variant appeared as the sole infectious clone able to provoke a moderate accumulation of viral particles in systemic leaves (Fig 5A-C). Mutating Trp36 in P1 within FL5 (FL5-P1^W36A^, Fig 5A) provoked a more important defect in RYMV particle accumulation in inoculated leaves, abolishing RYMV systemic movement to non-infected tissues (Fig 5A). Reversely, mutating Trp57 had minor effects on RYMV fitness, as plants showed similar symptom manifestation in rice plants to that of the wild-type FL5 infectious clone, especially in systemic tissues (Fig 5A) where it induced strong yellowing (Fig 5C). This suggests that Trp57 plays a minor role during RYMV infection. The further analysis of the related recombinant Trp29, Trp31 and Trp36 mutant proteins in *E. coli* (Appendix Table 1) indicated that recombinant P1^W29A^ and P1^W31A^ proteins were still soluble in *E. coli* (Appendix Fig S19, Appendix Table 2), the remaining insoluble fraction being probably due to protein overproduction. On the opposite, W36A mutation led to strong protein aggregation (Appendix Fig S19). We also observed that the W36A mutation provoked partial aggregation of both EGFP-P1^W36A^ and P1 ^W36A^-EGFP corresponding fusion proteins mostly in the plant cytosol, while W29A and W31A mutations did not affect P1 sub cellular patterns (Appendix Fig S17). These results indicate that Trp residues at positions 29, 31 and 36 are important for RYMV infectivity, either for their role in P1 oligomerization (Trp29 and Trp31) or in the maintenance of P1 structure in a close environment of the ZnF1 region (Trp36).

### C-terminal Zinc finger structure is required for P1 viral functions

According to our structural data, ZnF2 may ensure a central function in the folding of P1 C-terminal region. This region is also more sensitive to oxidation and less affine to zinc. We thus addressed the question of the biological role of ZnF2 during rice infection by RYMV RNAs through the use of FL5 RNAs C140H and C149H mutations generating artificial HHHC and HCHH ZnF2. Such mutations allow assessing redox behavior while keeping zinc coordination safe (Negi *et al*, 2004), and like C140S and C149S mutations, they do not alter P1 pattern in plants cells (Appendix Fig S17). Fig 6 shows that both FL5 infectious clones carrying C140H or C149H mutations were noninfectious in rice plants, as shown by the almost total lack of viral particles in inoculated and systemic tissues (Fig 6A), by the unaffected growth of the plants and their healthy leaf phenotypes (Fig 6B-C). This indicates that integrity of Cys140 and Cys149 is mandatory for RYMV infectivity, probably for ensuring ZnF2 structural lability and its redox-sensitive flexibility.

**Figure 6.**
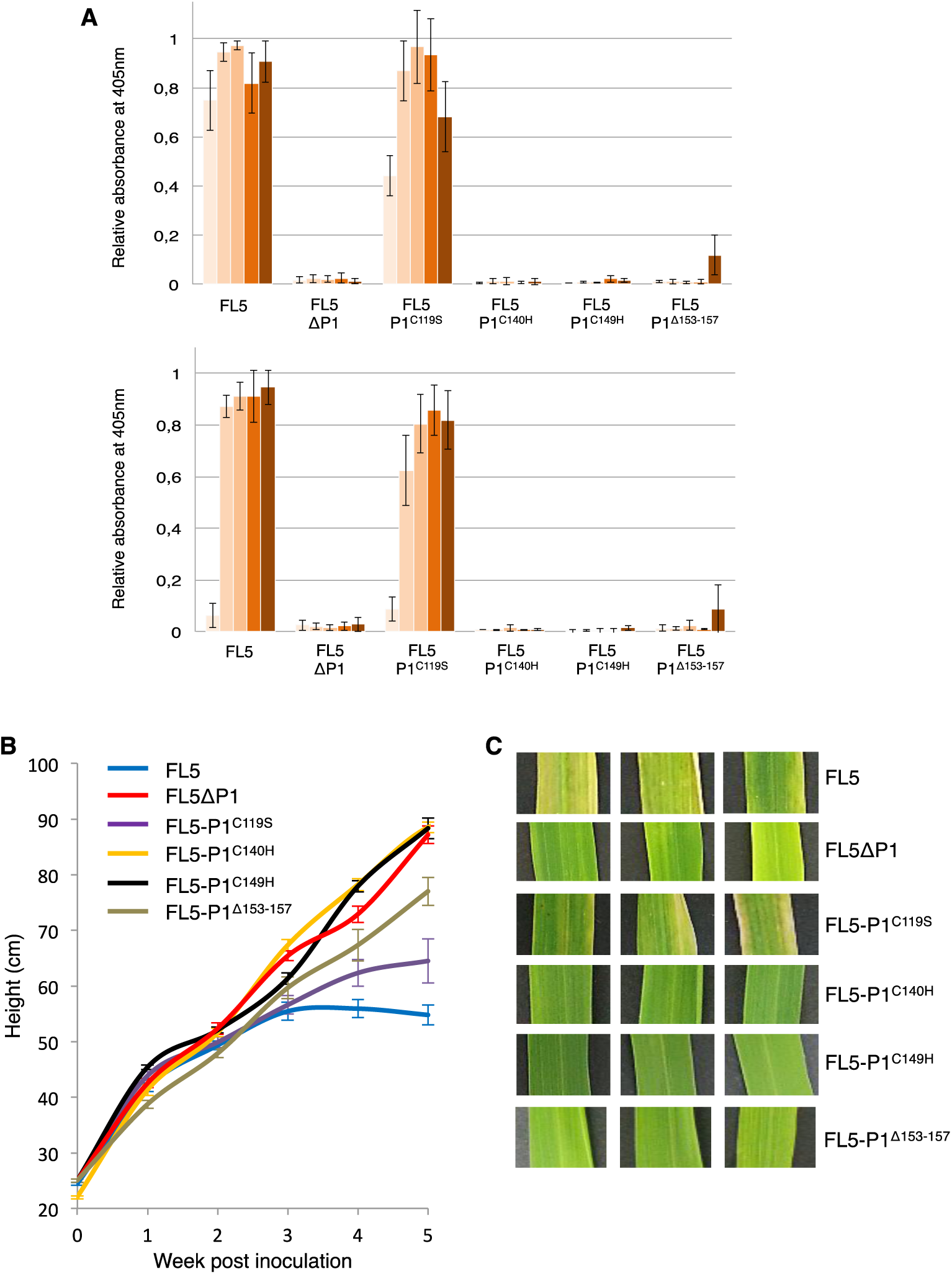
Incidence of mutations in P1 C-terminal region residues on RYMV infectivity in rice. **A** Consequences of cysteine and histidine residue mutations in P1, and of the KYNLF motif deletion (residues [153-157]) on RYMV particle accumulation in rice. The viral charge of rice plantlets infected by FL5-P1^H109A^, FL5-P1^C119S^, FL5-P1^C140H^, FL5-P1^C149H^, and FL5-P1^Δ153-157^ RNAs were recorded by DAS-ELISA on total proteins extracted from inoculated (upper panel) and systemic (lower panel) leaves collected every week during 5 weeks (light to dark bars). An antibody directed against the RYMV coat protein CP was used in DAS-ELISA for the detection of neo-synthesized viral particles. The original infectious clone FL5 (Brugidou *et al*, 1995) and its non-infectious version FL5ΔP1 (Appendix Fig S16) were used as positive and negative controls for infection and viral systemic spread, respectively. **B** Plant height (in cm) of inoculated plants recorded all along the 5-week infection process. **C** Yellowing symptom recording at the leaf surface of inoculated plants. Three samples of rice leaves are shown as representative of symptoms visible at the overall plant scale. Leaves were photographed 5 weeks post inoculation. **Data information:** An antibody directed against the RYMV coat protein CP was used in DAS-ELISA for the detection of neo-viral particles. Mock plants corresponding to rice plantlets inoculated with a buffer free of RYMV particles or FL5-derived RNAs also served as healthy plant controls for both height measurements and symptom recording. All data were obtained from 30 plants per treatment, in two independent inoculation assays.

We also challenged the importance of the strictly conserved C119 and the sole free Cysteine residue in P1 (Gillet *et al*., 2013) for the virus by creating a Cys119Ser mutation in FL5 to impair potential redox modifications of this residue. This mutation does not modify P1 solubility *in vitro* and in plant cells (Appendix Fig S10 and S17), or its zinc binding properties (Appendix Fig S10). Indeed, we found that infecting rice with FL5-P1^C119S^ did not provoke major differences in viral fitness, as this clone remained highly infectious in rice for all traits analyzed (Fig 6A-D). Cys119 residue is thus dispensable for P1 functions despite its structural accessibility.

Another P1 feature of interest is its highly hydrophobic and strongly conserved KYNLF sequence (NLF motif being strictly conserved in all P1s) ending the C-terminal part P1 (Appendix Fig S1). The KYNLF motif is also part of the aromatic basic pole identified in ZnF2. According to P1 structure (Fig 2 & 4), this region is largely solvent exposed and may be accessible for partner binding. Deleting this KYNLF stretch in P1 fused to EGFP did not alter P1 localization in rice cells (Appendix Fig S17), and a ^1D^NMR analysis of a corresponding recombinant P1^Δ153-157^ suggested that the KYNLF deletion does not generate protein instability or misfolding (Appendix Fig S20). But strikingly, the same KYNLF deletion in P1 ORF within FL5 strongly delayed the infectious capacities of the corresponding FL5-P1^Δ153-157^ viral RNAs, viral particles being detectable only from the 5^th^ week in both inoculated and systemic tissues (Fig 6A), with limited symptoms in term of plant growth retardation and yellowing (Fig 6B-C). This demonstrates that the five external amino acid stretch at C-terminal end of P1are essential for virus infectivity.

## DISCUSSION

Plant viruses count among the most damaging pathogens to plant growth, affecting not only plant fitness and crop yields, but also food quality. Determining the crystal structure of viral components mandatory for plant infection and spread in host tissues is a key step for a better understanding of the mechanisms underlying viral functions that in turn allows the development of adapted strategies for a better agronomy. Viral RNA silencing suppressors that specifically counteract the plant anti-viral RNAi pathway (Ding & Voinnet, 2007) are therefore particularly good candidates for which crystal structures are requested.

Here, we report the first complete 3D structure of an essential VSR protein from a member of the sobemovirus family, which infects the monocotyledonous model plant *Oryzae sativa*. To date, only few VSRs encoded by plant viruses have been crystallized (Baulcombe & Molnár, 2004; Ye *et al*, 2003; Chen *et al*, 2008; Yang *et al*, 2011) and of these, only one comes from a rice virus (Yang *et al*, 2011). Thus, the P1 structure described here contributes to fulfill an important gap of knowledge regarding VSRs. We show that P1 contains two sub-domains, harboring each a zinc finger but with completely different architectures. The N-terminal sub-domain folds as a compact sandwich composed by two repeats of 3 β-sheets, essentially maintained by a 4-Cys Zinc finger at its C-terminal extremity and by a short prominent helix at its N-terminal extremity. The C-terminal subdomain of P1 is composed by a long α-helix and a rigid part devoid of secondary structure, the latter elements being stabilized together by a second ZnF of a new type, namely His-Cys-His-Cys. The NMR analysis revealed that the individual structures of each sub-domain solved by X-Ray crystallography were retrieved in the whole protein, and behaved as semi-independent and flexible structural units in this context. Other VSR proteins were previously shown to be organized in modules, including ZnF recognized domains pointing to their involvement in diverse viral functions (López *et al*, 2000; Dong *et al*, 2003; Van Wezel *et al*, 2003; Trinks *et al*, 2005; Yang *et al*, 2007; Yambao *et al*, 2008; Valli *et al*, 2008; Chiba *et al*, 2013; Lukhovitskaya *et al*, 2013; Kenesi *et al*, 2017; Fujita *et al*, 2018). To our knowledge, we here report the first 3D structure revealing ZnF features within a plant VSR protein, with very few similarities with other proteins in the databases.

P1 protein is essential for RYMV systemic spread in rice tissues (Bonneau *et al*, 1998; Siré *et al*, 2008) and also plays an important role in viral replication despite being not strictly essential in this early step of RYMV infection (Bonneau *et al*, 1998; Nummert *et al*, 2017). Indeed, a mutation of P1 translation initiation codon rendering P1 undetectable in infected rice tissues almost abolished RYMV particles accumulation in infected and systemic leaves. Mutation on Cys64 and Cys95 had also been reported to have a negative impact on RYMV fitness (Siré *et al*, 2008). Our present work now brings a better understanding on how punctual mutations introduced at ZnF1 and ZnF2 on Cys residues impair viral infection, either by decreasing protein solubility and provoking protein misfolding and aggregation (Cys64S, Cys95S, Appendix fig S10 to S14 and S17), or by potentially decreasing P1 redox sensitivity, structural flexibility and/or ligand binding capacities (C140H, C149H, Fig 6 and Appendix Fig S17). Based on those results, the use of Cys mutants in different approaches revealed the necessity to conserve ZnF1 and ZnF2 integrity and structure for P1 functions. Similar results were observed for other viral ZnF proteins, e.g. is the replacement of His by Cys or ZnF depletion in NCp7 protein that resulted in a complete loss of HIV infectivity, as well as loss of interactions with viral protein partners such as Vpr and host partners RPL7 (Déméné *et al*, 1994; de Rocquigny *et al*, 1997; Mekdad *et al*, 2016). Another example is illustrated by AC2 or its homologous C2 protein from *mung bean yellow mosaic virus* and *tomato yellow leaf curl virus* respectively. Cys mutations involved in their ZnF decreased or totally abolished Zn affinity, VSR activity and transactivation properties (Dong *et al*, 2003; Trinks *et al*, 2005; van Wezel *et al*, 2002). However, the P1 protein presents the unusual particularity to possess two zinc-fingers of different types, an atypical feature never reported before to our knowledge. Of note, P1 C-terminal ZnF2 is at least hundred times more sensitive to Zn chelation and two times more sensitive to oxidation than ZnF1, indicating its higher chemical reactivity. These results are in accordance with their integration in P1 overall structure: ZnF1 functions as a closing cap of a compact beta-sheet sandwich, with limited access to the zinc atom, whereas ZnF2 bridges together structural elements that have minimal interacting secondary structures. Hence, zinc chelating and/or oxidation is easier in ZnF2, the Zn-release leading to unfolding of this domain. From all these data, we hypothesize that ZnF2 controls P1 redox flexibility. Different protein models have already been characterized for redox dependent Zn binding/release and subsequent 3D structure modifications that control *in vivo* functions. The bacterial Hsp33 possesses one ZnF composed of 4 Cys residues that undergoes disulfide bond formation and subsequent Zn release upon oxidative conditions (Barbirz *et al*, 2000), provoking the structural unfolding of Hsp33 C-terminal domain and the dimerization of the protein required to activate its chaperone function (Barbirz *et al*, 2000; Graumann *et al*, 2001; Winter *et al*, 2008; Ilbert *et al*, 2007). RsrA protein exemplifies a second mechanism of ZnF redox regulation. RsrA binds one Zn atom in its reduced state, this conformation being responsible for sigma factor sequestration. Oxidative stresses induce Zn release that accelerates sigma factor dissociation from RsrA, a process accompanied by disulfide bond formation and conformational changes by burying hydrophobic core necessary for sigma factor interaction. This overall process leads to RNA Pol activation through interaction with sigma factor, then RNA Pol-sigma factor complexes triggered transcription of redox balance genes (Paget *et al*, 2001; Li *et al*, 2003; Rajasekar *et al*, 2016). In P1 protein, the loss of ZnF2 structure upon zinc chelation/and or oxidation could serve as a new regulation level for the mutual orientation of the interaction motifs of ZnF1 and ZnF2 described in this study. We hypothesize that P1, in order to orchestrate such different functions during viral infection, has to fine-tune its structural conformation to modulate interactions with different partners.

Indeed, each subdomain was found to harbor a potential interaction motif with nucleic acids or proteins. This spot was constituted by a solvent-exposed helix for the N-terminal part, and a stretch of basic residues overlapping with aromatic residues for the C-terminal part. The importance of these motifs is highlighted by the fact that deletion of the solvent exposed helix at N-terminal extremity (P1^Δ12-21^) or of the five external amino acids at C-terminal end (P1^Δ153-157^) leads to a complete loss of virion infectivity *in planta* (Fig 6A). The relative flexibility of both sub-domains therefore offers a great potential in differently aligning these interaction motifs, multiplying the different regulation levels as does the oxidative stress by ZnF2 unfolding. Unexpectedly, an additional level of regulation is provided by the oligomerization properties of P1. Our structural data show that P1_1-100 crystal structure presents two potential dimeric interfaces but interaction surfaces and calculated binding energies suggest that those dimers are labile (Chen *et al*, 2013). Nevertheless, dimerization was confirmed *in vivo* by yeast two-hybrid for P1_1-100 and *in vitro* by SEC-MALLS experiments for P1_1-100 and P1 full length. Moreover, mutations of Trp29 and Trp31, which are involved in dimerization interface (asymmetric unit), negatively impact systemic spreading of the virus with a strong delay or a complete loss of virus accumulation in systemic leaves respectively. This result highlighted the biological relevance of P1 dimeric interface observed in the asymmetric unit. Similarly to P1, other known VSR structures were obtained in dimeric conformation such as p19, NS10, B2 and 2b (Vargason *et al*, 2003; Ye *et al*, 2003; Baulcombe & Molnár, 2004; Chao *et al*, 2005; Chen *et al*, 2008; Yang *et al*, 2011) or octameric form for p21 (Ye & Patel, 2005). The multimeric state of those proteins were always associated with dsRNA binding activity required for VSR functions, especially in p19 structure presenting a WxxW motif required for siRNA binding (Vargason *et al*, 2003; Ye *et al*, 2003). This motif is also found in the *Rice Dwarf Phytoreovirus* Pns10 protein, and mutation of this motif alters its VSR activity (Ren *et al*, 2010). In P1, Trp29 and Trp31 form a related motif W_29_xW_31_, however P1 seems to behave differently compare to other VSRs with known structures, as all attempt to observe RNA binding activities were unsuccessful in our experimental conditions. Therefore, we hypothesize that the mechanism of interference with 21 nts and 24 nts siRNA previously described seems to occur in an RNA binding independent manner not yet understood. Moreover we cannot exclude that Trp29 and Trp31 residues forming a WxW motif are also involved in cellular partner’s recruitment necessary for viral systemic infection. Taking into account that the phenotype of RYMV carrying Trp29 or Trp31 mutations relies on systemic movement reduction or deficiency, we hypothesize that dimerization properties and/or partners recruitment involving this motif play a crucial role in viral movement, which might be correlated or uncoupled with VSR properties and viral replication. We also hypothesize that those two mutants might be used to decipher the molecular mechanism of P1 involved in virus movement and/or RNAi evasion. This question has to be further addressed to fully decipher P1 functions. Regarding the C-terminal region of P1, the extreme C-terminal part presents a patch of highly conserved solvent-exposed lysine and arginine residues (Arg150 and Lys153) which only tolerates Arg to Lys or Lys to Arg replacement with the other positively charged residue, overlapping with a stretch of aromatic residues (Phe151, Tyr154 and Phe157) which tolerates no or only replacement with aromatic residues (Tyr154 excepted, for which a replacement with a cysteine is also observed in one sequence among the 51 inspected) (Appendix Fig S1). This region may constitute potential anchoring sites for interaction with nucleic acids and hydrophobic partners, respectively.

On the other hand, some VSRs such as p38 act in homodimeric or higher multimeric forms to mimic GW repeated motifs involved in Ago1 interactions (Azevedo *et al*, 2010; El-Shami *et al*, 2007; Till *et al*, 2007). Regarding P1 organization, Trp57 is involved in P1 dimeric interface in the crystallographic unit and also forming with Gly56 a potential GW motif. This GW motif might form a GW repeat through P1 dimerization for interacting with Ago proteins. However, we could not obtain experimental evidence for the biological relevance of this GW motif or of the dimeric interface observed in the crystallographic unit, as the mutation of Trp57 did not alter the infectivity profile, a feature that is in line with its low conservation among all P1 isolates (Sérémé *et al*, 2014 and Appendix figure S1). Punctual mutations in this second potential dimeric interface have to be further investigated.

Altogether our results lead to a better understanding of P1, a key protein for the RYMV virus, and a unique protein in the database with original features. They notably point out to the multiple levels of regulation of its structure, i.e. flexibility, oxidation and oligomerization and their impact on its function during the viral cycle. Future identification of P1 targets in rice will bring key information on the mechanisms by which P1 interferes with plant anti-viral functions, and assist the design of appropriate strategies to counteract RYMV. Very recently, putative 22 nts miRNAs encoded by the genome of *O. sativa* that can potentially target the RYMV genome have been identified using a consensus of four algorithms (Jabbar *et al*, 2019). Whether such sRNAs could be potential P1 targets for counteracting plant antiviral defenses remains to be evaluated. We have also initiated a deep search of rice proteins as potential P1 interactors and identified host proteins suspected to assist P1 folding. The availability of P1 3D structure coupled to our capacity to challenge the importance of key residues both at biochemical, structural and biological levels opens huge perspectives for the study of P1/host partner complexes and their role during rice infection by RYMV. They also open huge perspectives towards the understanding on how P1 structural properties sustain P1 sequence diversity among the numerous RYMV isolates identified so far in the African continent and neighboring islands (Rakotomalala *et al*, 2019) and much beyond among the sobemovirus gender.

## MATERIALS AND METHODS

### Plant culture and analyses

The rice *Oryza sativa*, cultivar Indica, Var. *IR64* was selected for its high sensitivity to the RYMV (Ndjiondjop *et al*, 1999). Rice plants were grown in confined greenhouse under 12 h light/12 h dark at 28°C and at a relative humidity of 75%, and were watered daily. For rice infection assays, series of 30 two week-old rice plants (three-leaf stage) were used and cultivated after infection under growth conditions described above. Inoculations of viral RNAs were performed after in vitro RNA transcription according to (Brugidou *et al*, 1995). All procedures related to RYMV isolates, infection and detection of viral particles in rice are described in details in the Appendix Materials and methods section. Both inoculated and systemic leaves were collected at different times post-infection (from 7 to 28 dpi) and stored in liquid N2 until use. Protoplasts from 15-day old etiolated rice leaves were prepared according to Chen *et al* (2009) and transfected according to Zhang *et al* (2011). All experiments related to confocal microscopy are described in the Appendix Materials and Methods section.

### Constructs

All constructs expressing P1 and primers used in this study are listed in the Appendix Table 1. For cloning by restriction/ligation, DNA was amplified using the Phusion High-Fidelity enzyme (Finnzymes) and restriction site-containing primers. DNAs were subsequently cloned into destination vectors after digestion by corresponding restriction enzymes (New England Biolabs) and ligation with T4 DNA ligase (Promega). Mutations required for domain excision and/or punctual changes within P1 nucleotidic sequence were introduced in the P1 open reading frame using the QuikChange® II XL Site-Directed mutagenesis kit (Agilent-Stratagene) and related primers (Appendix Table 1). All constructs and mutations created for this study were systematically checked by sequencing on both strands.

### Production and purification of recombinant proteins

For the production of P1 recombinant proteins, a full-length P1 nucleotide sequence originated from the RYMV Tz3 isolate was already cloned in a pET3b vector (Novagen) between *NdeI* and *BamHI* restriction sites to produce the P1 referent protein (Gillet *et al*, 2013). Additional truncated P1 proteins (P1_1-100, P1_50-100, P1_50-157 and P1_102-157, amino acid numbering) were generated similarly in the pET3b vector. These four constructs were used to generate pET3b collections of mutant P1 coding sequences by site-directed mutagenesis, including the pET3b. P1Δ153-157 construct carrying an early stop codon in place of Lys153 (Appendix Table 1).

All recombinant proteins were produced in the *E. coli* strain BL21-DE3 cells (Novagen) by induction with 0.4 mM IPTG at an OD_600_ of 0.6 and purified according to (Gillet *et al*, 2013) with modifications depend on P1 mutants (Appendix Table 2). Protein concentrations were determined by measuring absorbance at 280 nm using the calculated value of extinction coefficient for each different P1 variant with a Nanodrop spectrophotometer (ThermoScientific).

### Far-UV circular dichroism spectroscopy

CD spectroscopy analyses of reduced or oxidized P1 (12 µM) and its sub regions (25 µM) were as described for the wild type P1 protein (Gillet *et al*, 2013). For all experiments, molar ellipticity was calculated as described by Kelly and co-workers (Kelly *et al*, 2005) on an average of three independent measurements.

### SDS-PAGE analyses and immunoblotting

For bacterial fraction analyses, cells were collected by centrifugation in a buffer containing 50 mM Tris-HCl pH 8, 2 mM DTT, then sonicated and separated into insoluble and soluble fractions. Extraction of insoluble protein fraction were performed using Tris-HCl 50 mM pH 8 urea 6M under shaking conditions until insoluble pellets are fully resuspended (≈30 min), then bacterial debris remaining insoluble are eliminated by centrifugation. Renaturation of insoluble proteins were performed by dialysis overnight at 4°C under gentle shaking in Tris-HCl 50 mM pH8. Protein samples were treated with Laemmli buffer (40 mM Tris-HCl, pH 6.8, 1% SDS, 50 mM DTT, 7.5% glycerol, and 0.003% bromophenol blue) and analyzed by SDS-PAGE in denaturing conditions using 15% to 18% acrylamide gels (37.5:1 ratio acrylamide:bis-acrylamide, 40%) depending on P1 mutants. Purification of P1 mutants from bacterial extracts was performed according to (Gillet *et al*, 2013). Purified P1 protein (25 μM) and its derived truncations (50 μM) were analyzed similarly or in non-reducing conditions without DTT according to (Gillet *et al*, 2013). Redox treatments by different concentrations of H_2_0_2_ or DTT were applied to purified proteins prior to SDS-PAGE analyses either at 20 °C for 45 min and in different buffers depending on the experiment (see figure legends).

On-gel zinc detection in proteins after SDS-PAGE was performed prior to Coomassie blue staining using the zinc-complexing probe PAR (4-(2-Pyridylazo)resorcinol, Fluka) (Lee & Helmann, 2006).

Immunoblotting was performed after protein transfer onto Hybond-P membranes (GE Healthcare, RPN303F). After blocking membrane nonspecific binding sites in TBS (20 mM Tris, 75 mM NaCl, and 2.5mM MgCl_2_, pH 7.6, 5% milk powder), proteins were subjected to primary antibody recognition using an anti-P1 antibody raised in rabbit (Siré *et al*, 2008) and purified by Affi-Gel 15 immnunoaffinity chromatography (BIO-RAD) using the recombinant P1 protein as the ligand (used at a 1:1000 dilution). A peroxidase-coupled secondary antibody (anti-rabbit IgG; Immunopure Pierce) was used at a dilution of 1:20000 for primary antibody recognition. The presence of P1 proteins was revealed using the ECL Plus Western Blotting Detection System (GE Healthcare).

### Determination of P1_1-100 zinc binding constant

The affinity of different regions of P1 for zinc ions was determined according to (Jakob *et al*, 2000) with slight modifications. Purified P1_1-100 (40 µM) was first reconstituted after treatment by 2 mM DTT and 300 µM ZnSO4, then DTT and unbound zinc were removed using PD10 gel filtration columns (GE Heathlcare) equilibrated with a 40 mM HEPES-KOH, pH 7.5 buffer. Equilibrium between P1_1-100 and zinc was further determined by treating P1_1-100 overnight with increasing concentrations (37 µM to 5 mM) of the Zn chelator TPEN *(N,N,N′,N′*-tetrakis(2-pyridyl-methyl)ethylenediamine) at room temperature in a 50 mM HEPES-KOH buffer (pH7.5) supplemented with 1 mM DTT. TPEN complexed with Zn, free TPEN and free Zn were further removed by gel filtration on PD MidiTrap G-10 (GE Healthcare) equilibrated with 40 mM of the metal-free HEPES-KOH, pH 7.5 buffer. To determine the amount of Zn still bounded to proteins, we performed a PAR/PMPS assay as follows: 80µl of PAR probe were added to 720µl of protein to a final concentration of 100µM. Then we used 1 mM PMPS (*p*-hydroxymercuriphenylsulfonic acid) which forms mercaptide bond with Cys residues and thus induces the release of the remaining Zn still bound to protein. Zn released from P1 directly reacts with PAR probe to form Zn(PAR)_2_ complexes colored in far red. We used spectrophotometry absorbance analyses at A_500nm_ and the GraphPad Prism^R^ software for non-linear regression and graphic representation to determine the amount of Zn released by PMPS addition. Values from two independent experiments were used for non-linear regression. We used the following equation for Zn affinity determination according to Jakob *et al*, 2000:

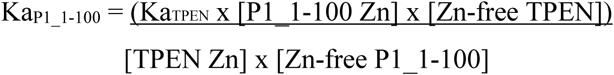

Where Ka_TPEN_ = 10^16^ M^-1^ (Anderegg *et al*, 1977), the TPEN concentrations necessary to release 50% of Zn atoms in P1_1-100 is determined graphically ([TPEN_Zn50%_] = 63 × 10^−6^ M). [P1_1-100 Zn] and [Zn-free P1_1-100] correspond to the concentrations of proteins with or without Zn respectively. [P1_1-100 Zn] = [Zn-free P1_1-100] = 20 × 10^−6^ M. [Zn-free TPEN] and [TPEN Zn] correspond to the concentrations of free TPEN or in complex with Zn atom. [Zn-free TPEN] = [TPEN_Zn50%_] - [Zn-free P1_1-100].

### Native mass spectrometry

Purified rP1 was buffer exchanged against 50 mM ammonium acetate, pH 8.0, using microcentrifuge gel-filtration columns (Zeba 0.5 mL, ThermoScientific, France) prior to MS analyses. Native MS experiments were performed on a high resolution electrospray quadrupole time-of-flight mass spectrometer (Synapt G2, Waters, Manchester, UK) equipped with an automated chip-based nanoelectrospray device (TriversaNanomate, Advion Biosciences, Ithaca, NY). Calibration was performed with the multiply charged ions produced by 2 μM horse heart myoglobin diluted in 1:1 (v:v) acetonitrile:water with 1% formic acid. Molecular mass, integrity, and homogeneity of rP1 were first checked in denaturing conditions by diluting rP1 down to 2 μM in 1:1 (v:v) acetonitrile:water acidified with 1% formic acid.

Analyses in native conditions were carried out using 5 μM to 10 μM of P1 and mutants in 50 mM ammonium acetate, pH 8.0. Optimal acceleration voltages applied on the sample cone (Vc) and a well-adapted pressure in the interface (Pi) was used to preserve weak noncovalent interactions in the gas phase. MS data were acquired and processed using MassLynx(tm) 4.1 (Waters). Peak intensities of the main charge states of the different species were used to calculate ratios of each detected species.

For EDTA treatment, rP1 (5 µM) was incubated overnight at 4 °C with 40 ou 200 equivalent EDTA and 1 mM DTT. For dynamic monitoring of zinc reversibility, EDTA-treated rP1 was incubated with 100 μM zinc acetate addition.

### X-ray crystallography

The purified P1_1-100 and P1_102-157 proteins were concentrated to 10 mg/ml mL and 50 mg/mL respectively in 20 mM Tris-HCl pH 8, 75 mM NaCl, 1 mM DTT and 300 µM ZnCl_2_. Crystallization trials were performed at 20°C using the hanging-drop vapor-diffusion method in 96 microplates and nano X8 (Cartesian) robot with 100 nl of protein mixed with 100 nl of reservoir. After 3 weeks, small crystals of P1_1-100 were obtained from condition 35 of the Classic Suite II kit (Qiagen) containing 0.1 M HEPES pH 7, 1 M Ammonium sulfate, and 0.5 %(w/v) PEG 8000. After further optimization, diffracting crystals were obtained from 0.1 M MES pH 6, 0.5 M Ammonium sulfate and 0.5 % PEG 8000 using the hanging-drop vapor-diffusion method. Crystals were cryo-protected by flash soaking in the same solution containing 20 % of 1-4 Butandiol (Sigma) before flash freezing in liquid N_2_. After 5 weeks, diffracting crystals of P1_102-157 were obtained at 4°C from condition 7 of the Classic kit (Qiagen) containing 0.1 M tri-Sodium citrate pH 5,6, 20 %(v/v) Isopropanol and 20 %(w/v) PEG 4000. Crystals were cryo-protected by flash freezing in liquid N_2_.

### X-Ray data collection, processing and structure determination

Crystal diffraction datasets were collected at the European Synchrotron Radiation Facility (ESRF, Grenoble) at beam lines ID23-1 and ID29 using a Pixel detector (Pilatus 6M) and processed by XDS (Kabsch, 2010) and Scala (Evans, 2006) from the CCP4 programs suite (Winn *et al*, 2011).

P1_1-100 crystals belong to the F4_1_32 space group and contain two molecules in the asymmetric unit. Their structure was determined to 2.10 Å by the single wavelength anomalous diffraction (SAD) method using HKL2MAP (Sheldrick, 2010) and ARP/WARP (Langer *et al*, 2008; Murshudov *et al*, 2011) from the CCP4 programs suite.

P1_102-157 crystals belong to the P12_1_1 space group and contain two molecules in the asymmetric unit. P1_102-157 structure was determined to 2.10 Å by the single wavelength anomalous diffraction (SAD) method using AUTOSOL from PHENIX (Adams *et al*, 2010) and ARP/WARP and P1_102-157 phasing data set. A higher resolution structure was determined using a higher resolution dataset (P1_102-157 at 1,98 Å) and molecular replacement with PHASER (McCoy *et al*, 2007).

After model building using Coot (Emsley *et al*, 2010) and refinement by REFINE from PHENIX, final structures exhibited an R(%) / R(%)_free_ of 0,185 / 0,214 for P1_1-100 and 0,193 / 0,223 for P1_102-157. Final refinement statistics for the structures are listed in Table 2. Figures were generated using PyMol (http://www.pymol.org/).

The atomic coordinates and structure factors of P1_1-100, as well as the P1_102-157, have been deposited in the Protein Data Bank with accession numbers 6TY0 and 6TY2, respectively.

### Nuclear magnetic resonance analyses and structure calculations

Production of proteins for NMR studies followed the protocol detailed in Yang *et al* (2019) and in Appendix Table 2. Briefly, ^15^N-edited HSQC-TOCSY and –NOESY experiments were used to assign resonances of the short constructs. The assignment of full P1_1-157 was deduced from these experiments, except the linker region (residue 98-103). ^15^N/^13^C/^1^H triple resonance experiments on the full length protein confirmed NMR resonance assignment for the N- and C-terminal domains and enabled the identification of chemical shifts of the linker region. Chemical shifts have been deposited in the BioMagResBank (http://www.brmb.wisc.edu) with accession number 27880.

Residual dipolar couplings were measured as differences of ^1^J_HN_ couplings between an aligned sample with a 14 mg/mL Pf1 phage suspension (Asla^Biotech^) and an isotropic sample of ^15^N labelled P1_1-157. ^1^J_HN_ couplings were measured as doubles of chemical shift differences in the ^15^N dimension of the ^15^N-^1^H correlation peaks of TROSY-HSQC spectra pair (Kontaxis *et al*, 2000), recorded each with 128 complex points inF1, and subsequently zero-filled to 256 complex points using linear forwards prediction. Extraction of the alignment tensor parameters was performed with the Module software (Dosset *et al*, 2001).

^15^N relaxation datasets were obtained according to published pulse sequences incorporating a heating compensation scheme between T1 and T2 experiments (Farrow *et al*, 1994). Experiments were recorded at 20°, with a recycle delay of 3 s. Delay times were 1 ms, 100 ms, 600 ms, 900 ms, 1200 ms and 1600 ms for the T1 experiments and 15.8, 31.6, 47.5 and 63.4 ms for the T2 experiments. Compensatory refocusing periods were introduced in the T1 and T2 experiments so that the spins experience an identical heating in all relaxation experiments (Mulder *et al*, 1999). The protein was slowly aggregating over times, leading to a slightly accelerated decay of magnetization, and resulting in an 0.94-0.96 scaling factor between two successive T2 or T1 experiments. Fits of the relaxation times were performed after a rescaling of the corresponding intensities. Estimation of the diffusion tensor was done with the Tensor software from T1/T2 ratios (Dosset *et al*, 2000).

Derivation of the mean structure of the whole protein was done by conjoined rigid-body/torsion angle/Cartesian simulated annealing driven by NMR restraints, according to the protocol developed by Clore and co-workers (Schwieters *et al*, 2010). The calculations were carried out in Xplor-NIH (Schwieters & Clore, 2001; Schwieters *et al*, 2006). The residues of the 1-96 fragment were held fixed in space, whereas the 105-157 fragment was treated as a rigid body. After a randomization step of backbone dihedral angles, atoms of the linker region connecting both zinc fingers (residues 98-104) were given first Cartesian and then torsional degrees of freedom, under residual couplings, restraints derived from ^1^H-^1^H Noes, and ϕ/Ψ restraints derived by TALOS (Shen & Bax, 2013) from HN, N, Cα, Cβ and C chemical shifts, in addition to covalent and geometrical energetic terms. The coordinates and experimental data have been deposited in the PDB, accession number 6XV2.

### Supplementary materials

All tools and methods for Y2H and BiFC experiments, as well as additional information concerning RYMV isolates, infection and detection in rice assays are described in the Appendix Materials and Methods section.

## ACKNOWLEDGEMENTS

We are grateful to Pr Grishin (University of Texas, USA) and Srikrishna Subramania (CSIR-Institute of Microbial Technology, India) for helpful discussion and expertise. We thank the ID23-1 and ID29 beams lines scientists at European Synchrotron Radiation Facility (Grenoble) for their excellent support during data collections. We also thank Caroline Y.I. Hsing and Lin-Yun Kuang (IPMB Institute, Academia Sinica, Taipei, Taiwan) for training V. P. to plant protoplasts isolation, and C. Alcon (Montpellier RIO-Imaging and PHIV platforms at http://www.mri.cnrs.fr/, Montpellier-France) for confocal microscopy assistance. V. P. and F. V. are also grateful to the Franco-Taiwanese Partenariat Hubert Curien Orchid program n°29354SD, to the Taiwanese Research and Practical Training Program, and to the Gaïa PhD School of Montpellier University, France, for their financial supports. The CBS is a GIS-IBIsA platform and is member of the French Infrastructure for Integrated Structural Biology (FRISBI), a French infrastructure supported by the National Research Agency (ANR-10-INBS-05). V. P. and F-X. G. were awarded with PhD Internships from the French Ministry of Education and Scientific Research.

## AUTHOR CONTRIBUTIONS

VP, SC, HD & FV designed the research. VP & FV designed and conducted DNA cloning & site-directed mutagenesis programs. VP, FXG & FV designed biochemical procedures, VP performed all recombinant protein processes, VP, FA & EL performed in vitro assays, and FV performed Y2H assays. GT & SC designed and performed mass spectrometry experiments. FH, YY, VP, FXG & HD designed and conducted crystallographic and NMR experiments. VP, JHK & FV collected plant samples, and conducted infection processes, cellular experiments & confocal microscopy analyses. VP, FH, GT, YY, JHK, SC, HD & FV analyzed the data. VP, FH, YY, FXG, CB, HD & FV reviewed the manuscript. VP, HD & FV wrote the manuscript.

