## Appendix text and figures for "An original structural fold underlies the multitask P1, a silencing suppressor encoded by the Rice yellow mottle virus"

### APPENDIX TEXTS

### SUPPLEMENTARY RESULTS

#### Delineation of P1 zinc binding domains, in more details

The design of recombinant P1 sub-regions required for biochemical and structural purposes was mostly based on *in silico* analyses. Protein alignments performed against the databases suggested that the [50-100] region may constitute a potential zinc-binding domain considering its similarity of its CXXC microdomains with that of few zinc binding proteins (Gillet *et al*, 2013). The strict conservation of cysteine/histidine residues at the C-terminal region of P1 together with the demonstration that P1 binds two zinc atoms (Gillet *et al*, 2013) also suggested that this region could constitute another Zn-binding region albeit being hardly predictable. We thus took into consideration these predictions to generate four different truncated proteins to challenge the hypothesis that they could bind one (regions [1-100], [50-100] and [102-157]) or two (region [50-157]) zinc atoms. The PredictProtein server (Rost *et al*, 2004) was also used to assist the design of P1 regions to be subcloned in order to preserve predicted alpha helices and beta strands (Fig 1). All resized recombinant P1 regions were efficiently produced in the *Escherichia coli* BL21 strain (Appendix Fig S2) and appeared properly folded after purification (Appendix Fig S3).

#### P1 N- and C-terminal regions exhibit different structures and zinc binding geometries in more details

To support the structural data obtained on the different P1 sub-domain, we first produced a set of full-length P1 monocysteinic and monohistidinic mutant proteins to evaluate their zinc binding properties (Appendix Tables 1 & 2). All mutants assayed were efficiently produced in *E. coli* (Appendix Fig S10) but several mutants carrying a mutation in the P1 region [50-100] ($P1^{C64S}$, $P1^{C67S}$, $P1^{C92S}$, $P1^{C95S}$) or in the P1 Cter region [102-157] ($P1^{H109A}$) were insoluble despite co-localizing with zinc (Appendix Fig S10A-B, upper and lower panels), suggesting a major role of these residues in P1 folding. Strikingly, further analyses of these mutants using native ESI-MS confirmed that they all bounded two zinc atoms despite single Cys and His mutations except $P1^{H109A}$ that remained improper for ESI-MS analyses (Appendix Fig S10C & S11). We pushed on our strategy by generating single and double Cys and His mutations directly into the shorter P1_50-100 and in P1_102-157 recombinant proteins. Regarding the [50-100] central region (Appendix Fig S12 & S13), all the mutants tested were insoluble ($P1\_50\text{-}100^{C64S.C67S}$ even remained not producible) but were successfully renatured and purified (Appendix Fig S12A). Among them, $P1\_50\text{-}100^{H70A}$, $P1\_50\text{-}100^{H87A}$ and $P1\_50\text{-}100^{H70A.H87A}$ proteins remained able to bind one zinc atom despite mutations as indicated by native ESI-MS analyses (Appendix Fig S12A-C & S13) showing that His70 and His87 residue are not involved in zinc coordination. On the other hand, Cys-to-Ser mutations in P1_50-100 provoked different behaviors. $P1\_50\text{-}100^{C64S.C67S}$ could never be produced in *E. coli*, while $P1\_50\text{-}100^{C64S}$ and P1_50-

$100^{C67S}$ single mutant proteins could not co-localized anymore with zinc in gels and remained improper for native ESI-MS analyses (Appendix Fig S12A-C and S13) probably due to incorrect refolding after renaturation. A similar result was obtained for the double mutant protein P1_50-$100^{C92S.C95S}$, although single C92S and C95S mutations were found to have a minor effect of zinc release (Appendix Fig S12A-C and S13). Altogether with our P1 structural analysis, these data allowed us to propose that ZnF1 motif in P1 central region is [$C64x_2C67x_{23}C92x_2C95$], affiliated to the CCCC-type ZnF superfamily.

Regarding P1 C-terminal region, 5 mutations (H109A, C119S, C140S, H145A, and C149S) were introduced in P1_102-157 and independently analyzed. We first found that the shorter P1_102-$157^{C119S}$ protein still abundantly accumulated in soluble fractions (Appendix Fig 12B, upper panel), and strongly co-localized with zinc in gel fractions despite mutation (Appendix Fig S12B, lower panel). This was consistent with our previous data showing that Cys119 is the sole free cysteine residue in P1 (Gillet *et al*, 2013), and reinforced the hypothesis that Cys119 is not required for zinc coordination at P1 Cter region. All other mutant proteins accumulated at very low levels in soluble fractions with strong reduction of zinc binding (Appendix Fig 12B, lower panel), and none of them could be purified for further ESI-MS analyses. Altogether, these data supported the evidence that His109, Cys140, His145, and Cys149, but not Cys119, are essential for P1 Cter region folding and constitute the second [$H109x_{30}C140x_4His145x_3Cys149$] ZnF2 affiliated to a HCHC ZnF superfamily.

**P1 oligomerization is essential for viral spread in rice, in more details**

To evaluate the incidence of mutations in P1 during viral infections, we selected the Ivorian Cost RYMV Isolate (*Cia*) and its derived infectious RNA clone, the so-called FL5 (Brugidou *et al*, 1995) as this RYMV strain provokes strong infection symptoms in the *O. sativa IR64* rice variety. Compared to healthy plants (Appendix Fig S16A, left panel), RYMV-infected plants showed wild spread yellowing symptoms at the leaf surface and strong decrease of growth (Appendix Fig S16A-B and D). RNAs originated from FL5 transcription provoked similar symptoms (Appendix Fig S16A-B, right panel and S16C, upper panel), with a slight decrease in the amount of RYMV particles accumulated in infected tissues (Appendix Fig S16E). A mutation of P1 ATG start codon in FL5 sequence (FL5∆P1 in all panels) that renders the protein undetectable using a purified anti-P1 antibody (Appendix Fig S16F) abolished all viral symptoms.

**Appendix Materials and methods**

**Binary yeast-two hybrid assays**

The yeast strain YRG2 (Stratagene, Agilent) was used to evaluate P1 domain dimerization properties. P1 regions of interest (P1_1-100 and P1_102-157) were excised by *NdeI*/*BamHI* restriction from the pET3b collection (Appendix Table 1) and ligated at compatible sites in the pGAD.T7 and pGBK.T7 yeast two-hybrid vectors (Clontech). pGAD and pGBK constructs were co-introduced in yeast cells and assayed for binary yeast-two hybrid interactions as described previously (Vignols *et al*, 2005). All constructs were checked by sequencing.

**RYMV isolates, infection and detection in rice**

Rice plants were infected either using a RYMV inoculum or with synthetic viral RNAs, both originated from the *Ivorian Cost* RYMV isolate (Yassi *et al*, 1994). RYMV inoculum was obtained from infected leaves and mechanically inoculated to rice plants using carborundum (Fisher Scientific) as described earlier (Konaté *et al*, 1997). Full length synthetic RYMV RNAs were obtained with the T7 RNA polymerase (Promega) using as DNA template the linearized pUC19-based infectious clone FL5 (Brugidou *et al*, 1995) carrying the entire RYMV *Cia* genome under the control of the bacteriophage T7 promoter. FL5 was also used as DNA template for mutagenesis to produce synthetic RYMV RNAs carrying mutations in the P1 open reading frame (listed in Appendix Table 1). Rice plant infections by synthetic viral RNAs were performed according to Brugidou *et al*, (1995).

For all infected rice plants, total RNA was extracted from one-week-old leaves (100 mg pieces) using the RNeasy Plant Mini kit (Qiagen) and subjected to reverse transcription using the RevertAid First Strand cDNA Synthesis Kit (Fisher Scientific). Occurrence of the viral RNA was checked by PCR among cDNAs using the GoTaq polymerase (Promega) and RYMV_19F ($^{5'}$CTCTACGACTATGCTGACACC$^{3'}$) and RYMV_20R ($^{5'}$CTCCCCCACCCATCCCGAGA$^{3'}$) primers (Appendix Table 1). To check the non-reversion of mutations designed in the P1 coding region within FL5-derived RNA during infection, cDNAs from 3 weeks infected plants were also amplified with the Phusion High-Fidelity enzyme using RYMV_1F ($^{5'}$ACAATTGAAGCTAGGAAAGGAGC$^{3'}$) and RYMV_4R ($^{5'}$GGTCGCTTTCTCACTCGCACC$^{3'}$) primers (Appendix Table 1) surrounding P1 open reading frame in FL5 and systematically checked by sequencing.

Detection of viral particles in infected plants was done by Das-ELISA according to Konaté *et al* (1997) with slight modifications. Leaf samples (1 g) were ground with 10 ml of PBS containing 0.05% Tween 20 (PBS-T) and 2% polyvinylpyrolidone (PVP). ELISA plates (Nunc-Immuno™ MicroWell™ 96-well solid plates, Thermo Scientific) were coated with purified uncoupled rabbit IgGs directed against $RYMV_{MG}$ particles (an RYMV isolate from Madagascar) diluted at 1 µg.mL$^{-1}$ in a carbonate coating buffer (15 mM $Na_2CO_3$m, 34 mM $NaHCO_3$, pH 9,6, 100 µL/well) for 2h at 37°C. After washing for 10 min with a TBS-T buffer (20 mM Tris-HCl pH 7.5, 150 mM NaCl, 0,05% Tween20, v/v), non occupied sites in wells were blocked for 1h at 37°C using TBS-T buffer containing 5% milk powder (200 µL/well). After a second plate wash, 100 µl of leaf extracts diluted to 1/100 were delivered to individual wells and incubated for 16 h at 4°C. After washing, secondary antibodies against $RYMV_{MG}$ conjugated to alkaline phophatase and diluted in TBS-T at a 1/1000 ratio were added to wells (100 µL/well) and incubated 2h at 37°C. The disodium 4-nitrophenyl phosphate hexahydrate (PNPP) substrate for alkaline phosphatase was diluted at 1 mg.mL$^{-1}$ in diethanolamine buffer (1 M diethanolamine pH 9,8, 0,5 mM $MgCl_2$) and added to wells (100 µL/well) after ultimate plate washing. Chromogenic reactions developed during 1h at dark, and optical densities (O.D) were measured at 405 nm in a V-1200 UVisco spectrophotometer. Data normalization was performed using data from wells treated with particle-free buffer as the reference values.

For P1 immunodetection analyses by western blots, a purified anti-P1 antibody from Siré *et al* (2008) was used (see plain text, Materials and Methods section).

**Transient expression assays and confocal microscopy**

The pEZS-NL and pEZS-CL vectors (http://deepgreen.stanford.edu/) carrying the CaMV 35S-MCS-$(Ala)_{10}$-EGFP (pEZS-NL vector) or the CaMV 35S-EGFP-$(Ala)_{10}$-MCS cassettes (pEZS-CL vector), respectively, were selected to express P1-EGFP and EGFP-P1 fusion proteins and related variants in rice cells. P1-encoding DNA sequences of interest were amplified from the pET3b collection and sub-cloned between *SalI* and *BamHI* restriction sites to give rise to pEZS.P1-EGFP and pEZS.EGFP-P1 constructs (Appendix Table 1). pEZS.P1-EGFP and pEZS.EGFP-P1 constructs were subsequently used as templates to generate two collections of mutant P1 in frame with the EGFP protein by site-directed mutagenesis (Appendix Table 1).

Localization of EGFP-P1 and P1-EGFP fusion proteins and related variants in rice protoplasts was performed using a maximum of 10 µg total DNA per transfection event. EGFP fluorescence (Cormack *et al*, 1996) was recorded 24 hours post transfection after excitation with an argon laser at 488 nm (fluorescence detection range: 500-550 nm) with a SP8 laser-scanning confocal microscope (Leica Microsystems), equipped with 40x water immersion objective. The Leica LAS X software was used to obtain images at a high resolution and without maximum Z-stack intensity projection (1024 pixels for both X and Y dimensions with a physical length of 83 µm, Z dimension being variable in stacks depend on cell size). All data were treated with Adobe Photoshop CS3 software at high resolution. All transfections were performed three times using independent protoplast preparations of which 10 to 20 cells were analyzed.

### SUPPLEMENTARY FIGURE LEGENDS

**Appendix Figure S1.** *In silico* analysis of RYMV-encoded P1 sequence diversity.

The amino-acid physico-chemical property is color-encoded with MVIEW. Cysteines and histidines residues are highlighted in yellow and red respectively, negatively and positively charged residues in pink and dark blue (Brown *et al*, 1998).

**Appendix Table 1. P1 constructs generated in this study.**

Primers used for restriction/ligation-based cloning (R) and for mutagenesis (M) experiments are given in regular and italic letters, respectively. Restriction sites and mutated bases in primers are in bold letters. Start and stop codons in primers are underlined. *Indicates artificial methionine residues introduced in P1 sequence for protein translation initiation purpose. [(1)] From (Gillet *et al*, 2013) ; [(2)] from (Brugidou *et al*, 1995).

**Appendix Table 2. Production and purification of recombinant P1 isomers.**

Buffers used: **Q1**: 50mM Tris pH8, 2mM DTT; **Q2**: 50mM Tris pH9, 2mM DTT; **QE**: 50mM Tris pH8, 1M NaCl, 2mM DTT; **GF1**: 25mM Tris pH8, 75mM NaCl, 2mM DTT, 300µM $ZnSO_4$; **GF2**: 25mM Tris pH8, 200mM NaCl, 2mM DTT, 300µM $ZnSO_4$; **GF3**: 25mM Tris pH8, 500mM NaCl, 2mM DTT, 300µM $ZnSO_4$; **GF4**: 25mM Tris pH8.5, 50mM $Na_2SO_4$, 2mM DTT, 150µM $ZnSO_4$, 2,5% glycerol; **GF5**: PBS pH7-7.5, 2mM DTT, 150µM $ZnSO_4$; **TS**: thermal shock on ice during 2 hours.

**Appendix Figure S2. Production of P1 recombinant sub regions in *E. coli* BL21 cells.**

**A-B** Bacteria were cultured and induced for P1 and P1 subdomains protein expression using IPTG as previously described (Gillet *et al*, 2013). Bacterial pellets from cell cultures exhibiting the same $OD_{600nm}$ were separated into soluble (**A**) and insoluble (**B**) fractions are analyzed at equal volumes on non-reducing 18% SDS-PAGE. Zinc occurrence in gels was detected using a PAR probe (lower panels) prior to Coomassie staining (upper panels). All proteins tested (P1_1-100, P1_50-157, P1_50-100, P1_102-157) were mainly soluble and colocalized with Zn. For both insoluble and soluble fractions, a bacterial extract without IPTG induction (Ni in panels) and a bacterial extract producing the full-length protein (P1 in panels) are shown as references.

**Appendix Figure S3. Circular dichroism analyses of recombinant P1 and of its sub regions.**

Spectral deconvolution of each protein (left) is given together with the percentage of the different secondary structures (right) measured at 20°C. An average of three independent measurements performed for each recombinant protein is given.

**Appendix Figure S4. Primary amino acid sequence and theoretical mass of P1 and its sub regions.**

Molecular Mass (MM) were predicted using PeptideMass software (http://web.expasy.org/peptide_mass/) (Wilkins *et al*, 1997; Gasteiger *et al*, 2005) and are given in Daltons (Da). (**A**) Full-length P1. (**B**) P1_1-100. (**C**) P1_50-100. (**D**) P1_50-157. (**E**) P1_102-157. Methionine residues artificially introduced as Met1 in proteins are in blue letters. Mutations in proteins are in red letters.

**Appendix Figure S5**. **ESI-MS quantification of zinc atoms in P1 and in its sub regions.**

Deconvoluted ESI mass spectra were obtained under denaturing conditions (upper panels) and native conditions (lower panels) for the full length P1 protein (referent protein) and its sub regions. Asterisks (*) in the peak forests and within insets indicate the detection of minor protein isomers without their N-terminal methionine (- 131 Da).

**Insets** highlight the major peak (and eventually its associated minor form depleted of the N-ter Met1) for each analyzed protein (in Da) as determined by native ESI-MS. P, PZ and PZ2 represent proteins whose molecular mass determined by ESI-MS is compatible with the absence (P), or with the presence of one (PZ) or two (PZ2) zinc atoms, respectively.

**Appendix Figure S6. ESI-MS quantification of zinc atoms in P1 and in its sub regions upon chelation by EDTA.**

All deconvoluted ESI mass spectra were obtained under native conditions. Asterisks (*) in peak forests and within insets indicate the detection of minor protein isomers without their N-terminal methionine (-131 Da).

**Insets** highlight the major peak (and eventually its associated minor form depleted of its in N-terminal Met) for each protein analyzed (in Da) as determined by native ESI-MS. P, PZ and PZ2 represent proteins whose molecular mass determined by ES-MS is compatible with the absence, or with the presence of one (PZ) or two (PZ2) zinc atoms, respectively.

**Appendix Figure S7. Determination of P1 N-terminal region zinc binding constant.**

**A** The P1_1-100 protein (40 µM) was incubated in a 40 mM HEPES-KOH buffer with or without increasing concentrations of the zinc chelator TPEN. After an overnight incubation at 25°C, the different samples were recovered and purified by PD10 column filtration. A combined PAR/PMPS assay was applied to release the remaining zinc bound to P1_100 cysteines. Values obtained arise from two independent measurements.

**B** The P1_1-100 zinc binding constant ($Ka_{P1\_1\text{-}100}$) was estimated as 2 x $10^{16}$ $M^{-1}$ according to (Jakob *et al*, 2000).

**Appendix Figure S8. Redox reactivity of P1 N- and C-terminal regions.**

**A-C** Twenty five µM of recombinant P1 (A), P1_1-100 (B) and hundred µM of P1_102-157 (C) were oxidized with increasing concentration of $H_2O_2$ (31 µM to 8 mM) at 20°C and separated on non-reducing SDS-PAGE 18%. Proteins were visualized by Coomassie blue staining (upper panel) and zinc using PAR probe (lower panel) as described in Figure 1B. Conformational changes after oxidation by $H_2O_2$ at the secondary structure level were monitored by far UV-circular dichroism (right panel) using 12 µM of each protein. $H_2O_2$ concentrations used are given on the right side of the graphic (from 31 µM in dark green to 8 mM in light green).

**D** Correlations between concentrations and $H_2O_2$/P1 ratios.

**Appendix Figure S9. Homo- and hetero-dimerization capacities of P1 N- and C-terminal regions in a yeast two-hybrid assay.**

The YRG2 strain was co-transformed with different combinations of pGAD and pGBK constructs, each one bearing a region of the P1 protein in frame with the corresponding Gal4 domain as mentioned in the table. YRG2 growth was challenged in the presence of all required amino acids including (+His) or in the absence of (-*his*) histidine. Pictures show that only the N-terminal region comprising residues from 1 to 100 can form a homodimer. The [1-100] and [102-157] regions of P1 were not able to interact in yeast cells. For all panels, cell growth was recorded at 30°C for 4 days when plates at an OD of $5.10^{-2}$ at 600 nm. Results shown here are representative of 3 independent transformation events.

**Appendix Figure S10. Zinc occurrence in P1 cysteinic and histidinic mutants.**

**A–B** Single mutations including Cys (C) to Ser (S) and His (H) to Ala (A) were introduced into the P1 nucleotide sequence by site-directed mutagenesis to produce P1 mutated recombinant proteins in the *E. Coli* BL21. Bacterial soluble (panels in A) and insoluble (panels in B) fractions expressing P1 mutants were separated by centrifugation and boiled for 5 minutes in Laemmli buffer prior to 18% SDS-PAGE analyses. Zinc occurrence in proteins was detected using a PAR staining assay (lower panels in A and B), prior to Coomassie staining (upper panels in A and B). A non-mutated P1 recombinant protein co-localizing with zinc is shown as the referent protein.

**C** Quantification of zinc atoms in P1 mutated recombinant proteins by native ESI-MS. The molecular mass determined in Daltons for each isomer is compared to that obtained using denaturing ESI-MS and to theoretical values determined based on the primary sequence of the different P1 isomers (Appendix Fig S4). Only values from major forms P (without zinc), PZ (with one zinc atom) and PZ2 (with two zinc atoms) of the different isomers comprising the N-terminal methionine are given. Deconvoluted ESI mass spectra are given in the Appendix Fig S11. Nd: not detected.

**Appendix Figure S11. ESI-MS quantification of zinc atoms in P1 cysteinic and histidinic mutants.**

Deconvoluted ESI mass spectra were obtained under denaturing (upper panels) and native (lower panels) conditions for the full length P1 related mutants. Asterisks (*) in the peak forests and within insets indicate the detection of minor protein isomers without their N-terminal methionine (-131 Da).

**Insets** highlight the major peak (and eventually its associated minor form depleted of the N-ter Met) for each protein analyzed (in Da) as determined by native ESI-MS. P, PZ and PZ2 represent proteins whose molecular mass determined by ES-MS is compatible with the absence, or with the presence of one (PZ) or two (PZ2) zinc atoms, respectively.

**Appendix Figure S12**. **Determination of zinc ion occurrence in P1_50-100 and P1_102-157 monocysteinic and monohistidinic recombinant proteins.**

**A** After production in *E. coli* and purification (Appendix Table 2), the 10 proteins carrying single (left panels, purified proteins) or double (right panels, bacterial extracts) mutations were separated on 18% acrylamide gels and incubated with the PAR probe (lower panels) prior to Coomassie staining (upper panels). Ten µg of total proteins were analyzed except for P1_50-$100^{C64S}$ and P1_50-$100^{C64S}$ (2 and 5µg, respectively) due to their very low recovery after renaturation. The mutant P1_50-$100^{C64S.C67S}$ was very poorly expressed in *E. coli* and remained undetectable in bacterial extract. Both the bacterial extract without IPTG induction and the bacterial extract producing the wild type P1 domain are shown as reference.

**B** Bacterial protein extract containing P1_102-157 protein with single Cys or His mutation were separated on 18% SDS-PAGE. Zinc occurrence in gels was detected using a PAR probe (lower panels) prior to Coomassie staining (upper panels). All proteins tested were only found in soluble fractions, among which P1_102-$157^{C119S}$ is the sole protein co-localizing with Zn. Both the bacterial extract without IPTG induction and the bacterial extract producing the wild type P1 domain are shown as references.

**C** Quantification of zinc atoms in P1_50-100 related mutants by native ESI-MS. The molecular mass determined in Daltons for each isomer is compared to that obtained using denaturing ESI-MS and to theoretical values determined based on the primary sequence of the different P1 isomers (Appendix Fig S4). Only values from major forms P (without zinc), PZ (with one zinc atom) and PZ2 (with two zinc atoms) of the different isomers comprising the N-terminal methionine are given. Deconvoluted ESI mass spectra are given in the Appendix Fig S13.

**Appendix Figure S13. ESI-MS quantification of zinc atoms in P1_50-100 monocysteinic and monohistidinic recombinant proteins.**

Deconvoluted ESI mass spectra were obtained under denaturing (upper panels) and native (lower panels) conditions for the truncated P1_50-100 protein and related mutants. Asterisks (*) in the peak forests and within insets indicate the detection of minor protein isomers without their N-terminal methionine (-131 Da).

**Insets** highlight the major peak (and eventually its associated minor form depleted of its N-ter Met) for each protein analyzed (in Da) as determined by native ESI-MS. P, PZ and PZ2 represent proteins whose molecular mass determined by ES-MS is compatible with the absence, or with the presence of one (PZ) or two (PZ2) zinc atoms, respectively.

**Appendix Figure S14. Close-ups of P1_[102-157] $^{1}$H NMR spectrum recorded at 700 MHz.**

In the presence of 1 equivalent of zinc (+Zn, upper spectrum), the $^{1}$H NMR spectrum is characteristic of a well-folded peptide, with well-scattered proton resonances ranging from 0.2 to 10 ppm. Addition of 20 mM EDTA (-Zn, lower spectrum) causes a centering of resonances in a narrower range, from 8.2 to 0.9 ppm, typical from global unfolding. The increased line widths also indicate that the unfolded peptide is prone to aggregation.

**Appendix Table 3. Phi, Psi angles estimated by Talos** (Shen & Bax, 2013) **from $^{1}$Hα, $^{13}$CO, $^{13}$Cα, $^{13}$Cβ chemical shifts.**

**Appendix Table 4. Components of the rotational diffusion tensors derived separately for P1 subdomains from experimental P1_1-157 $^{15}$N $T_1$ and $T_2$ values or computed for the NMR derived structure.**

**Appendix Figure S15. Scheme of the interactions observed at the dimeric interfaces of P1_1-100 visualized by LIGPLOT** (Wallace *et al*, 1995).

**A** Details of the interactions are depicted according to their nature: green dashed lines for H-bonds and corresponding residue names colored in yellow-orange, and dark dashed lines for hydrophobic contacts with corresponding residue names in light blue-blue. For clarity, only the main hydrophobic contact between two residues is shown. Blue spheres refer to water molecules.

**B** Summary and organization of interactions observed for P1 dimer interfaces in asymmetric and crystallographic units.

**Appendix Table 5. Surface areas of crystallographic and asymmetric P1 dimer units.**

Area surfaces were determined using the PISA software (http://www.ebi.ac.uk/msd-srv/prot_int/pistart.html) (Krissinel & Henrick, 2007).

**Appendix figure S16. Infectivity of RYMV particles and of RYMV-derived FL5 RNAs in rice.**

**A** Eleven-day old rice plantlets were manually infected either with RYMV particles (RYMV in panels) or with viral RNAs obtained by *in vitro* transcription of RYMV-derived FL5 infectious clones, either reproducing the original RYMV RNA (FL5 in panels) or carrying an ATG-to-AAG codon mutation suppressing the Met1 codon in P1 ORF (FL5ΔP1 in panels). Plants inoculated with the buffer only (Mock) were used as healthy plant controls. Plants were photographed 5 weeks post infection.

**B** Rice leaves after infection by viral RYMV particles (upper left panel) or by FL5-derived RNA (upper right panel). White arrows point-out the typical yellow mottle patterns of RYMV infected plants.

**C** Close-up view of rice leaves inoculated by viral RNAs. By comparison to the FL5 original clone (upper panels), a mutation of the P1 ATG start codon into AAG within the FL5 clone (that gives rise to FL5ΔP1) provokes a loss of viral infectivity and a healthy leaf phenotype (lower panels). Three leaf sections showing representative phenotypes observed after viral RNA inoculation are shown for each treatment.

**D** Plant height kinetics upon infection and time. Twenty plants per pot were measured every week and viral symptoms recorded. The taller leaf for each plant was used for height recording.

**E** Immunodetection of CP proteins in rice plants infected with the RYMV particles or FL5-derived RNAs. An anti-CP directed against the RYMV coat protein (Brugidou *et al*, 1995; noted @CP in panels) was used at a dilution of 1/2000 after protein transfer on membranes (15 µg of total proteins loaded per lane), and revealed by cheluminescence. The RuBisco protein levels served as loading controls.

**F** Immunodetection of P1 in rice plants inoculated with viral RYMV RNAs after separation of total rice proteins on SDS-PAGE. A purified IgG fraction from the anti-P1 polyclonal antibody (Siré *et al*, 2008; noted @P1 in panels) was used at a dilution of 1/1000 after protein transfer on membranes (15 µg of total proteins loaded per lane), and revealed by cheluminescence. The RuBisco protein levels served as transfer controls.

**Data information:** Plants were analyzed as described in Fig 5 and in Appendix Materials and Methods. All data were obtained from 30 plants per treatment, in two independent inoculation assays.

**Appendix figure S17. Effect of mutations in P1 on its localization patterns and behavior in a rice protoplast reporter system.**

pEZS.EGFP-P1 (left panels) or pEZS.P1-EGFP (right panels) constructs were transfected into rice protoplasts to analyze both P1 localization and behavior *in vivo*. The fluorescence of the EGFP protein fused to P1 or to its mutated variants (indicated in bold letters at left on the figure) was recorded 24h post-transfection by confocal microscopy. For each cell analyzed (including EGFP fluorescence, bright field and merged sections) and for both EGFP position in the fusion protein, 2 steps selected in the Z stack (here called Z step_1 and Z step_2 at left sides of the cells) are shown to highlight the cyto-nucleoplamic patterns of P1. Data shown are representation of at least 10 cells in three independent transfection assays. Bars = 10 µm.

**Appendix Figure S18. Focus on tryptophan residue positions in P1 structure.**

**A** Close-up of Trp29 (blue sticks), Trp31 (blue sticks), Trp36 (orange sticks) and Trp57 (blue sticks) side chains in P1 dimeric interface of the asymmetric unit. Residues involved in dimerization are also shown in stick representation (Met1, Thr35).

**B** Close-up of the buried position of Trp36 (orange sticks) in the hydrophobic core of P1_1-100 monomer, next to ZnF1. Interacting residues with Trp57 are also shown in stick representation (Ala41, Phe62, Thr35, Tyr71, Val90).

**Appendix Figure S19. Production and solubility of P1 Trp mutants in *E. coli***

Bacteria were cultured and induced for expression of $P1^{W29A}$, $P1^{W31A}$ and $P1^{W36A}$ recombinant mutant proteins using IPTG as previously described (Gillet *et al*, 2013). Bacterial pellets from cell cultures exhibiting the same $OD_{600nm}$ were separated into soluble (**A**) and insoluble (**B**) fractions and analyzed at equal volumes on non-reducing 18% SDS-PAGE. Zinc occurrence in gels was detected using a PAR probe (lower panels) prior to Coomassie staining (upper panels). For both insoluble and soluble fractions, the recombinant purified P1 is shown as the reference for Zinc binding.

**Appendix Figure S20. Comparison of P1 and $P1^{\Delta153\text{-}157}$ $^{1}H$ NMR spectrum recorded at 700 MHz.**

P1 and $P1^{\Delta153\text{-}157}$ $^{1}H$ NMR spectra are characteristic of well-folded peptides, with well-scattered proton resonances ranging from 0.2 to 10 ppm. No major difference is observed between P1 and its mutant indicating that deletion of the C-terminal five amino acids KYLNF does not alter P1 folding.

Poignavent *et al,* Appendix Figure S1

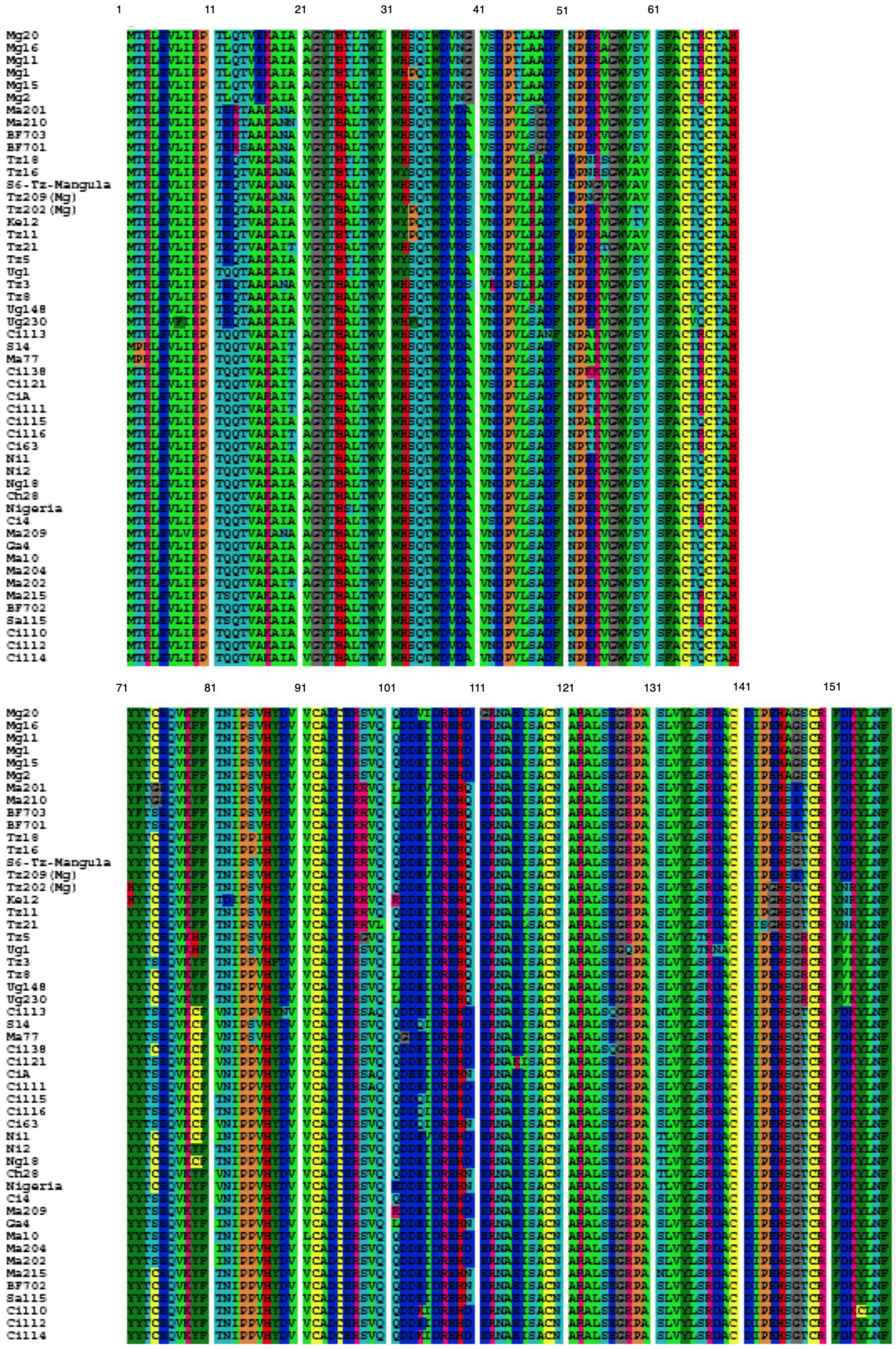

Poignavent *et al*, Appendix Table 1

Primers used for restriction/ligation-based cloning (R) and for mutagenesis (M) experiments are given in regular and italic letters, respectively. Restriction sites and mutated bases in primers are in bold letters. Start and stop codons in primers are underlined. *Indicates artificial methionine residues introduced in P1 sequence for protein translation initiation purpose. [(1)] From Gillet *et al* (2013) ; [(2)] from Brugidou *et al* (1995).

| | P1 isomer (+ mutations) | Destination vector | Cloning strategy | Template | Forward primer | Reverse primer |
|---|---|---|---|---|---|---|
| ***Full-length P1*** | | | | | | |
| P1[(1) (2)] | Met1 to Phe157 | pET.P1[(1)], FL5[(2)], pGADT7, PGBKT7 | R | | CGC**CATATG**ACACGGTTGGAAGTTCT | GCG**GGATCC**TCAGAAATTGAGGTACTTAACAAAC |
| P1 | Met1 to Phe157 | pEZS.CL, pEZS.NL | R | pET.P1[(1)] | CTGCA**GTCGAC**GATGACACGGTTGGAAGTTCTT | CGGT**GGATCC**TCAGAAATTGAGGTACTTAACAAAC |
| P1 | Met1 to Phe157 | pEZS.CL, pEZS.NL | R | pET.P1[(1)] | CTGCA**GTCGAC**GATGACACGGTTGGAAGTTCTT | CGGT**GGATCC**GAAATTGAGGTACTTAACAAAC |
| P1$^{H25A}$ | Met1 to Phe157 + *His25Ala* | pET3b | *M* | pET.P1[(1)] | *CGCCGTGGGCTATACG**GC**CGCACTCACCTGGGTTTGG* | *CCAAACCCAGGTGAGTGCG**GC**CGTATAGCCCACGGCG* |
| P1$^{W29A}$ | Met1 to Phe157 + *Trp29Ala* | pET3b, pEZS,FL5 | *M* | pET.P1[(1)], FL5[(2)] | *CGCACGCACTCACC**GC**GGTTTGGCATTCTCAG* | *CTGAGAATGCCAAACC**GC**GGTGAGTGCGTGCG* |
| P1$^{W31A}$ | Met1 to Phe157 + *Trp31Ala* | pET3b, pEZS,FL5 | *M* | pET.P1[(1)], FL5[(2)] | *GCACTCACCTGGGTT**GC**GCATTCTCAGACCTGG* | *CCAGGTCTGAGAATGC**GC**AACCCAGGTGAGTGC* |
| P1$^{H32A}$ | Met1 to Phe157 + *His32Ala* | pET3b | *M* | pET.P1[(1)] | *GCACTCACCTGGGTTTGG**GC**TTCTCAGACCTGGGACG* | *CGTCCCAGGTCTGAGAA**GC**CCAAACCCAGGTGAGTGC* |
| P1$^{W36A}$ | Met1 to Phe157 + *Trp36Ala* | pET3b, pEZS,FL5 | *M* | pET.P1[(1)], FL5[(2)] | *GGTTTGGCATTCTCAGACC**GC**GGACGTTGATTCTGTGAG* | *CTCACAGAATCAACGTCC**GC**GGTCTGAGAATGCCAAACC* |
| P1$^{W57S}$ | Met1 to Phe157 + *Trp57Ser* | pET3b, pEZS,FL5 | *M* | pET.P1[(1)], FL5[(2)] | *CCGAGAAGGTTGGTT**C**GGTGTCTGTGTCGTTC* | *GAACGACACAGACACC**G**AACCAACCTTCTCGG* |
| P1$^{C64S}$ | Met1 to Phe157 + *Cys64Ser* | pET3b, pEZS,FL5 | *M* | pET.P1[(1)] | *CTGTGTCGTTCGCCT**C**TACTCAGTGCACGGC* | *GCCGTGCACTGAGTA**G**AGGCGAACGACACAG* |
| P1$^{C67S}$ | Met1 to Phe157 + *Cys67Ser* | pET3b | *M* | pET.P1[(1)] | *TCGCCTGTACTCAGT**C**CACGGCTCACTACTA* | *TAGTAGTGAGCCGTG**G**ACTGAGTACAGGCGA* |
| P1$^{H70A}$ | Met1 to Phe157 + *His70Ala* | pET3b | *M* | pET.P1[(1)] | *CTGTACTCAGTGCACGGCT**GC**CTACTATACGAGTGAGCAG* | *CTGCTCACTCGTATAGTAG**GC**AGCCGTGCACTGAGTACAG* |
| P1$^{H87A}$ | Met1 to Phe157 + *His87Ala* | pET3b | *M* | pET.P1[(1)] | *CCAACATCCCGCCGGTT**GC**TTTCGACGTGGTGTGTG* | *CACACACCACGTCGAAA**GC**AACCGGCGGGATGTTGG* |
| P1$^{C92S}$ | Met1 to Phe157 + *Cys92Ser* | pET3b | *M* | pET.P1[(1)] | *ATTTCGACGTGGTGT**C**TGCCGATTGCGAGCG* | *CGCTCGCAATCGGCA**G**ACACCACGTCGAAAT* |
| P1$^{C95S}$ | Met1 to Phe157 + *Cys95Ser* | pET3b, pEZS,FL5 | *M* | pET.P1[(1)] | *TGGTGTGTGCCGATT**C**CGAGCGTAGTGTTCA* | *TGAACACTACGCTCG**G**AATCGGCACACACCA* |
| P1$^{H109A}$ | Met1 to Phe157 + *His109Ala* | pET3b | *M* | pET.P1[(1)] | *GACGAGATCGACCGCGAA**GC**CCAAGAGCGTAACGCAGAG* | *CTCTGCGTTACGCTCTTGG**GC**TTCGCGGTCGATCTCGTC* |
| P1$^{C119S}$ | Met1 to Phe157 + *Cys119Ser* | pET3b, pEZS,FL5 | *M* | pET.P1[(1)], FL5[(2)] | *GCAGAGATTTCTGCCT**C**CAACGCTCGGGCCTTG* | *CAAGGCCCGAGCGTTG**G**AGGCAGAAATCTCTGC* |
| P1$^{C140H}$ | Met1 to Phe157 + *Cys140His* | pEZS,FL5 | *M* | pET.P1[(1)], FL5[(2)] | *CTCTCTCGGGACGCT**CA**TGATATTCCTGAGCAC* | *GTGCTCAGGAATATCA**TG**AGCGTCCCGAGAGAG* |
| P1$^{C140S}$ | Met1 to Phe157 + *Cys140Ser* | pET3b, pEZS | *M* | pET.P1[(1)], FL5[(2)] | *CTCTCTCGGGACGCTT**C**TGATATTCCTGAGCAC* | *GTGCTCAGGAATATCA**G**AAGCGTCCCGAGAGAG* |
| P1$^{H145A}$ | Met1 to Phe157 + *His145Ala* | pET3b | *M* | pET.P1[(1)] | *GCTTGTGATATTCCTGAG**GC**CTCCGGAAGGTGCCGG* | *CCGGCACCTTCCGGAG**GC**CTCAGGAATATCACAAGC* |
| P1$^{C149H}$ | Met1 to Phe157 + *Cys149His* | pEZS,FL5 | *M* | pET.P1[(1)], FL5[(2)] | *GAGCACTCCGGAAGG**CA**CCGGTTTGTTAAGTAC* | *GTACTTAACAAACCGG**TG**CCTTCCGGAGTGCTC* |
| P1$^{C149S}$ | Met1 to Phe157 + *Cys149Ser* | pET3b, pEZS | *M* | pET.P1[(1)], FL5[(2)] | *GAGCACTCCGGAAGGT**C**CCGGTTTGTTAAGTAC* | *GTACTTAACAAACCGG**G**ACCTTCCGGAGTGCTC* |
| P1$^{C140S,C145S}$ | Met1 to Phe157 + *Cys140Ser+ Cys149Ser* | pET3b, FL5 | | pET.P1[(1)], FL5[(2)] | Same primers as for P1$^{C140S}$ and P1$^{C149S}$ | Same primers as for P1$^{C140S}$ andP1$^{C149S}$ |
| ***P1 regions*** | | | | | | |
| P1_1-100 | Met1 to Qln100 | pET3b | R | pET.P1[(1)] | CGC**CATATG**ACACGGTTGGAAGTTCT | CA**GGATCC**CTATTGAACACTACGCTCGCAAT |

| | | | | | | |
|---|---|---|---|---|---|---|
| P1_50-100 | Met50* to Qln100 | pET3b | R | pET.P1[1] | GC**CATATG**AACCCCGAGAAGGTTGGT | CA**GGATCC**CTATTGAACACTACGCTCGCAAT |
| P1_50-100$^{C64S}$ | Met50* to Qln100 + *Cys64Ser* | pET3b | *M* | pET.P1_50-100 | *CTGTGTCGTTCGCCT**C**TACTCAGTGCACGGC* | *GCCGTGCACTGAGTA**G**AGGCGAACGACACAG* |
| P1_50-100$^{C67S}$ | Met50* to Qln100 + *Cys67Ser* | pET3b | *M* | pET.P1_50-100 | *TCGCCTGTACTCAGT**C**CACGGCTCACTACTA* | *TAGTAGTGAGCCGTG**G**ACTGAGTACAGGCGA* |
| P1_50-100$^{C64S,C67S}$ | Met50* to Qln100 + *Cys64Ser,Cys67Ser* | pET3b | *M* | pET.P1_50-100$^{C64S}$ | *TCGCCTCTACTCAGT**C**CACGGCTCACTACTA* | *TAGTAGTGAGCCGTG**G**ACTGAGTAGAGGCGA* |
| P1_50-100$^{H70A}$ | Met50* to Phe100 + *His70Ala* | pET3b | *M* | pET.P1_50-100 | *CTGTACTCAGTGCACGGCT**GC**CTACTATACGAGTGAGCAG* | *CTGCTCACTCGTATAGTAG**GC**AGCCGTGCACTGAGTACAG* |
| P1_50-100$^{H87A}$ | Met50* to Phe100 + *His87Ala* | pET3b | *M* | pET.P1_50-100 | *CCAACATCCCGCCGGTT**GC**TTTCGACGTGGTGTGTG* | *CACACACCACGTCGAAA**GC**AACCGGCGGGATGTTGG* |
| P1_50-100$^{C92S}$ | Met50* to Qln100 + *Cys92Ser* | pET3b | *M* | pET.P1_50-100 | *ATTTCGACGTGGTGT**C**TGCCGATTGCGAGCG* | *CGCTCGCAATCGGCA**G**ACACCACGTCGAAAT* |
| P1_50-100$^{C95S}$ | Met50* to Qln100 + *Cys95Ser* | pET3b | *M* | pET.P1_50-100 | *TGGTGTGTGCCGATT**C**CGAGCGTAGTGTTCA* | *TGAACACTACGCTCG**G**AATCGGCACACACCA* |
| P1_50-100$^{C92S,C95S}$ | Met50* to Qln100 + *Cys92Ser,Cys95Ser* | pET3b | *M* | pET.P1_50-100$^{C92S}$ | *TGGTGTCTGCCGATT**C**CGAGCGTAGTGTTCA* | *TGAACACTACGCTCG**G**AATCGGCAGACACCA* |
| P1_50-157 | Met50* to Phe157 | pET3b | R | pET.P1[1] | GC**CATATG**AACCCCGAGAAGGTTGGT | GCG**GGATCC**TCAGAAATTGAGGTACTTAACAAAC |
| P1_102-157 | Met101* to Phe157 | pET3b | R | pET.P1[1] | GC**CATATG**GACGACGAGATCGACCGC | GCG**GGATCC**TCAGAAATTGAGGTACTTAACAAAC |
| P1_102-157$^{C119S}$ | Met101* to Phe157 + *Cys119Ser* | pET3b | *M* | pET.P1[1] | *GCAGAGATTTCTGCCT**C**CAACGCTCGGGCCTTG* | *CAAGGCCCGAGCGTTG**G**AGGCAGAAATCTCTGC* |
| P1_102-157$^{C140S}$ | Met101* to Phe157 + *Cys140Ser* | pET3b | *M* | pET.P1[1] | *CTCTCTCGGGACGCTT**C**TGATATTCCTGAGCAC* | *GTGCTCAGGAATATCA**G**AAGCGTCCCGAGAGAG* |
| P1_102-157$^{C149S}$ | Met101* to Phe157 + *Cys149Ser* | pET3b | *M* | pET.P1[1] | *GAGCACTCCGGAAGGT**C**CCGGTTTGTTAAGTAC* | *GTACTTAACAAACCGG**G**ACCTTCCGGAGTGCTC* |
| P1$^{\Delta 153-157}$ | Met1 to Asp152 + *Lys153Stop* | pEZS,FL5 | *M* | pET.P1[1], FL5[2] | *GGTGCCGGTTTGTT**T**AGTACCTCAATTTCTG* | *CAGAAATTGAGGTACT**A**AACAAACCGGCACC* |
| P1$^{\Delta 12-21}$ | [Met1-Thr11] fused to [Gly22-Phe157] | pEZS,FL5 | *M* | pET.P1[1], FL5[2] | *TATACGACCGACTGGCTATACGCACGCACTCACCTGGGTT* | *GTATAGCCAGTCGGTCGTATAAGAACTTCCAACCGTGTCA* |

Poignavent *et al*, Appendix Table 2

| **Protein and mutants** | **Production** | **Solubility** | **Renaturation** | **Ion exchange chromatography** | **Exclusion chromatography** |
|---|---|---|---|---|---|
| P1_ | 6°C | Soluble | Not required | Q1, QE | GF1, GF2, GF3, GF4, GF5 |
| P1 $^{15}$N | TS + 30°C | Insoluble | yes | Q1, QE | GF1, GF2, GF3, GF4, GF5 |
| P1 $^{15}$N $^{13}$C | TS + 30°C | Insoluble | yes | Q1, QE | GF1, GF2, GF3, GF4, GF5 |
| P1$^{H25A}$ | 6°C | Soluble | Not required | Q1, QE | GF1 |
| P1$^{W29A}$ | 6°C | Soluble | Not required | Not tested | Not tested |
| P1$^{W31A}$ | 6°C | Soluble | Not required | Not tested | Not tested |
| P1$^{H32A}$ | 6°C | Insoluble | yes | Q1, QE | GF1 |
| P1$^{W36A}$ | 6°C | Soluble | Not required | Not tested | Not tested |
| P1$^{C64S}$ | 6°C | Insoluble | yes | Q1, QE | GF1 |
| P1$^{C67S}$ | 6°C | Insoluble | yes | Q1, QE | GF1 |
| P1$^{H70A}$ | 6°C | Soluble | Not required | Q1, QE | GF1 |
| P1$^{H87A}$ | 6°C | Soluble | Not required | Q1, QE | GF1 |
| P1$^{C92S}$ | 6°C | Insoluble | yes | Q1, QE | GF1 |
| P1$^{C95S}$ | 6°C | Insoluble | yes | Q1, QE | GF1 |
| P1$^{H109A}$ | 6°C | Insoluble | yes | Q1, QE | GF1, GF3 |
| P1$^{C119S}$ | 6°C | Soluble | Not required | Q1, QE | GF1 |
| P1$^{C140S}$ | 6°C | Insoluble | yes | Q1, QE | GF1 |
| P1$^{H145A}$ | 6°C | Insoluble | yes | Q1, QE | GF1, GF3 |
| P1$^{C149S}$ | 6°C | Insoluble | yes | Q1, QE | GF1 |
| P1_1-100 | 6°C | Soluble | Not required | Q1, QE | GF1 |
| P1_1-100 $^{15}$N | TS + 25°C | Insoluble | yes | Q1, QE | GF1, GF2, GF4, GF5 |
| P1_50-100 | 6°C | Soluble | Not required | Q1, QE | GF1 |
| P1_50-157 | 6°C | Insoluble | yes | Q1, QE | GF1 |
| P1_102-157 | 6°C | Soluble | Not required | Q1, Q2, QE | GF1, GF2, GF5 |
| P1_102-157 $^{15}$N | TS + 20°C | Soluble | Not required | Q1, Q2, QE | GF1, GF2, GF5 |
| P1_50-100 $^{C64S}$ | 6°C | Insoluble | yes | Q1, QE | GF1 |
| P1_50-100 $^{C67S}$ | 6°C | Insoluble | yes | Q1, QE | GF1 |
| P1_50-100 $^{H70A}$ | 6°C | Insoluble | yes | Q1, QE | GF1 |
| P1_50-100 $^{H87A}$ | 6°C | Insoluble | yes | Q1, QE | GF1 |
| P1_50-100 $^{C92S}$ | 6°C | Insoluble | yes | Q1, QE | GF1 |
| P1_50-100 $^{C95S}$ | 6°C | Insoluble | yes | Q1, QE | GF1 |
| P1_50-100 $^{H70A-H87A}$ | 6°C | Insoluble | yes | Q1, QE | GF1 |
| P1_102-157 $^{H109A}$ | 6°C | Soluble | Not required | unsuccessful | - |
| P1_102-157 $^{C119S}$ | 6°C | Soluble | Not required | Q1, QE | GF1 |
| P1_102-157 $^{C140S}$ | 6°C | Soluble | Not required | unsuccessful | - |
| P1_102-157 $^{H145A}$ | 6°C | Soluble | Not required | unsuccessful | - |
| P1_102-157 $^{C145S}$ | 6°C | Soluble | Not required | unsuccessful | - |

Poignavent *et al*, Appendix Figure S2

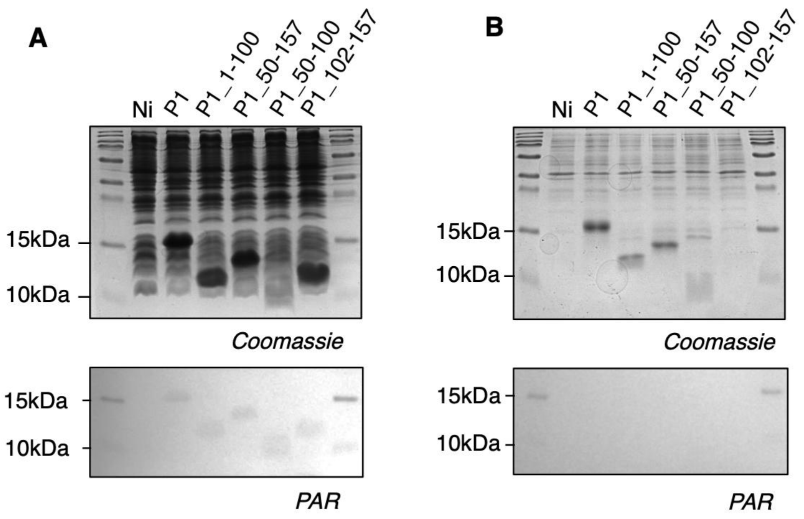

Poignavent *et al*, Appendix Figure S3

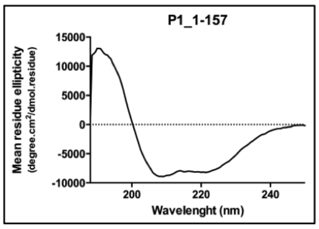

| P1_1-157 | 180-260 nm | 185-260 nm | 190-260 nm | 195-260 nm | 200-260 nm | 205-260 nm | 210-260 nm |
|---|---|---|---|---|---|---|---|
| α Helix | 43,20% | 41,40% | 40,40% | 39,30% | 39,60% | 40,20% | 39,20% |
| β Antiparallel | 3,50% | 3,90% | 4,40% | 6,50% | 6,80% | 6,50% | 6,80% |
| β Parallel | 6,70% | 7,00% | 7,30% | 7,30% | 7,30% | 7,30% | 7,40% |
| β Turn | 14,90% | 15,20% | 15,50% | 15,70% | 15,70% | 15,60% | 15,80% |
| Non structured | 27,60% | 28,00% | 28,50% | 28,80% | 28,80% | 28,80% | 29,40% |
| Total | 95,90% | 95,50% | 96,10% | 97,60% | 98,20% | 98,40% | 98,60% |

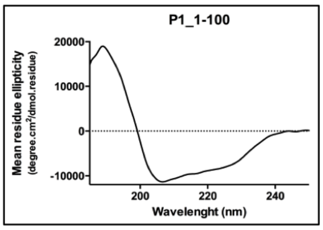

| P1_1-100 | 180-260 nm | 185-260 nm | 190-260 nm | 195-260 nm | 200-260 nm | 205-260 nm | 210-260 nm |
|---|---|---|---|---|---|---|---|
| α Helix | 32,60% | 31,40% | 30,80% | 29,50% | 29,60% | 32,50% | 30,20% |
| β Antiparallel | 11,20% | 12,50% | 14,10% | 12,90% | 10,10% | 8,10% | 9,10% |
| β Parallel | 7,80% | 8,10% | 8,10% | 8,90% | 9,30% | 9,20% | 9,50% |
| β Turn | 17,70% | 17,90% | 18,00% | 17,90% | 17,90% | 16,90% | 17,50% |
| Non structured | 27,80% | 28,40% | 28,30% | 30,20% | 32,50% | 33,70% | 35,30% |
| Total | 97,30% | 98,30% | 99,30% | 99,50% | 99,30% | 100,50% | 101,60% |

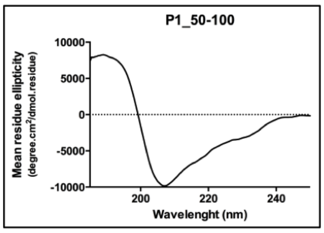

| P1_50-100 | 180-260 nm | 185-260 nm | 190-260 nm | 195-260 nm | 200-260 nm | 205-260 nm | 210-260 nm |
|---|---|---|---|---|---|---|---|
| α Helix | 25,00% | 25,00% | 25,10% | 24,50% | 23,60% | 24,40% | 21,90% |
| β Antiparallel | 23,00% | 23,30% | 24,00% | 17,40% | 12,80% | 10,40% | 12,40% |
| β Parallel | 9,70% | 9,60% | 9,40% | 10,60% | 11,50% | 12,30% | 12,20% |
| β Turn | 19,80% | 19,70% | 19,40% | 19,00% | 19,40% | 18,50% | 19,40% |
| Non structured | 31,50% | 32,10% | 31,60% | 34,30% | 37,30% | 40,40% | 42,20% |
| Total | 109,10% | 109,80% | 109,60% | 105,80% | 104,50% | 105,90% | 108,20% |

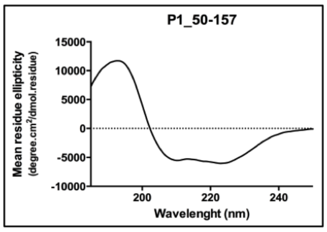

| P1_50-157 | 180-260 nm | 185-260 nm | 190-260 nm | 195-260 nm | 200-260 nm | 205-260 nm | 210-260 nm |
|---|---|---|---|---|---|---|---|
| α Helix | 25,50% | 25,40% | 25,40% | 23,80% | 21,40% | 20,60% | 20,10% |
| β Antiparallel | 15,90% | 16,40% | 16,50% | 15,30% | 13,10% | 11,70% | 13,30% |
| β Parallel | 11,10% | 10,80% | 10,70% | 11,90% | 13,20% | 14,20% | 12,90% |
| β Turn | 18,30% | 18,40% | 18,30% | 18,60% | 19,30% | 19,10% | 19,80% |
| Non structured | 39,30% | 39,70% | 39,20% | 40,90% | 43,40% | 44,60% | 44,00% |
| Total | 110,10% | 110,60% | 110,10% | 110,50% | 110,40% | 110,30% | 110,20% |

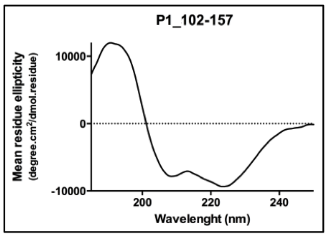

| P1_102-157 | 180-260 nm | 185-260 nm | 190-260 nm | 195-260 nm | 200-260 nm | 205-260 nm | 210-260 nm |
|---|---|---|---|---|---|---|---|
| α Helix | 28,50% | 28,20% | 28,10% | 27,30% | 26,40% | 26,90% | 25,90% |
| β Antiparallel | 12,80% | 13,20% | 13,50% | 12,70% | 10,70% | 9,40% | 10,50% |
| β Parallel | 9,90% | 9,70% | 9,70% | 10,30% | 10,90% | 11,20% | 10,70% |
| β Turn | 17,80% | 17,90% | 17,80% | 17,90% | 18,10% | 17,70% | 18,30% |
| Non structured | 35,70% | 36,00% | 35,80% | 36,70% | 38,20% | 38,60% | 38,70% |
| Total | 104,70% | 105,00% | 105,00% | 104,90% | 104,30% | 103,80% | 104,10% |

Poignavent *et al*, Appendix Figure S4

(**A**) **Full-length P1 protein** (NCBI : gi|45535303|emb|CAE81331.1|), **157 aminoacids**

MM : 17929.01 Da
MTRLEVLIRPTEQTAAKANAVGYTHALTWVWHSQTWDVDSVRDPSLRADFNPEKVGWVSVSFACTQ
CTAHYYTSEQVKYFTNIPPVHFDVVCADCERSVQLDDEIDREHQERNAEISACNARALSEGRPASL
VYLSRDACDIPEHSGRCRFVKYLNF

Mutants in Full-length P1 :

P1$^{C64S}$ : 17912.9 Da

MTRLEVLIRPTEQTAAKANAVGYTHALTWVWHSQTWDVDSVRDPSLRADFNPEKVGWVSVSFA**S**TQ
CTAHYYTSEQVKYFTNIPPVHFDVVCADCERSVQLDDEIDREHQERNAEISACNARALSEGRPASL
VYLSRDACDIPEHSGRCRFVKYLNF

P1$^{C67S}$ : 17912.9 Da

MTRLEVLIRPTEQTAAKANAVGYTHALTWVWHSQTWDVDSVRDPSLRADFNPEKVGWVSVSFACTQ
**S**TAHYYTSEQVKYFTNIPPVHFDVVCADCERSVQLDDEIDREHQERNAEISACNARALSEGRPASL
VYLSRDACDIPEHSGRCRFVKYLNF

P1$^{H70A}$ : 17862.9 Da

MTRLEVLIRPTEQTAAKANAVGYTHALTWVWHSQTWDVDSVRDPSLRADFNPEKVGWVSVSFACTQ
CTA**A**YYTSEQVKYFTNIPPVHFDVVCADCERSVQLDDEIDREHQERNAEISACNARALSEGRPASL
VYLSRDACDIPEHSGRCRFVKYLNF

P1$^{H87A}$ : 17862.9 Da

MTRLEVLIRPTEQTAAKANAVGYTHALTWVWHSQTWDVDSVRDPSLRADFNPEKVGWVSVSFACTQ
CTAHYYTSEQVKYFTNIPPV**A**FDVVCADCERSVQLDDEIDREHQERNAEISACNARALSEGRPASL
VYLSRDACDIPEHSGRCRFVKYLNF

P1$^{C92S}$ : 17912.9 Da

MTRLEVLIRPTEQTAAKANAVGYTHALTWVWHSQTWDVDSVRDPSLRADFNPEKVGWVSVSFACTQ
CTAHYYTSEQVKYFTNIPPVHFDVV**S**ADCERSVQLDDEIDREHQERNAEISACNARALSEGRPASL
VYLSRDACDIPEHSGRCRFVKYLNF

P1$^{C95S}$ : 17912.9 Da

MTRLEVLIRPTEQTAAKANAVGYTHALTWVWHSQTWDVDSVRDPSLRADFNPEKVGWVSVSFACTQ
CTAHYYTSEQVKYFTNIPPVHFDVVCAD**S**ERSVQLDDEIDREHQERNAEISACNARALSEGRPASL
VYLSRDACDIPEHSGRCRFVKYLNF

P1$^{H109A}$ : 17862.9 Da

MTRLEVLIRPTEQTAAKANAVGYTHALTWVWHSQTWDVDSVRDPSLRADFNPEKVGWVSVSFACTQ
CTAHYYTSEQVKYFTNIPPVHFDVVCADCERSVQLDDEIDRE**A**QERNAEISACNARALSEGRPASL
VYLSRDACDIPEHSGRCRFVKYLNF

Poignavent *et al*, Appendix Figure S4

P1$^{C119S}$ : 17912.9 Da

MTRLEVLIRPTEQTAAKANAVGYTHALTWVWHSQTWDVDSVRDPSLRADFNPEKVGWVSVSFACTQ
CTAHYYTSEQVKYFTNIPPVHFDVVCADCERSVQLDDEIDREHQERNAEISA**S**NARALSEGRPASL
VYLSRDACDIPEHSGRCRFVKYLNF

P1$^{C140S}$ : 17912.9 Da

MTRLEVLIRPTEQTAAKANAVGYTHALTWVWHSQTWDVDSVRDPSLRADFNPEKVGWVSVSFACTQ
CTAHYYTSEQVKYFTNIPPVHFDVVCADCERSVQLDDEIDREHQERNAEISACNARALSEGRPASL
VYLSRDA**S**DIPEHSGRCRFVKYLNF

P1$^{H145A}$ : 17862.9 Da

MTRLEVLIRPTEQTAAKANAVGYTHALTWVWHSQTWDVDSVRDPSLRADFNPEKVGWVSVSFACTQ
CTAHYYTSEQVKYFTNIPPVHFDVVCADCERSVQLDDEIDREHQERNAEISACNARALSEGRPASL
VYLSRDACDIPE**A**SGRCRFVKYLNF

P1$^{C149S}$ : 17912.9 Da

MTRLEVLIRPTEQTAAKANAVGYTHALTWVWHSQTWDVDSVRDPSLRADFNPEKVGWVSVSFACTQ
CTAHYYTSEQVKYFTNIPPVHFDVVCADCERSVQLDDEIDREHQERNAEISACNARALSEGRPASL
VYLSRDACDIPEHSGR**S**RFVKYLNF

**(B) P1_1-100 protein, 100 aminoacids**

MM : 11408.80 Da
MTRLEVLIRPTEQTAAKANAVGYTHALTWVWHSQTWDVDSVRDPSLRADFNPEKVGWVSVSFACTQ
CTAHYYTSEQVKYFTNIPPVHFDVVCADCERSVQ

**(C) P1_50-100 protein (alias ZnF1), 51 aminoacids**

MM : 5816.57 Da
MNPEKVGWVSVSFACTQCTAHYYTSEQVKYFTNIPPVHFDVVCADCERSVQ

Mutants in ZnF1 :
P1_50-100$^{C64S}$ : 5800.51 Da
MNPEKVGWVSVSFA**S**TQCTAHYYTSEQVKYFTNIPPVHFDVVCADCERSVQ
P1_50-100$^{C67S}$ : 5800.51 Da
MNPEKVGWVSVSFACTQ**S**TAHYYTSEQVKYFTNIPPVHFDVVCADCERSVQ
P1_50-100$^{C64S/C67S}$ : 5784.45 Da
MNPEKVGWVSVSFA**S**TQ**S**TAHYYTSEQVKYFTNIPPVHFDVVCADCERSVQ
P1_50-100$^{H70A}$ : 5750.50 Da
MNPEKVGWVSVSFACTQCTA**A**YYTSEQVKYFTNIPPVHFDVVCADCERSVQ
P1_50-100$^{H87A}$ : 5750.50 Da
MNPEKVGWVSVSFACTQCTAHYYTSEQVKYFTNIPPV**A**FDVVCADCERSVQ
P1_50-100$^{H70A/H87A}$ : 5684.44 Da
MNPEKVGWVSVSFACTQCTA**A**YYTSEQVKYFTNIPPV**A**FDVVCADCERSVQ

Poignavent *et al*, Appendix Figure S4

P1_50-100$^{C92S}$ : 5800.51 Da
MNPEKVGWVSVSFACTQCTAHYYTSEQVKYFTNIPPVHFDVV**S**ADCERSVQ
P1_50-100$^{C95S}$ : 5800.51 Da
MNPEKVGWVSVSFACTQCTAHYYTSEQVKYFTNIPPVHFDVVCAD**S**ERSVQ
P1_50-100$^{C92S/C95S}$ : 5784.45 Da
MNPEKVGWVSVSFACTQCTAHYYTSEQVKYFTNIPPVHFDVV**S**AD**S**ERSVQ

(**D**) **P1_50-157 protein, 108 aminoacids**

MM : 12336.78 Da
MNPEKVGWVSVSFACTQCTAHYYTSEQVKYFTNIPPVHFDVVCADCERSVQLDDEIDREHQERNAE
ISACNARALSEGRPASLVYLSRDACDIPEHSGRCRFVKYLNF

(**E**) **P1_102-157 protein (alias ZnF2), 57 aminoacids**

MM : 6556.26 Da
MDDEIDREHQERNAEISACNARALSEGRPASLVYLSRDACDIPEHSGRCRFVKYLNF

Mutants in P1_102-157:
P1_102-157$^{H109A}$ : 6490.20 Da
MDDEIDRE**A**QERNAEISACNARALSEGRPASLVYLSRDACDIPEHSGRCRFVKYLNF

P1_102-157$^{C119S}$ : 6540.20 Da
MDDEIDREHQERNAEISA**S**NARALSEGRPASLVYLSRDACDIPEHSGRCRFVKYLNF

P1_102-157$^{H145A}$ : 6490.20 Da
MDDEIDREHQERNAEISACNARALSEGRPASLVYLSRDACDIPE**A**SGRCRFVKYLNF

P1_102-157$^{C140S}$ : 6540.20 Da
MDDEIDREHQERNAEISACNARALSEGRPASLVYLSRDA**S**DIPEHSGRCRFVKYLNF

P1_102-157$^{C149S}$ : 6540.20 Da
MDDEIDREHQERNAEISACNARALSEGRPASLVYLSRDACDIPEHSGR**S**RFVKYLNF

P1_102-157$^{C140S/C149S}$ : 6524.14 Da
MDDEIDREHQERNAEISACNARALSEGRPASLVYLSRDA**S**DIPEHSGR**S**RFVKYLNF

Poignavent *et al*, Appendix Figure S5

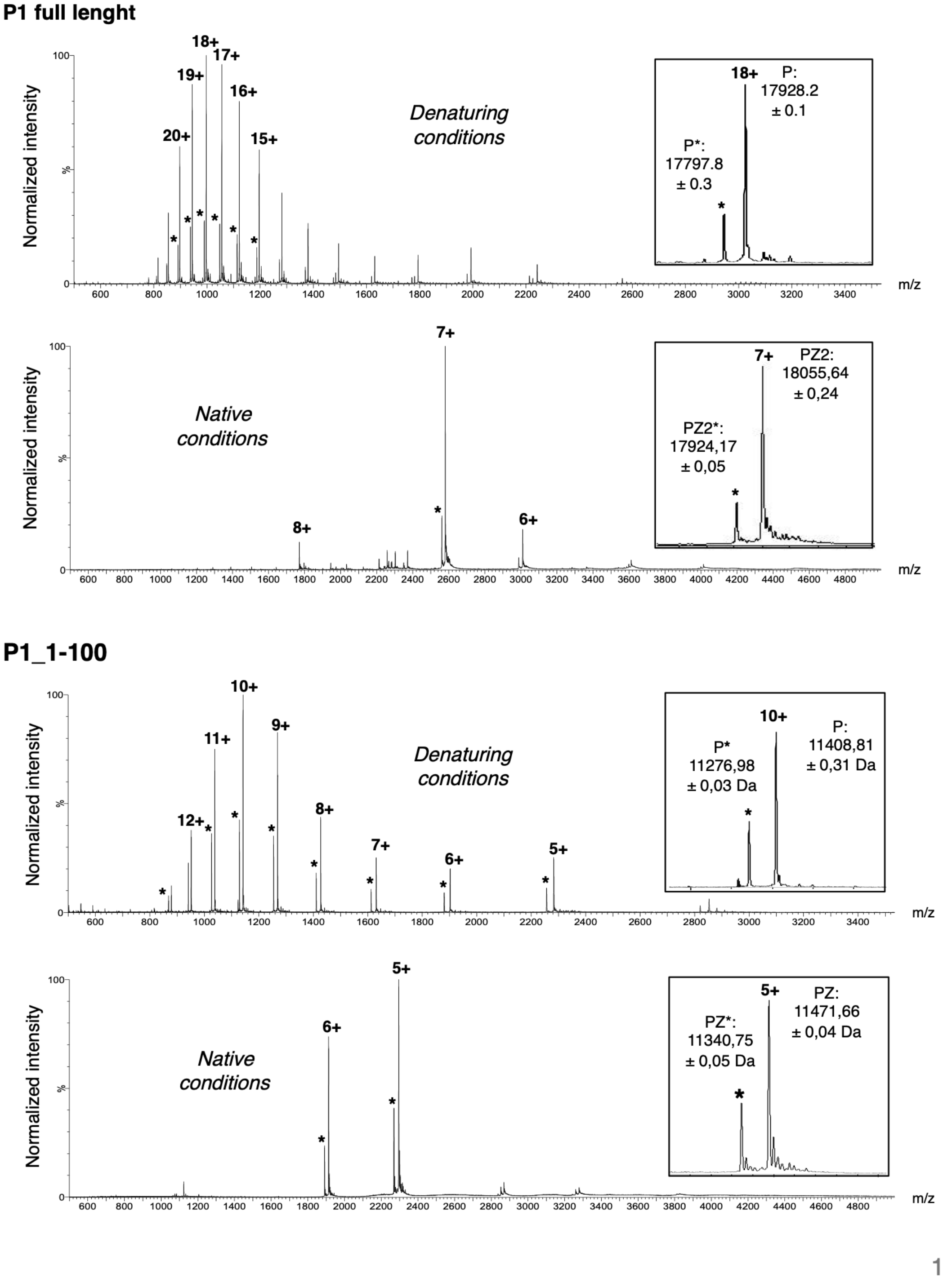

Poignavent *et al*, Appendix Figure S5

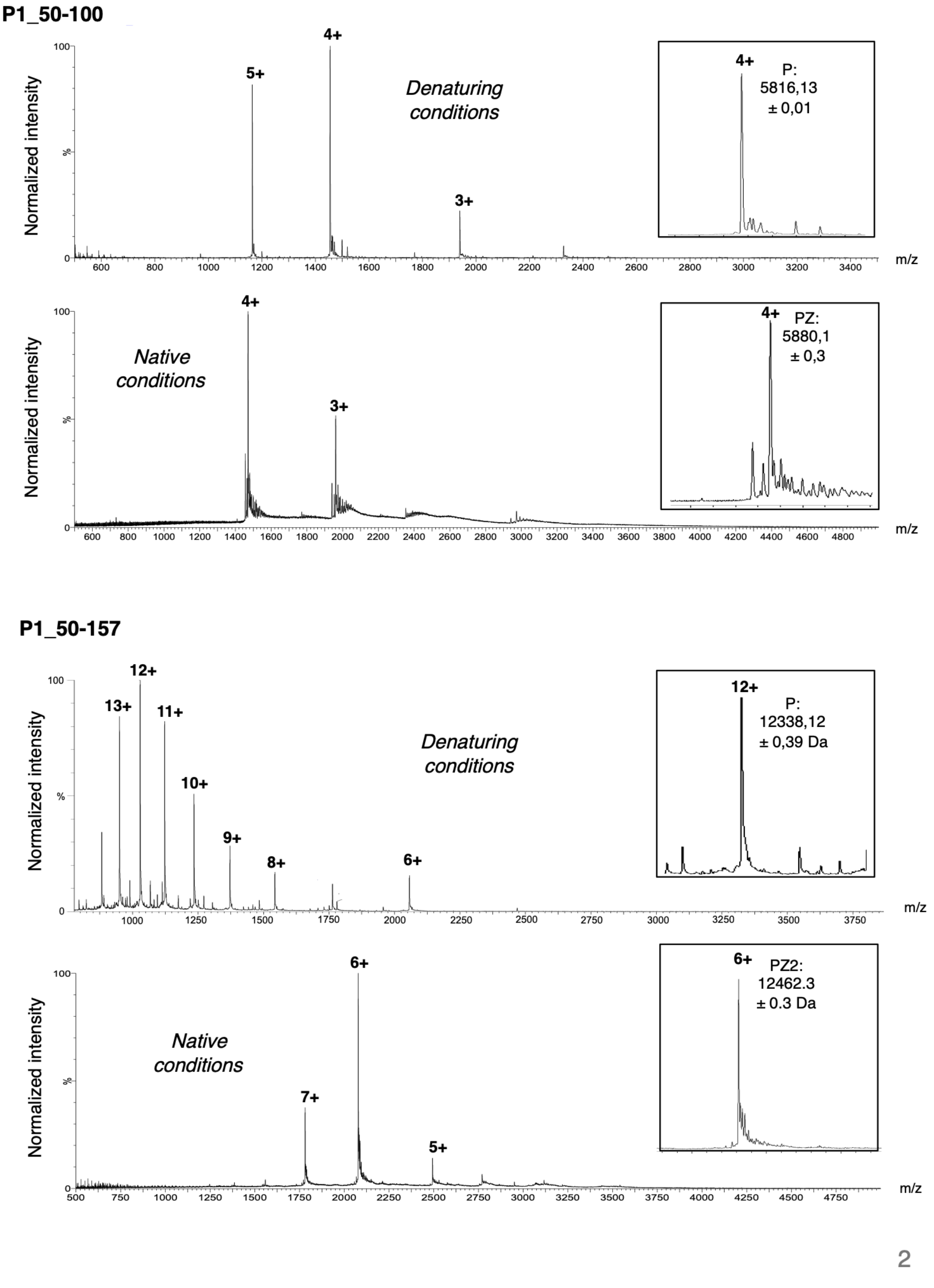

Poignavent *et al*, Appendix Figure S5

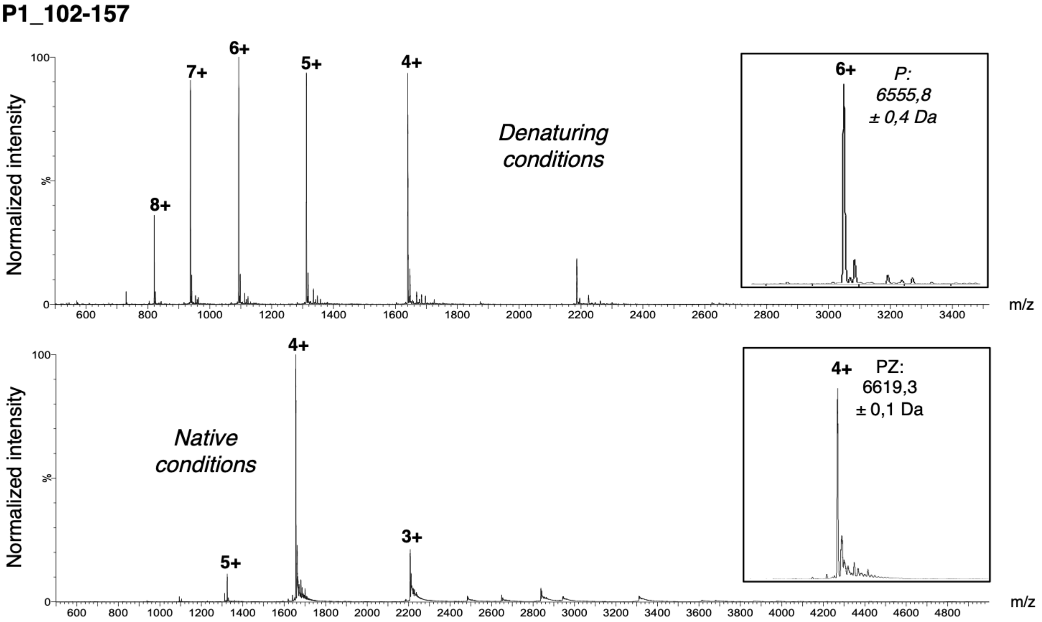

3

Poignavent *et al*, Appendix Figure S6

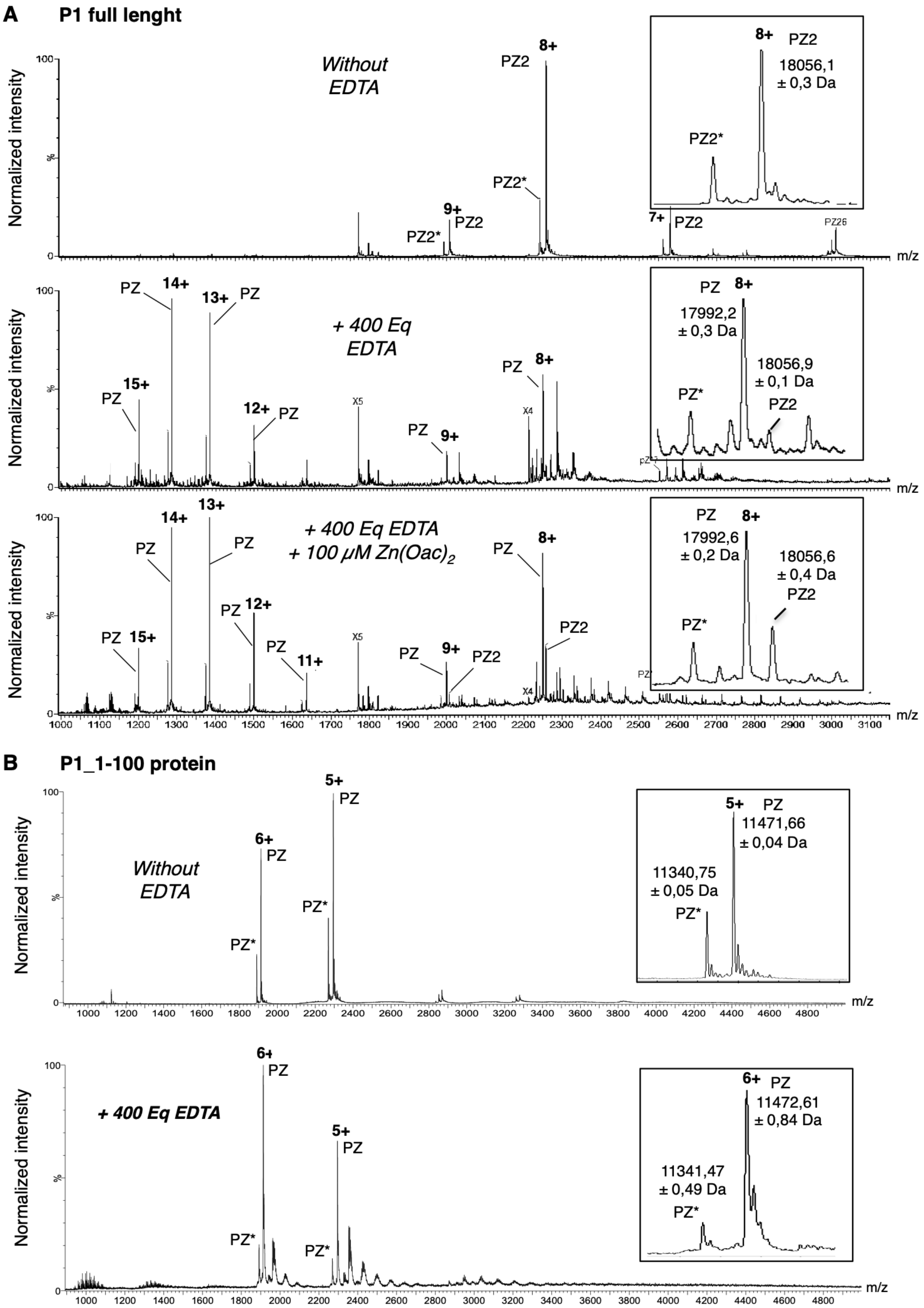

Poignavent *et al*, Appendix Figure S6

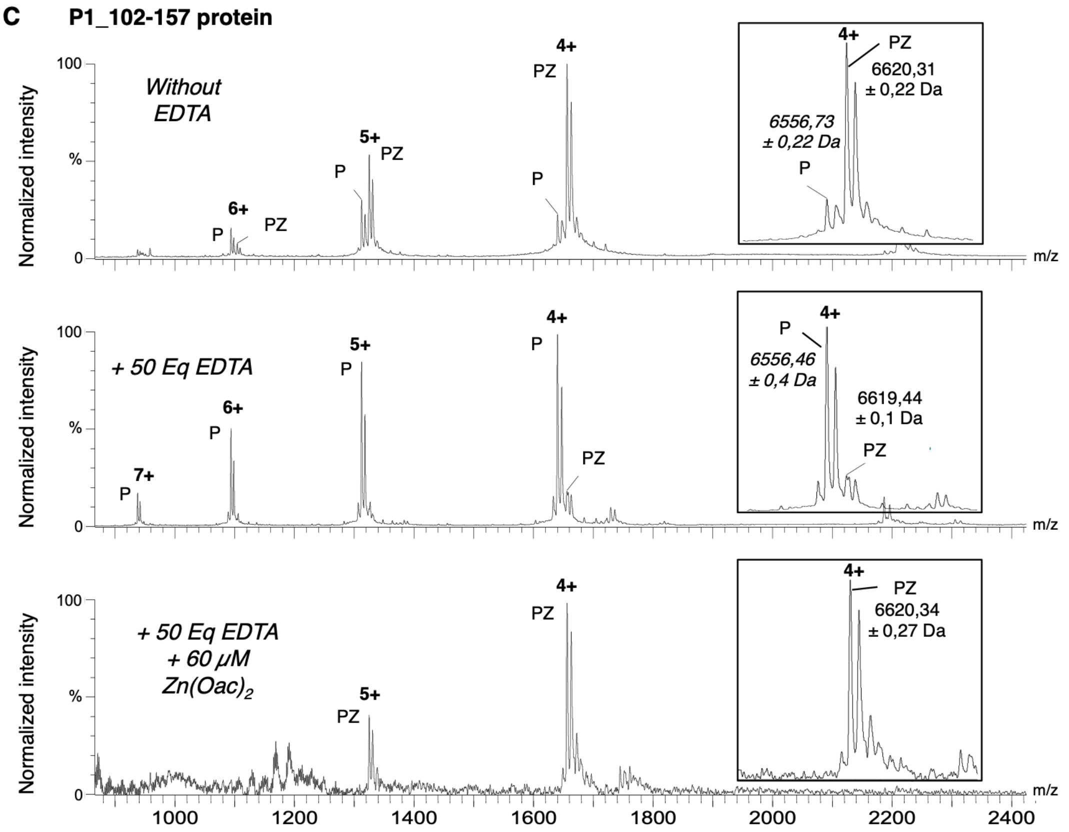

Poignavent *et al*, Appendix Figure S7

A

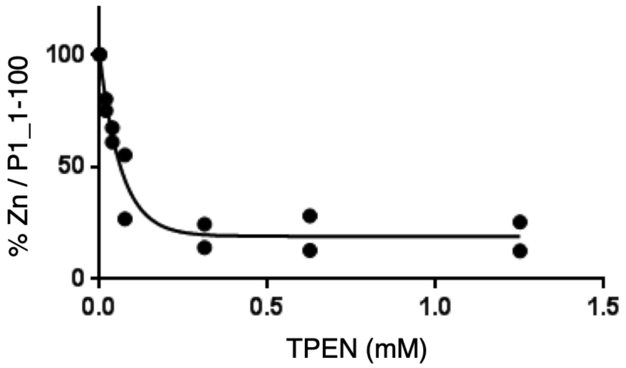

B

$$Ka_{P1\_1\text{-}100} = \frac{(Ka_{\ TPEN} \times [P1\_1\text{-}100\ Zn] \times [free\ TPEN])}{[TPEN\ Zn] \times [free\ P1\_1\text{-}100]} = 2{,}15 \times 10^{16}\ M^{-1}$$

Poignavent *et al*, Appendix Figure S8

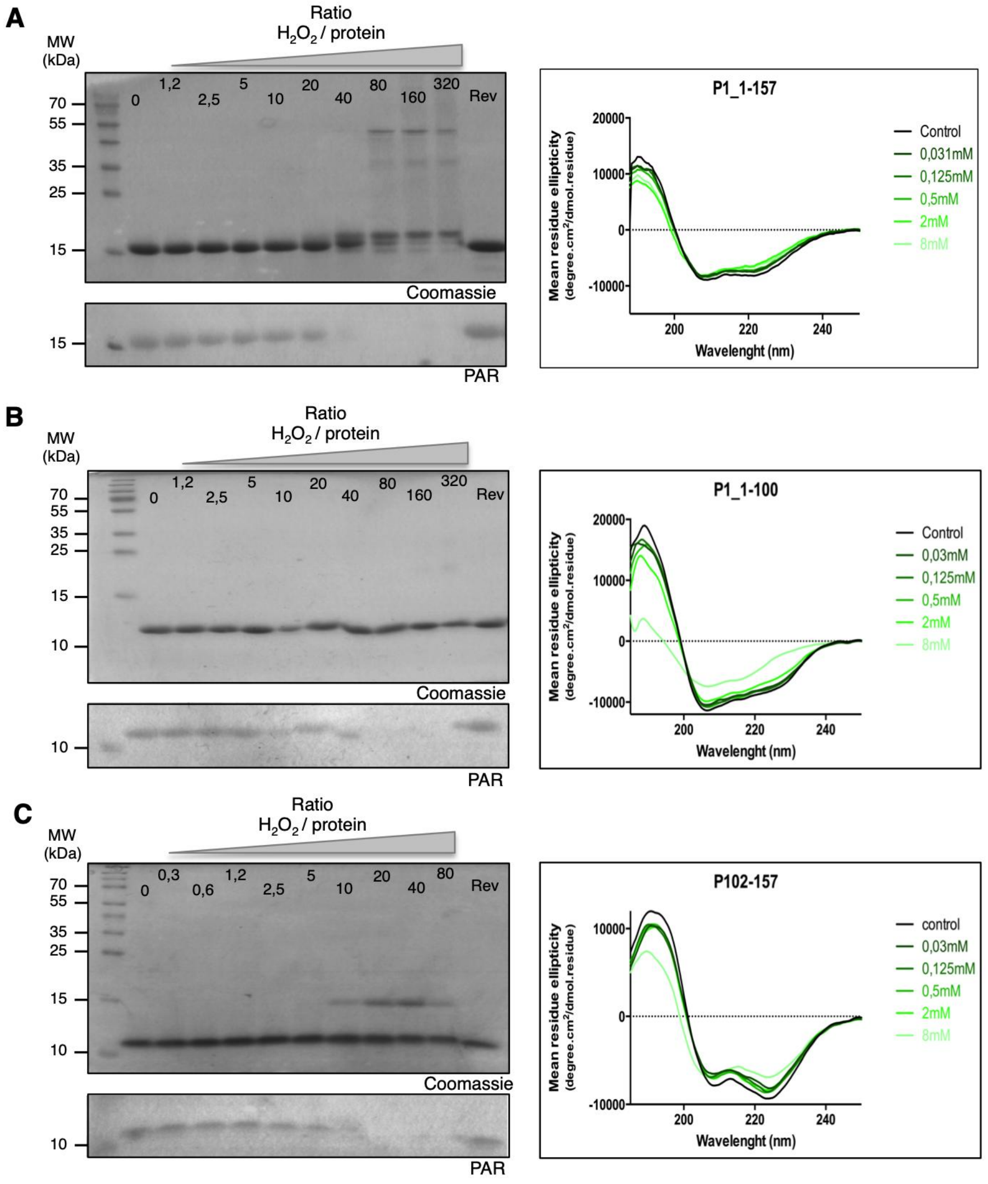

**D**

| **$[H_2O_2]$ in mM** | **0.031** | **0.062** | **0.125** | **0.250** | **0.5** | **1** | **2** | **4** | **8** |
|---|---|---|---|---|---|---|---|---|---|
| **$H_2O_2$/Protein (25 *µ*M)** | 1.2 | 2.4 | 5 | 10 | 20 | 40 | 80 | 160 | 320 |
| **$H_2O_2$/Protein (100 *µ*M)** | 0.3 | 0.6 | 1.2 | 2.5 | 5 | 10 | 20 | 40 | 80 |

Poignavent *et al*, Appendix Figure S9

| AD-Gal4 construct | BD-Gal4 construct |
|---|---|
| P1_1-100 | empty |
| P1_102-157 | empty |
| empty | P1_1-100 |
| empty | P1_102-157 |
| P1_1-100 | P1_1-100 |
| P1_102-157 | P1_102-157 |
| P1_102-157 | P1_1-100 |

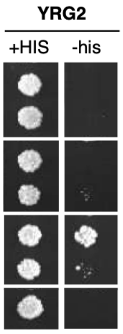

Poignavent *et al*, Appendix Figure S10

**A**

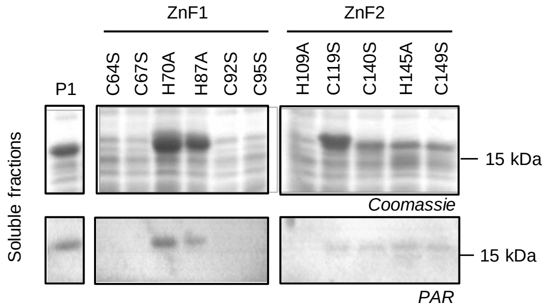

**B**

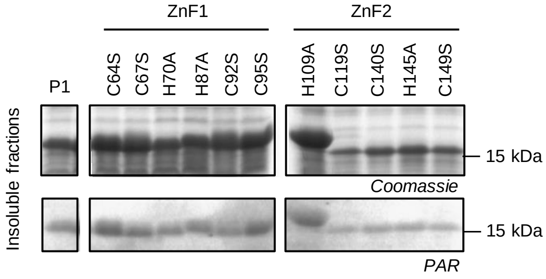

**C**

| Proteins | Theoretical molecular mass | Molecular mass by ESI-MS denaturant | Molecular mass by native ESI-MS (major form PZ) | Difference between native ESI-MS and theoretical mass | Number of zinc atom (63 to 65 Da) |
|---|---|---|---|---|---|
| **P1** | 17929.01 | 17928,2 ± 0,01 | 18055,64 ± 0,24 | 126,63 | 2 |
| **P1**$^{C64S}$ | 17912.9 | 17911,96 ± 0,33 | 18039,9 ± 0,55 | 127 | 2 |
| **P1**$^{C67S}$ | 17912.9 | 17911,89 ±0,15 | 18041,06 ± 0,9 | 128,16 | 2 |
| **P1**$^{H70A}$ | 17862.9 | 17861,98 ± 0,11 | 17991,36 ± 0,67 | 128,46 | 2 |
| **P1**$^{H87A}$ | 17862.9 | 17861,96 ± 0,36 | 17991,38 ± 2,1 | 128,48 | 2 |
| **P1**$^{C92S}$ | 17912.9 | 17911,31 ±0,23 | 18040,88 ± 0,14 | 127,98 | 2 |
| **P1**$^{C95S}$ | 17912.9 | 17911,68 ±0,26 | 18041,42 ± 4,15 | 128,52 | 2 |
| | | | 17979,37 ± 0,5 | 66,47 | 1 |
| **P1**$^{H109A}$ | 17862.9 | Nd | Nd | Nd | Nd |
| **P1**$^{C119S}$ | 17912.9 | 17913,0 ±0,20 | 18039,3 ± 0,4 | 126,4 | 2 |
| **P1**$^{C140S}$ | 17912.9 | 17911,44 ±0,26 | 18042,41 ± 3,04 | 129,51 | 2 |
| **P1**$^{H145A}$ | 17862.9 | 17861,05 ± 0,13 | 17993,59 ± 1,46 | 130,69 | 2 |
| **P1**$^{C149S}$ | 17912.9 | Nd | 18043,61 ± 0,39 | 130,71 | 2 |

Poignavent *et al*, Appendix Figure S11

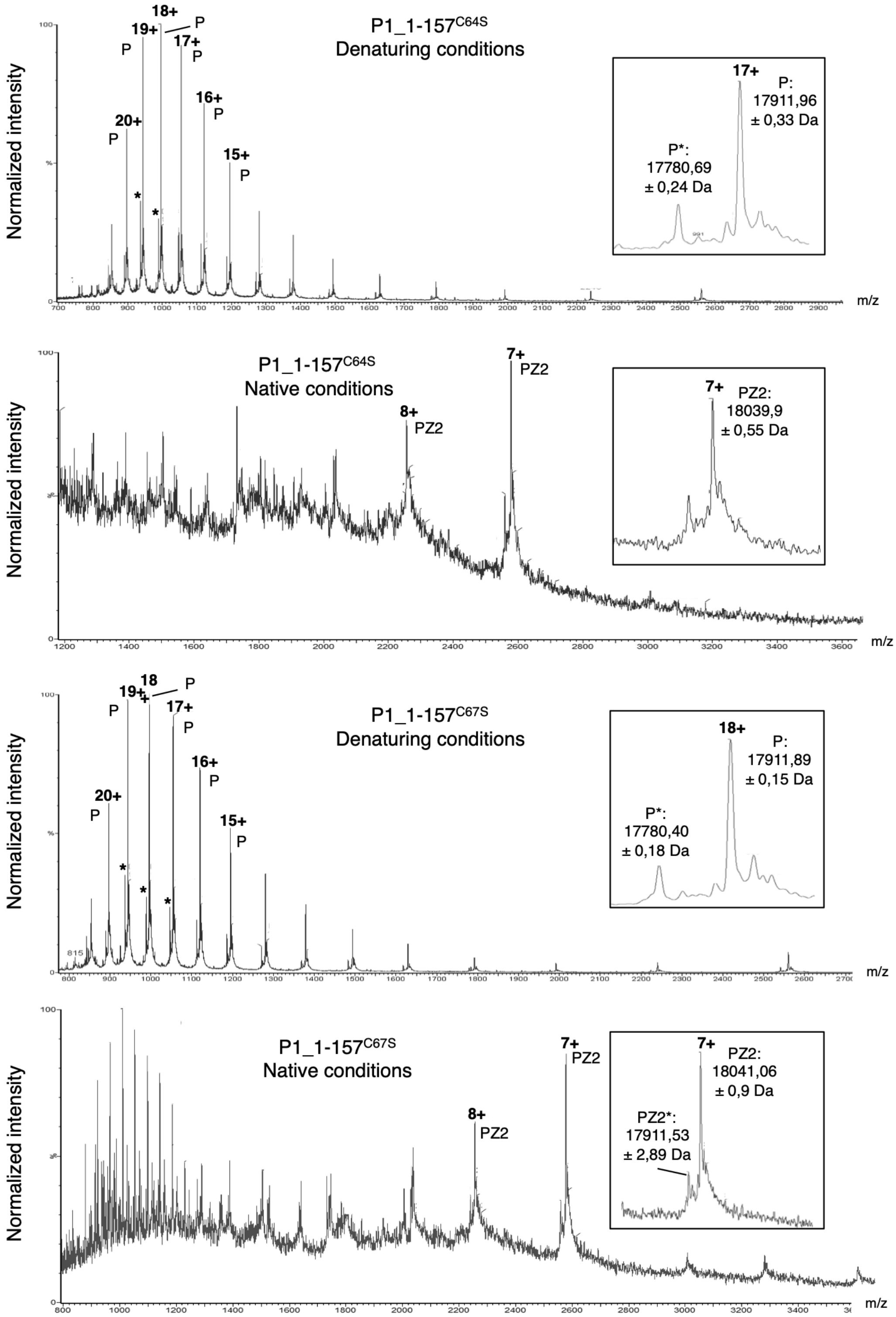

Poignavent *et al*, Appendix Figure S11

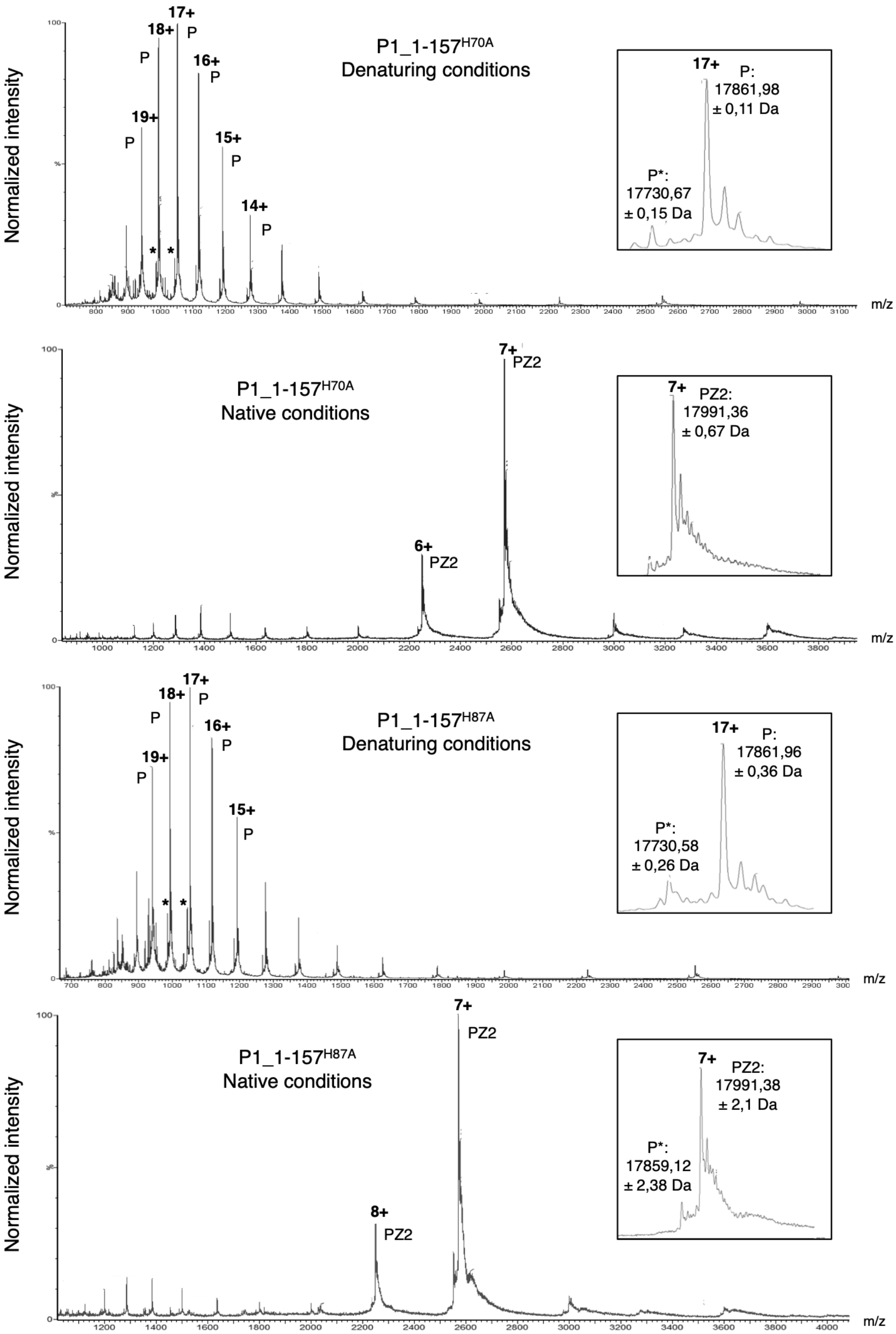

Poignavent *et al*, Appendix Figure S11

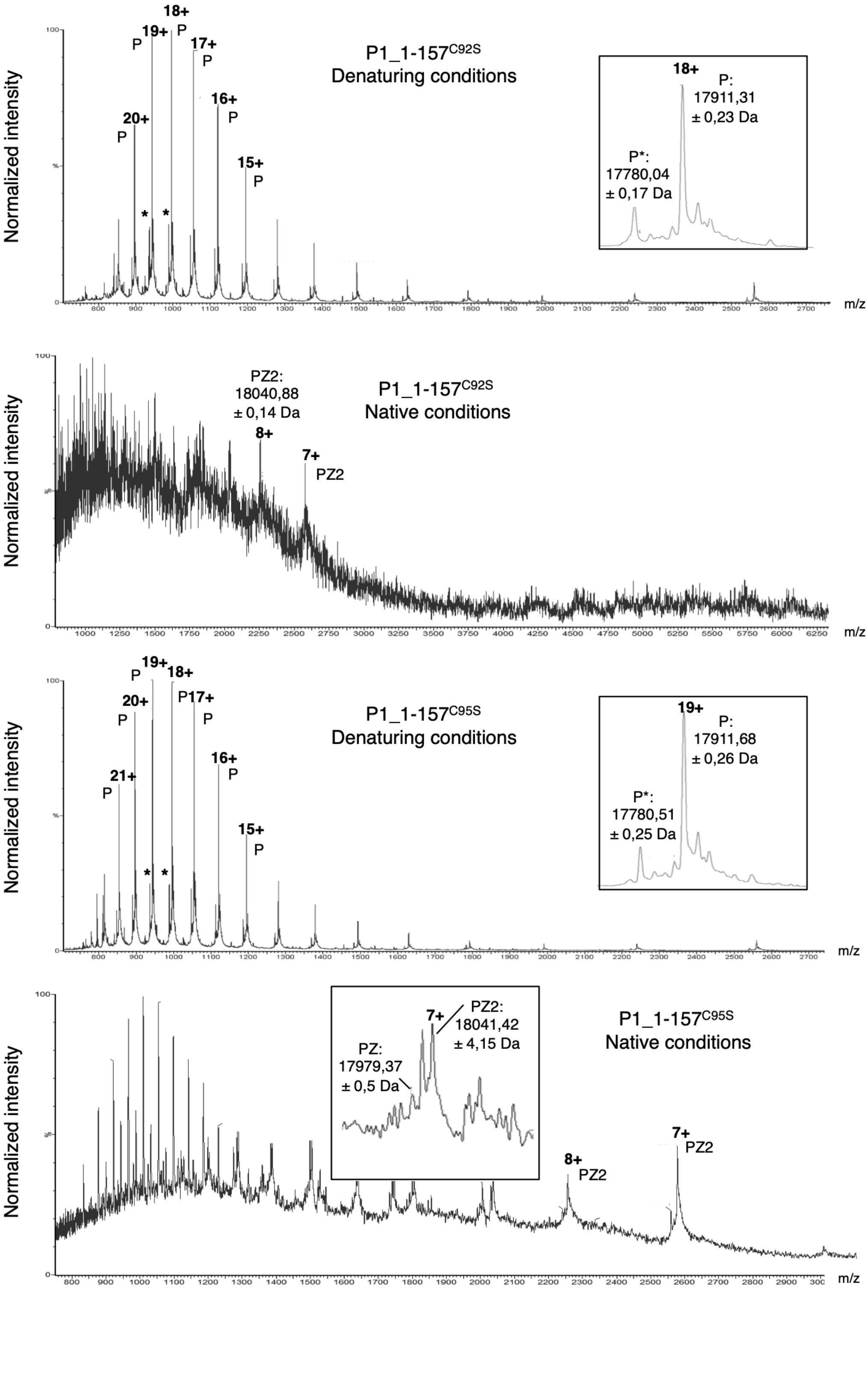

Poignavent *et al*, Appendix Figure S11

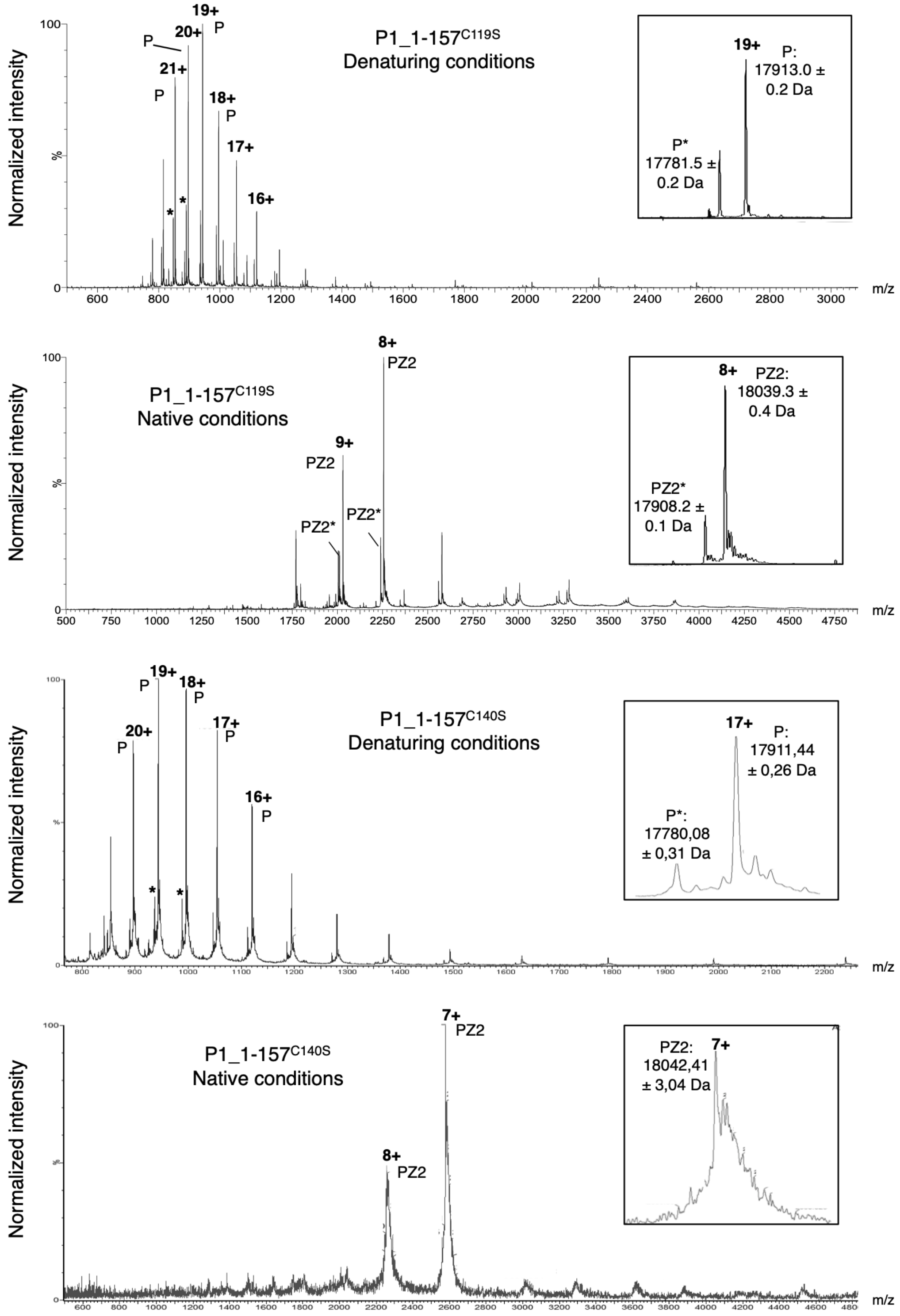

Poignavent *et al*, Appendix Figure S11

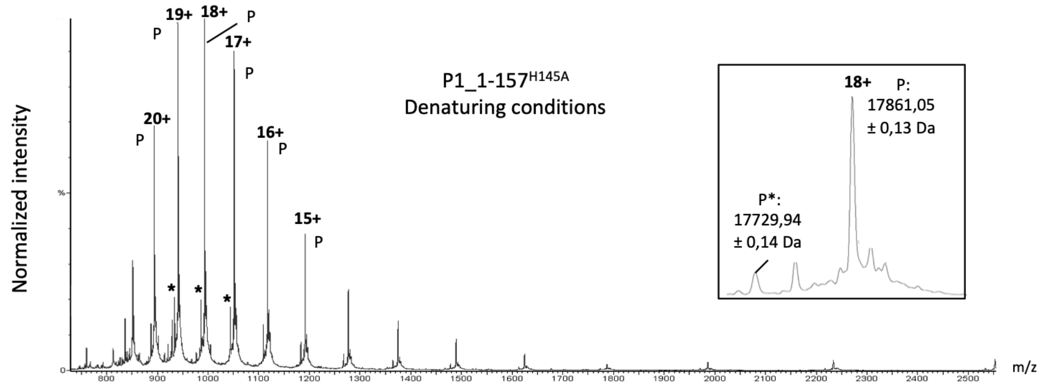

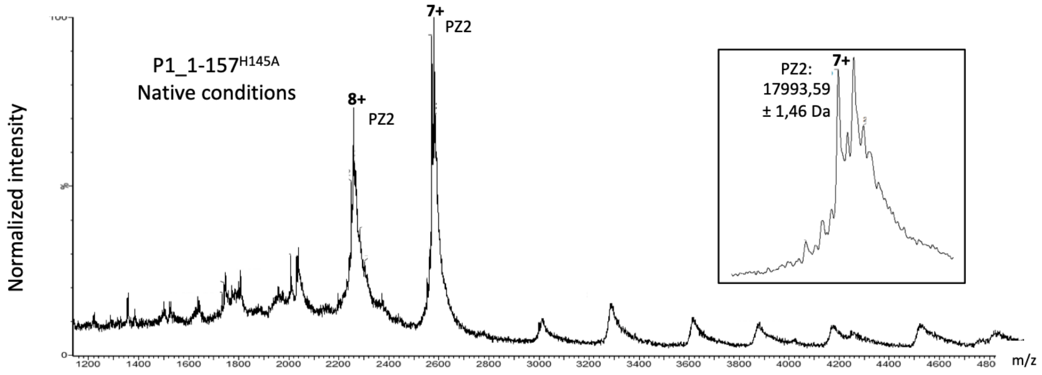

**P1_1-157$^{C149S}$**

Denaturing conditions

*No spectrum available*

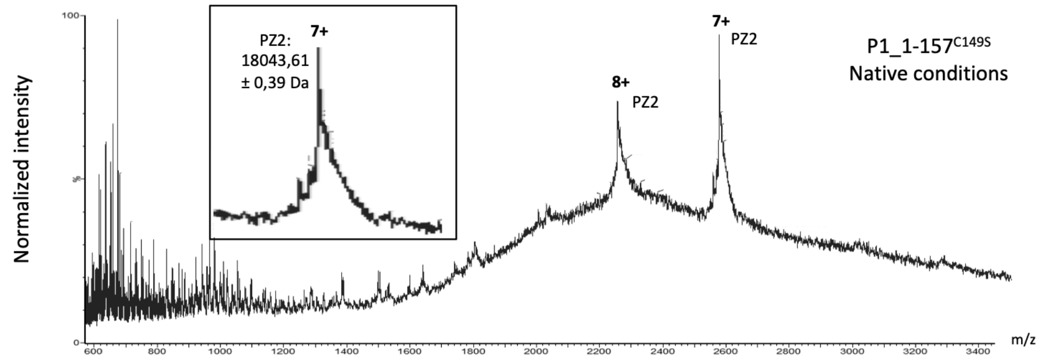

Poignavent *et al*, Appendix Figure S12

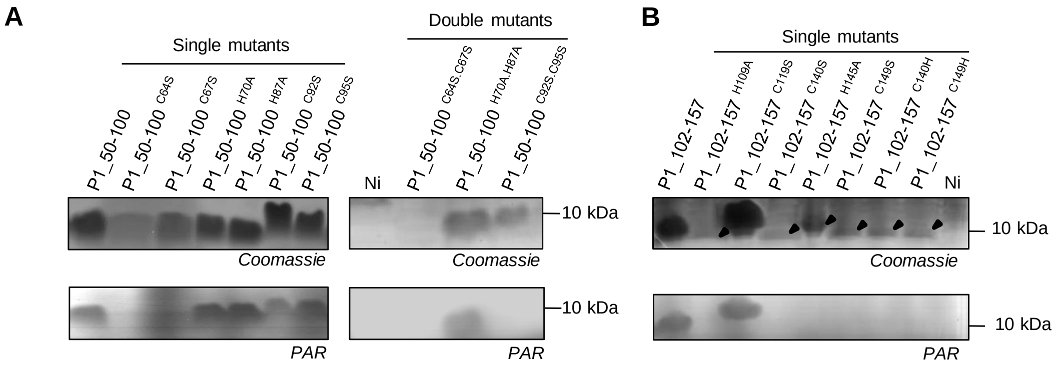

C

| P1_50-100 mutant proteins | Theoretical molecular mass | Molecular mass by denaturing ESI-MS | Molecular mass by native ESI-MS (major form PZ) | Difference between native ESI-MS and theoretical mass | Number of zinc atom (63 to 65 Da) |
|---|---|---|---|---|---|
| P1_50-100$^{H70A}$ | 5750.50 | 5750.5 ± 0.1 | 5813.6 ± 0.4 | 63,1 | 1 |
| P1_50-100$^{H87A}$ | 5750.50 | 5750.6 ± 0.1 | 5814.6 ± 0.9 | 64,1 | 1 |
| P1_50-100$^{H70A.H87A}$ | 5684.44 | 5684.7 ± 0.1 | 5748.2 ± 0.1 | 63,76 | 1 |
| P1_50-100$^{C64S}$ | 5800.51 | 5800.1 ± 0.1 | - | - | - |
| P1_50-100$^{C67S}$ | 5800.51 | 5800.1 ± 0.4 | - | - | - |
| P1_50-100$^{C64S.C67S}$ | 5784.45 | Not productible | - | - | - |
| P1_50-100$^{C92S}$ | 5800.51 | 5800.4 ± 0.2 | 5864.5 ± 0.1 | 63,99 | 1 |
| P1_50-100$^{C95S}$ | 5800.51 | 5800.7 ± 0.1 | 5863.9 ± 0.6 | 63,2 | 1 |
| P1_50-100$^{C92S.C95S}$ | 5784.45 | Not possible to purify | - | - | - |

Poignavent *et al*, Appendix Figure S13

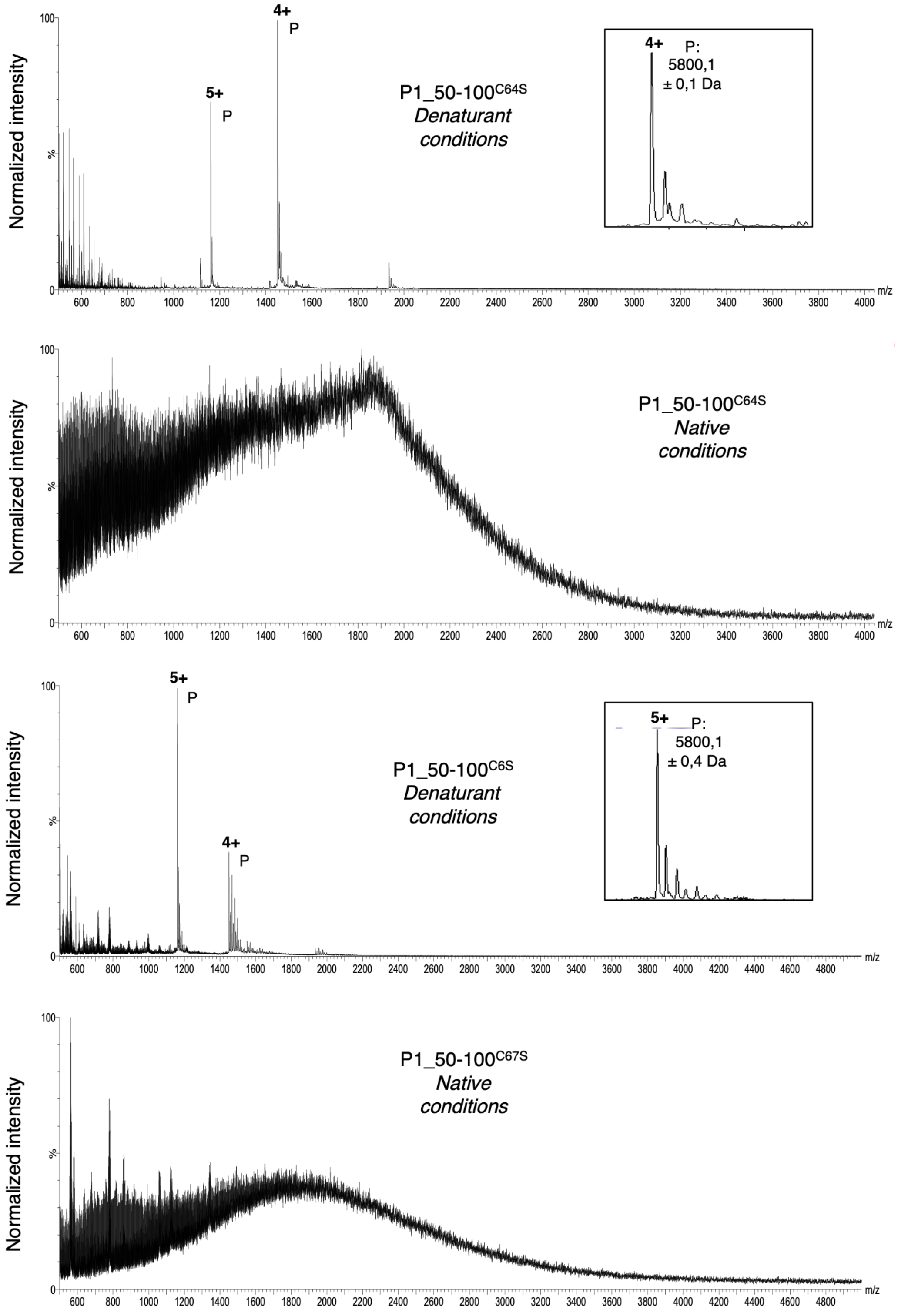

Poignavent *et al*, Appendix Figure S13

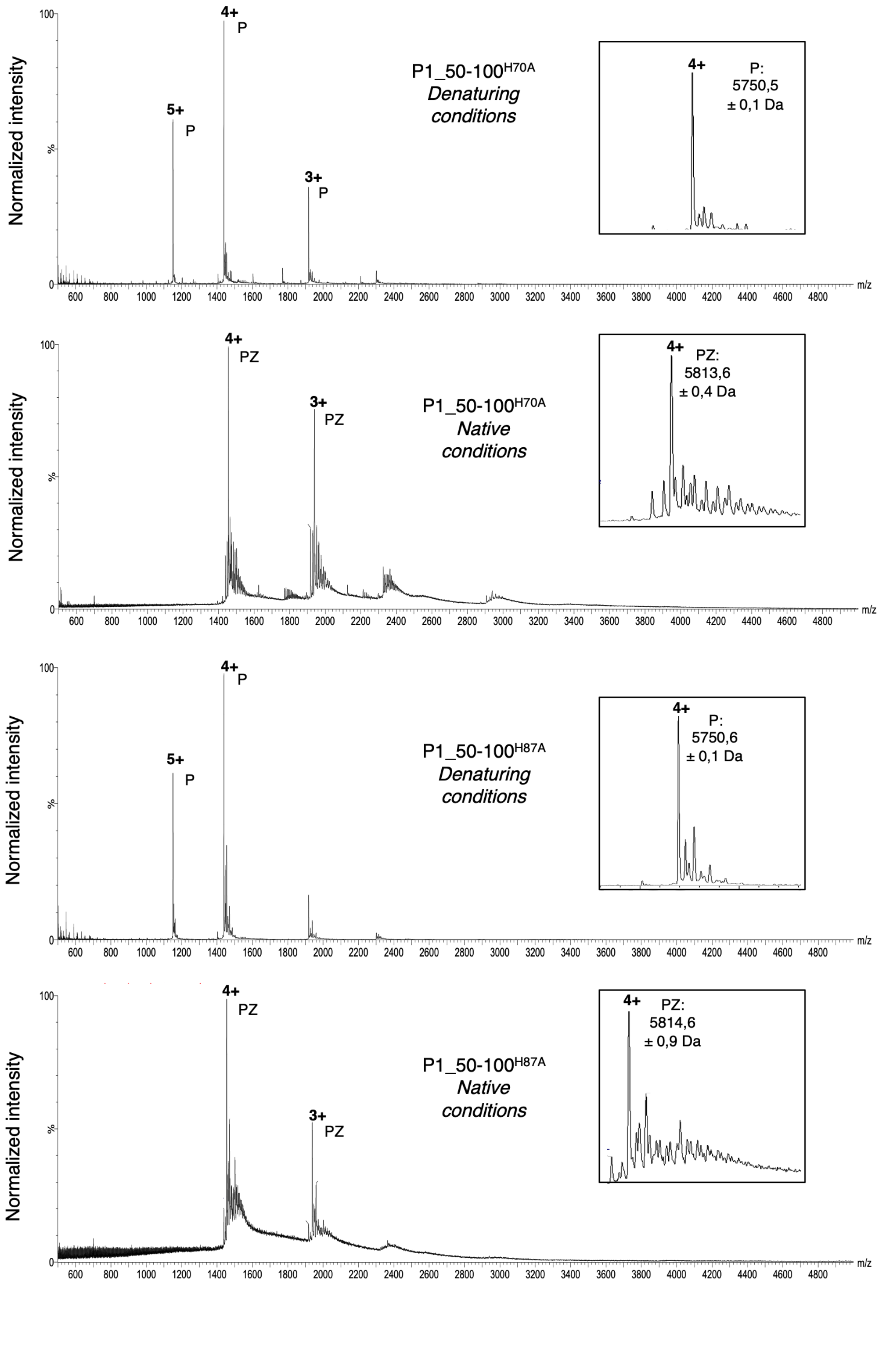

Poignavent *et al*, Appendix Figure S13

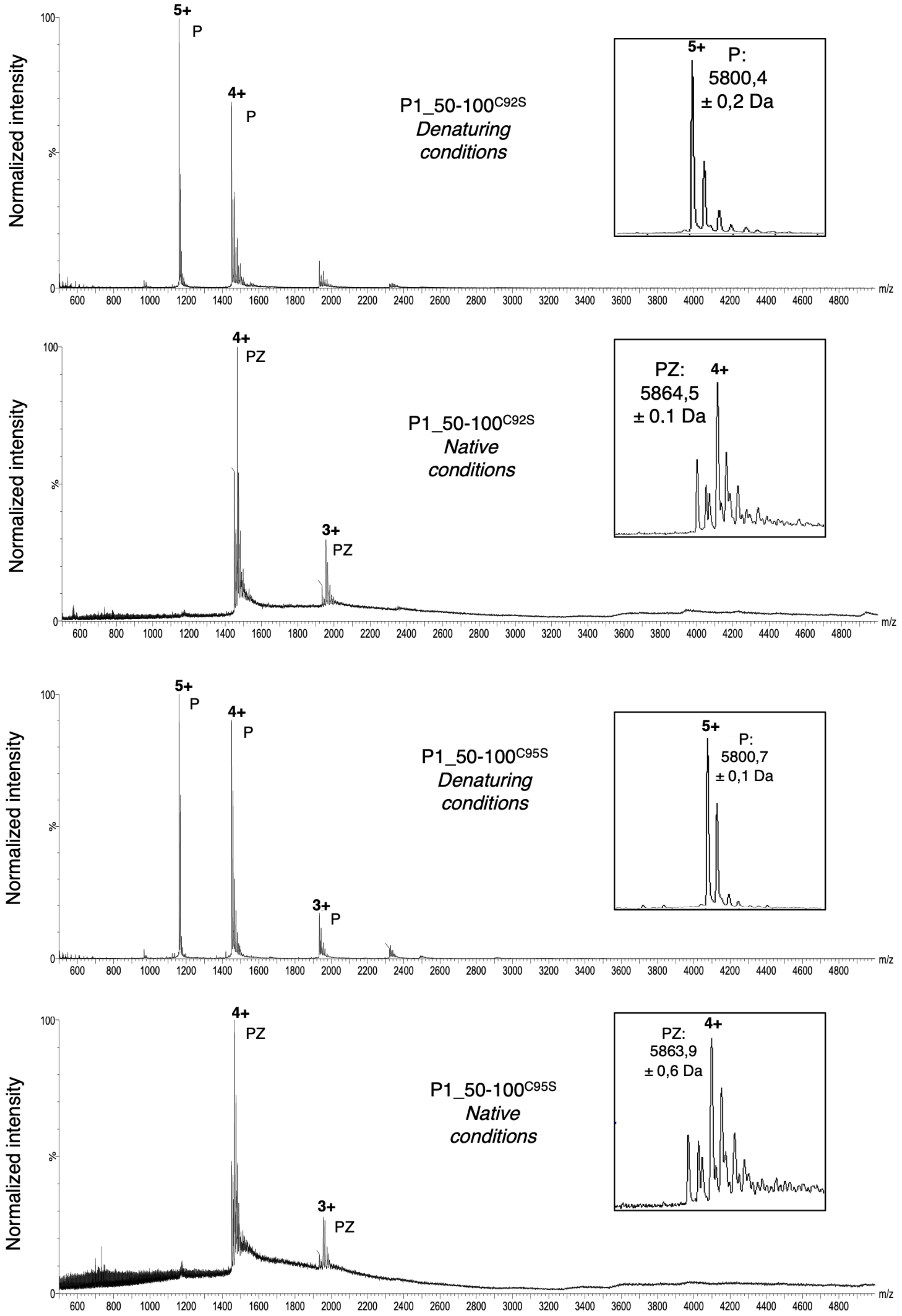

Poignavent *et al*, Appendix Figure S13

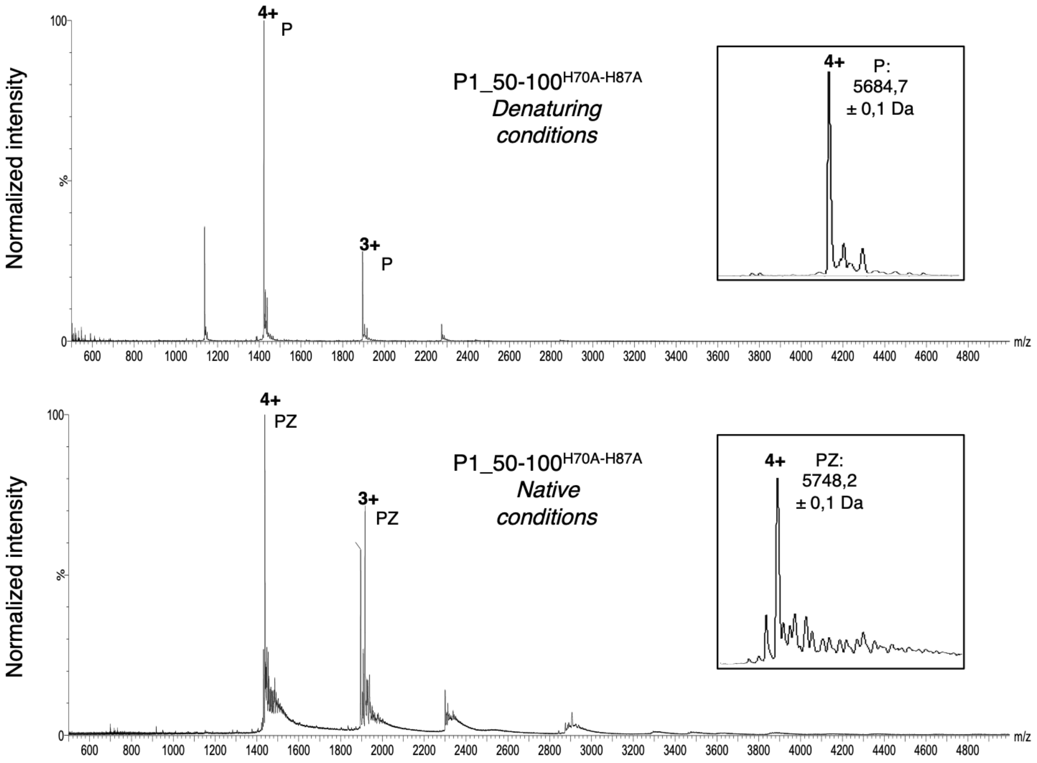

Poignavent *et al*, Appendix Figure S14

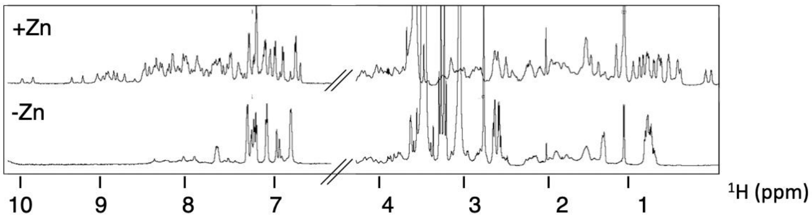

Poignavent *et al*, Appendix Table 3

| **Nb** | **residue** | **Phi (°)** | **Psi (°)** |
|---|---|---|---|
| 95 | CYS | -79 ± 39 | -59 ± 19 |
| 96 | GLU | -88 ± 48 | -55 ± 15 |
| 97 | ARG | -100 ± 36 | -60 ± 8 |
| 98 | SER | -100 ± 52 | -59 ± 1 |
| 99 | VAL | -141 ± 41 | -68 ± 52 |
| 101 | LEU | -83 ± 43 | -60 ± 16 |
| 102 | ASP | -104 ± 36 | -56 ± 16 |
| 103 | ASP | -84 ± 44 | -60 ± 20 |
| 104 | GLU | -85 ± 45 | -63 ± 23 |
| 105 | ILE | -83 ± 43 | 56 ± 16 |
| 106 | ASP | -82 ± 42 | 60 ± 20 |
| 107 | ARG | -82 ± 42 | 63 ± 23 |
| 108 | GLU | -86 ± 46 | 55 ± 15 |

Poignavent *et al*, Appendix Table 4

| Domain | $D_{\|\|}^{exp}$ ($10^{7}$ $s^{-1}$)[a] | $D_{\perp}^{exp}$($10^{-7}$ $s^{-1}$)[a] | $\tau_c^{exp}$ ($10^{-9}$ s) [b] | $D_{\|\|}^{calc}$($10^{7}$ $s^{-1}$)[c] | $D_{\perp}^{calc}$($10^{-7}$ $s^{-1}$)[c] | $\tau_c^{calc}$($10^{-9}$ s) [d] |
|---|---|---|---|---|---|---|
| P1_1-100 | 1.55 ± 0.06 | 0.91 ± 0.07 | 14.5 ± 0.2 | | | |
| P1_102-157 | 1.84 ± 0.07 | 1.12 ± 0.02 | 12.9 ± 0.2 | | | |
| Full length P1 | | | | 1.7 | 0.4 | 19.9 |

[a]:Diffusion tensor components were estimated by TENSOR2 (Dosset *et al*, 2000) from $^{15}$N relaxation data recorded on full-length P1_1-157 but taking the data for each domain (P1_1-100 and P1_102-157) separately. Estimated components are smaller for the N-terminal domain than for the C-terminal domain, reflecting the slower tumbling time of the former $\tau_c^{exp}$ in the full-length protein.

[b]: Overall global estimated correlation time computed from the components of the rotational diffusion tensor estimated from the relaxation times.

[c]: Diffusion tensor components were computed from the NMR derived structure of P1_1-157 by HYDRONMR (García de la Torre *et al*, 2000). The calculated components are smaller than the estimated components from experimental data, as is expected for a two-domain protein exhibiting inter-domain flexibility.

[b]: Overall global estimated correlation time calculated from the NMR-derived structure.

Poignavent *et al*, Appendix Figure S15

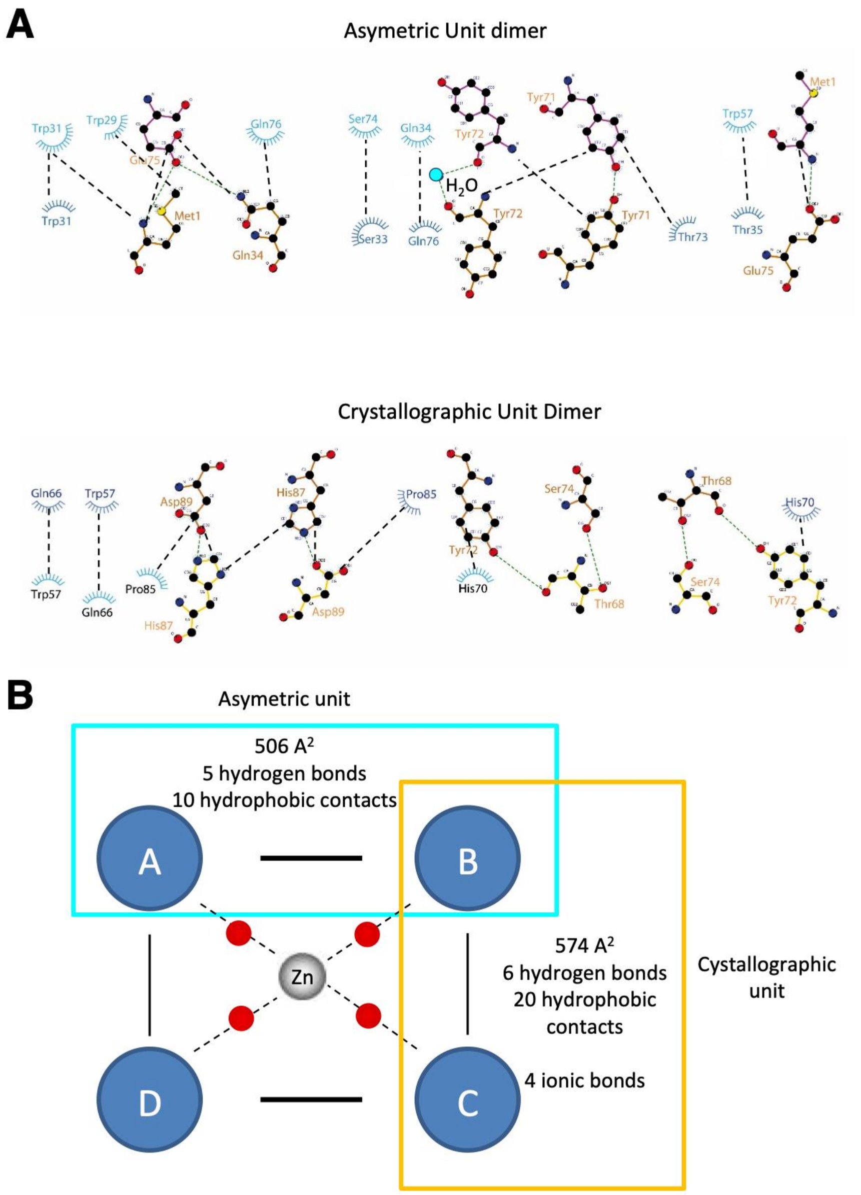

Poignavent *et al.*, Appendix Table 5

| **Interfacing structures** | **Buried area, Å2** | **$\Delta^{i}G$, kcal/mol**[a] | **NHB**[b] | **NSB**[c] |
|---|---|---|---|---|
| B + D (cryst.) | 584.6 (9.9%) | -6.3 (9%) | 6 (30%) | 4 (25%) |
| A + C | 565.1 (9.3%) | -8.6 (14%) | 4 (20%) | 4 (25%) |
| Average: | 574.1 (9.6%) | -7.4 (11%) | 5 (25%) | 4 (25%) |
| A + B (asym.) | 506.2 (8.6%) | -5.7 (9%) | 5 (25%) | 4 (25%) |
| C + D | 506.2 (8.6%) | -5.7 (9%) | 5 (25%) | 4 (25%) |
| Average: | 506.2 (8.6%) | -5.7 (9%) | 5 (25%) | 4 (25%) |
| B + C | 111.6 (1.9%) | -0.3 (0%) | 0 (0%) | 0 (0%) |
| A + D | 111.6 (1.9%) | -0.3 (0%) | 0 (0%) | 0 (0%) |
| Average: | 111.6 (1.9%) | -0.3 (0%) | 0 (0%) | 0 (0%) |
| D + [ZN]C:3 | 27.8 (1%) | -10.3 (16%) | 0 (0%) | 0 (0%) |
| B + [ZN]C:3 | 27.5 (1%) | -10.2 (16%) | 0 (0%) | 0 (0%) |
| A + [ZN]C:3 | 24.8 (0%) | -8.9 (14%) | 0 (0%) | 0 (0%) |
| C + [ZN]C:3 | 24.7 (0%) | -8.9 (14%) | 0 (0%) | 0 (0%) |
| Average: | 26.2 (1%) | -9.6 (15%) | 0 (0%) | 0 (0%) |

[a]:Negative $\boldsymbol{\Delta^{i}G}$ corresponds to hydrophobic interfaces, or to positive protein affinity. This value does not include the effect of satisfied hydrogen bonds and salt bridges across the interface.

[b]:number of potential hydrogen bonds across the interface

[c]:number of potential salt bridges across the interface

Poignavent *et al*, Appendix Figure S16

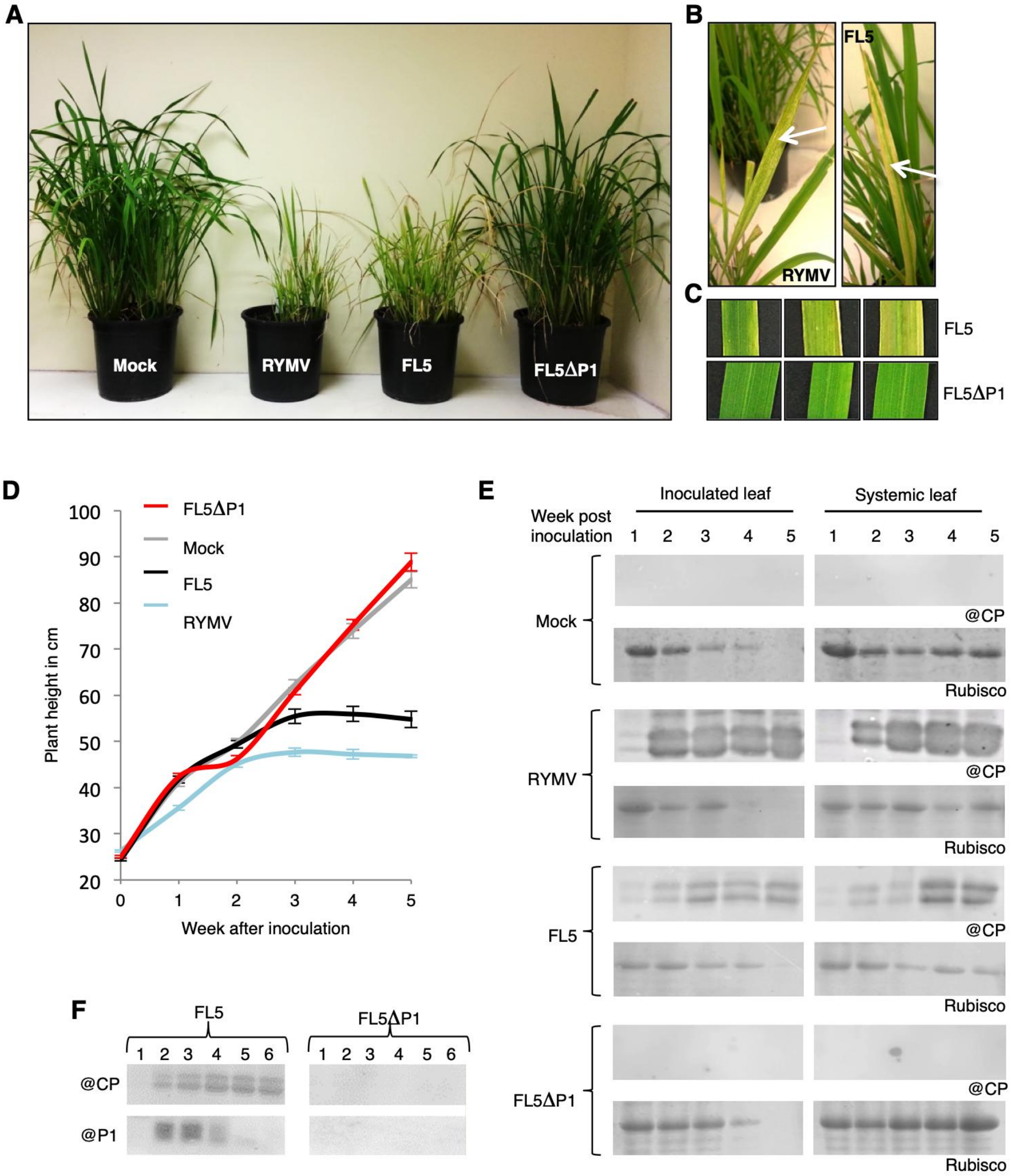

Poignavent *et al*, Appendix Figure S17

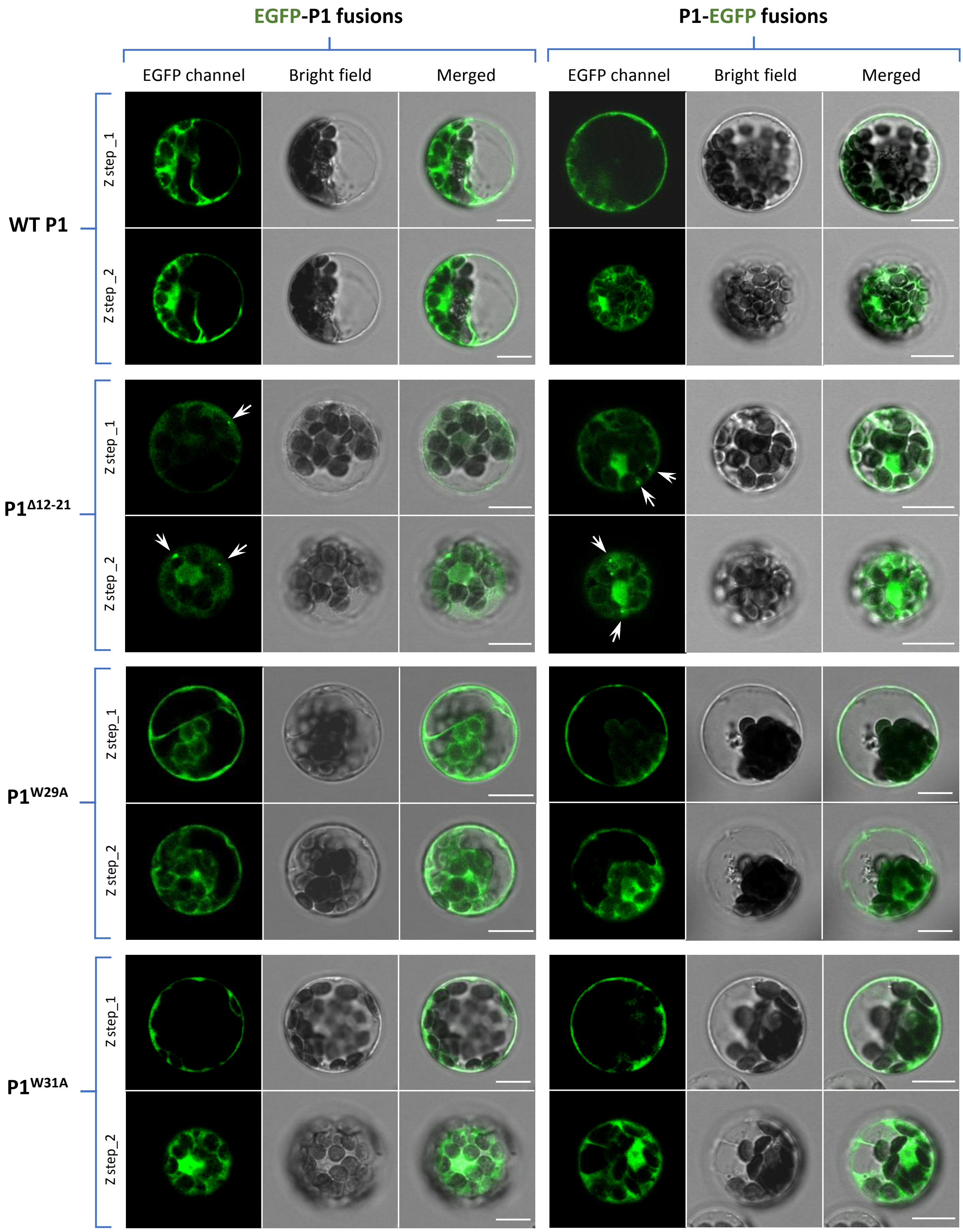

Poignavent *et al*, Appendix Figure S17

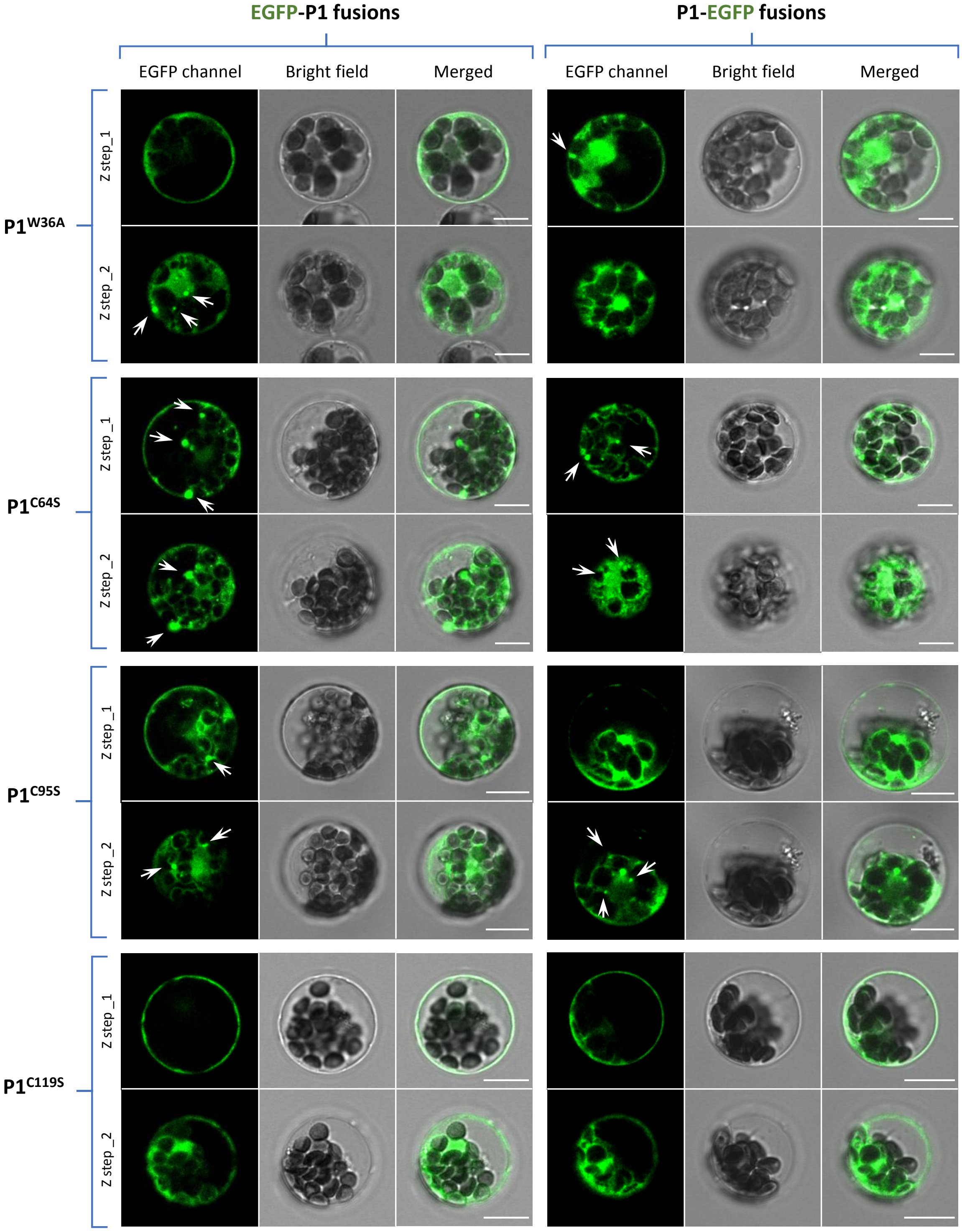

Poignavent *et al*, Appendix Figure S17

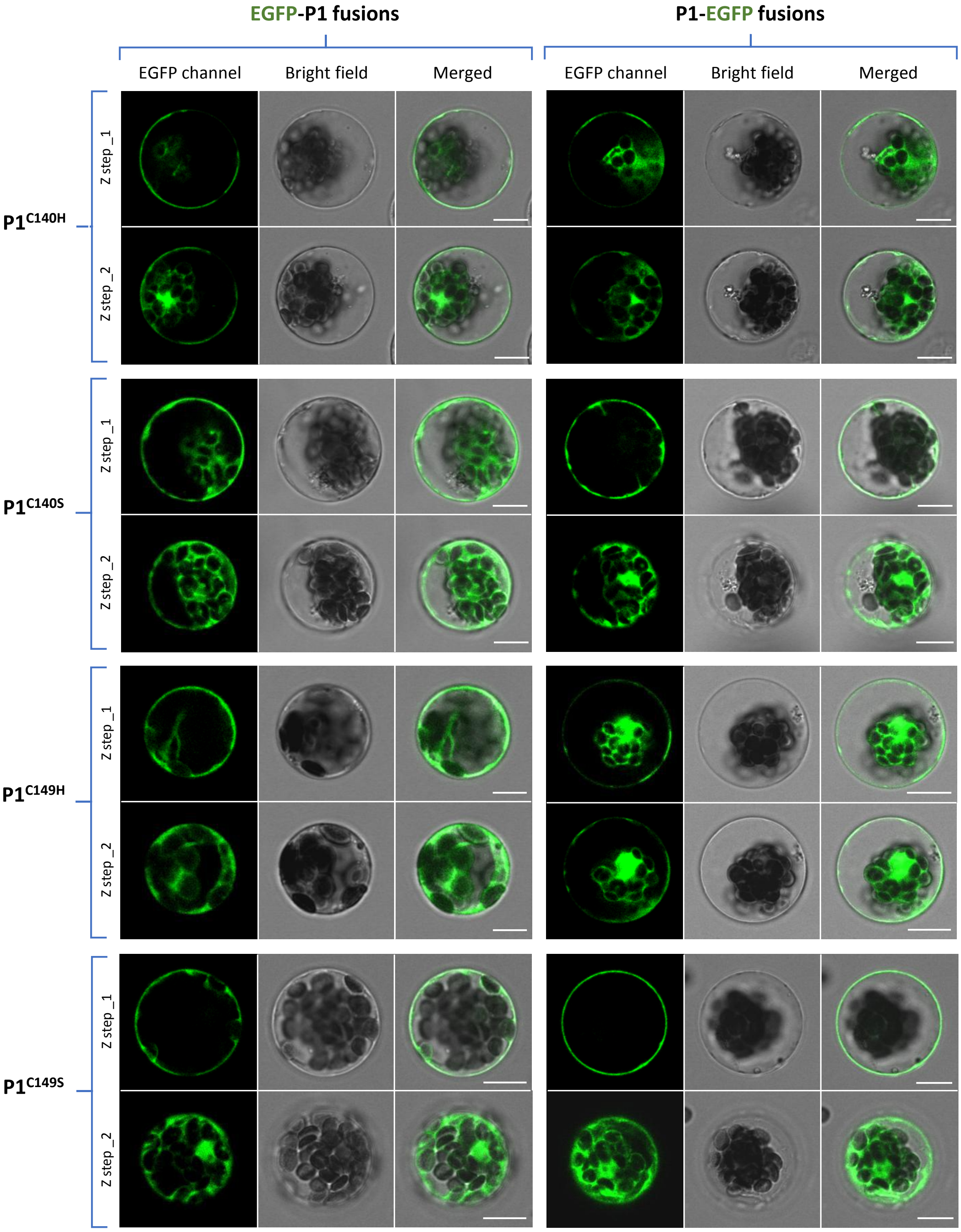

Poignavent *et al*, Appendix Figure S17

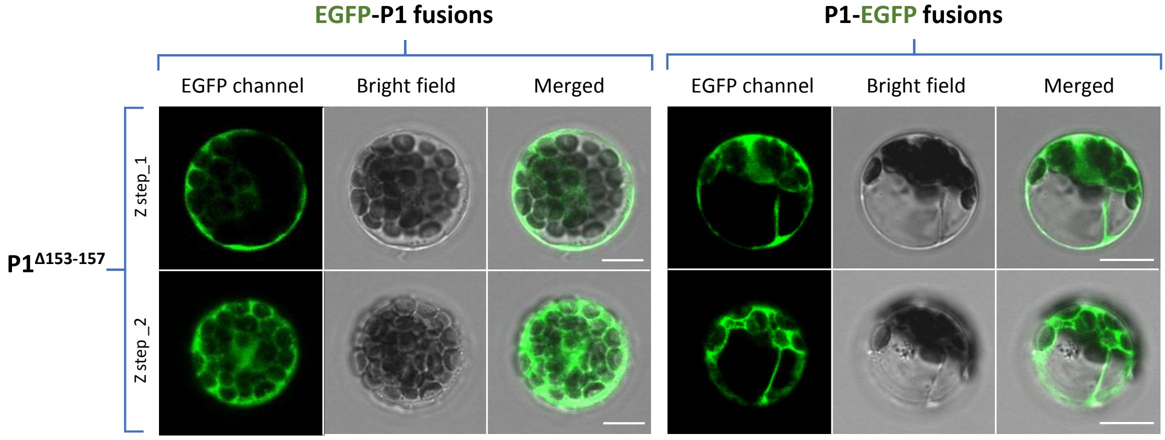

Poignavent *et al*, Appendix Figure S18

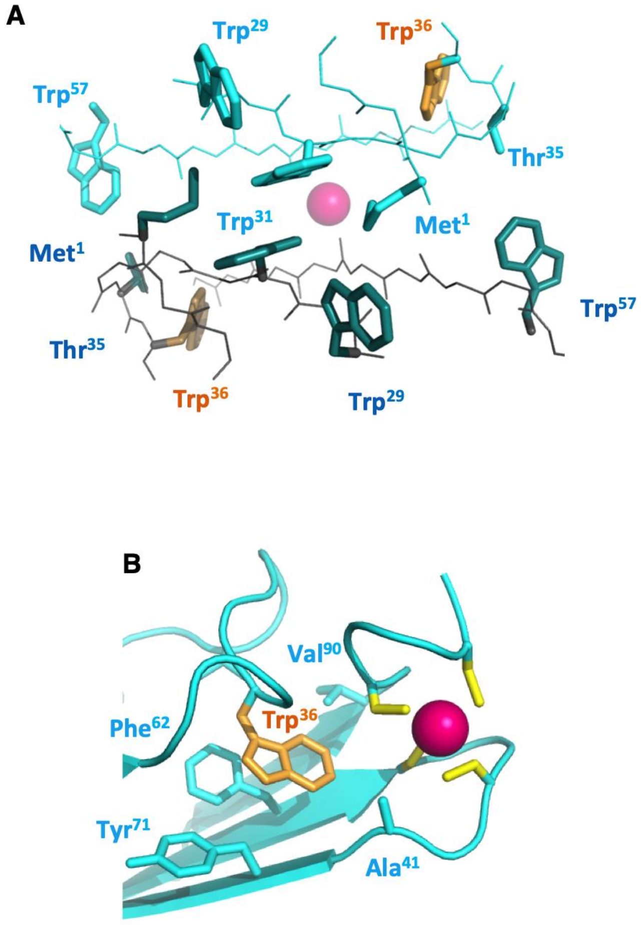

Poignavent *et al*, Appendix Figure S19

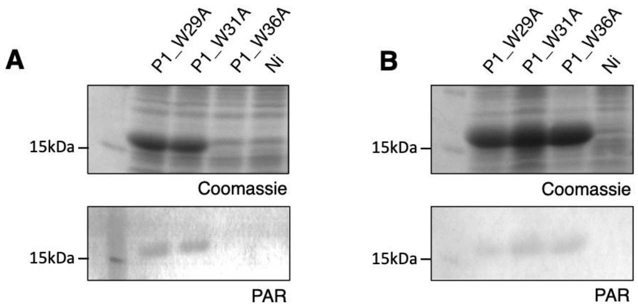

Poignavent *et al*, Appendix Figure S20

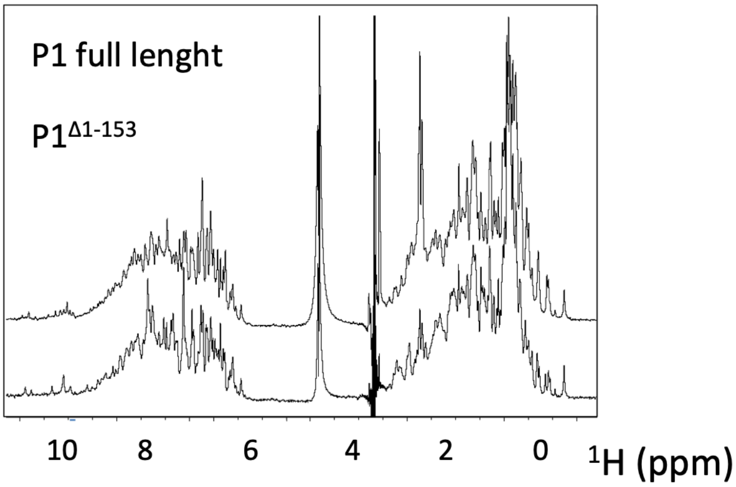
